# Large-scale bidirectional arrayed genetic screens identify *OXR1* and *EMC4* as modifiers of α-synuclein aggregation

**DOI:** 10.1101/2025.06.10.658866

**Authors:** Sandesh Neupane, Lea Nikolić, Lorenzo Maraio, Thomas Goiran, Nathan Karpilovsky, Stefano Sellitto, Vangelis Bouris, Jiang-An Yin, Ronald Melki, Edward A. Fon, Elena De Cecco, Adriano Aguzzi

**Affiliations:** Institute of Neuropathology, University of Zurich, 8091 Zurich, Switzerland; Laboratory of Prion Biology, Department of Neuroscience, Scuola Internazionale Superiore di Studi Avanzati (SISSA), Trieste, Italy; Neurodegenerative Diseases Research Group, Department of Neurology and Neurosurgery, Montreal Neurological Institute-Hospital, McGill University, Montreal, Quebec H3A 2B4, Canada; Institut François Jacob (MIRCen) and CNRS, Fontenay-aux-Roses, France

## Abstract

In Parkinson’s disease and other synucleinopathies, α-synuclein (α-Syn) misfolds and forms Ser^129^-phosphorylated aggregates (pSyn^129^). The factors controlling this process are largely unknown. Here, we used arrayed CRISPR-mediated gene activation and ablation to discover new pSyn^129^ modulators. Using quadruple-guide RNAs (qgRNAs) and Cas9, or an inactive Cas9 version fused to a synthetic transactivator, we ablated 2304 and activated 2428 human genes related to mitochondrial, trafficking and motility function in HEK293 cells. After exposure of cells to α-Syn fibrils, pSyn^129^ signals were recorded by high-throughput fluorescence microscopy and aggregates were identified by image analysis. We found that pSyn^129^ was increased by activating the mitochondrial protein *OXR1,* which decreased ATP levels and altered the mitochondrial membrane potential. Instead, pSyn^129^ was reduced by ablation of the endoplasmic reticulum (ER)-associated protein *EMC4,* which enhanced ER-driven autophagic flux and lysosomal clearance. OXR1 activation preferentially modulated cellular reactions to fibrils derived from multiple system atrophy (MSA) patients, whereas EMC4 ablation broadly reduced pSyn^129^ across diverse α-Syn polymorphs. These findings were confirmed in human iPSC-derived cortical and dopaminergic neurons, where OXR1 preferentially promoted somatic aggregation and EMC4 reduced both somatic and neuritic aggregates. These results uncover previously unrecognized roles for OXR1 and EMC4 in α-Syn aggregation, thereby broadening our mechanistic understanding of synucleinopathies.

## Introduction

Parkinson’s disease (PD) is the second most prevalent neurodegenerative disorder, affecting over 10 million individuals worldwide, with its incidence steadily rising due to the ageing population (Poewe *et al*, 2017; Su *et al*, 2025). Clinically, PD is characterized by progressive motor impairments (bradykinesia, rigidity, resting tremor) and a range of non-motor symptoms frequently including cognitive decline (Schapira *et al*, 2017; Stocchi *et al*, 2024). PD belongs to a broader class of disorders collectively termed synucleinopathies, which includes multiple system atrophy (MSA) and dementia with Lewy bodies (DLB) (Park *et al*, 2025; Simuni *et al*, 2024). A unifying histopathological hallmark of these conditions is the accumulation of misfolded α-synuclein (α-Syn) aggregates within neuronal and glial cells, giving rise to the formation of Lewy bodies, Lewy neurites, and glial cytoplasmic inclusions (Spillantini *et al*, 1997).

α-Syn, encoded by the *SNCA* gene, is a small presynaptic protein that modulates synaptic vesicle trafficking and neurotransmitter release (Lashuel *et al*, 2013). Under pathological conditions, α-Syn aggregates into oligomers and protofibrils (Tofaris & Spillantini, 2007; Trojanowski & Lee, 1998), eventually forming amyloid inclusions through nucleated polymerization in a prion-like fashion (Aguzzi & Rajendran, 2009; Mahul-Mellier *et al*, 2020; Neupane *et al*, 2023; Scheckel & Aguzzi, 2018). More than 90% of α-Syn aggregates in postmortem brains from individuals with synucleinopathies are phosphorylated at Ser^129^ (pSyn^129^) (Ramalingam & Dettmer, 2023). The functional role of phosphorylation in disease progression remains debated, with studies proposing both pro-aggregatory (Ma *et al*, 2016) and neuroprotective effects (Ghanem *et al*, 2022; Kontaxi & Edwards, 2023). Nevertheless, its consistent association with pathological α-Syn inclusions has led to its widespread adoption as a surrogate marker of α-Syn aggregation in cellular and animal models (Parra-Rivas *et al*, 2023). These aggregates disrupt proteostasis, impair endoplasmic reticulum (ER)-Golgi trafficking and mitochondrial function, and induce oxidative and nitrosative stress, ultimately leading to synaptic dysfunction and neuronal death (Lv *et al*, 2019; Mehra *et al*, 2019; Stykel & Ryan, 2022).

Genetic and biochemical evidence implicates the dysfunction of mitochondria and lysosomes in idiopathic and genetic forms of PD (Chen *et al*, 2023; Ganjam *et al*, 2019; Geibl *et al*, 2024). 85–90% of PD cases are sporadic, with aging and exposure to environmental toxins as major risk factors (Pang *et al*, 2019; Paul *et al*, 2023), whereas the remaining 10–15% are clustered in familial patterns. Studies of familial PD genes and genome-wide association studies (GWAS) have highlighted polymorphisms and mutations of lysosomal/endosomal genes (*GBA1, LRRK2, TMEM175*, and *VPS35*) and mitochondrial quality-control genes (*PINK1*, *PRKN*, *DJ-1*), linking dysfunction in these organelles to α-Syn aggregation and neurodegeneration (Brooker *et al*, 2024). Accumulation of misfolded proteins induces ER stress and subsequent activation of stress-response pathways can contribute to neurodegenerative progression (Mnich *et al*, 2023; Mou *et al*, 2020). These findings suggest that dissecting the causal links between organelle dysfunction and cellular pathology will enhance our mechanistic understanding of synucleinopathies.

Functional genomics provides a powerful toolbox to identify cellular modifiers of α-Syn aggregation and propagation, and indeed siRNA and shRNA-based screens have identified candidate regulators (Bieri *et al*, 2019; Gonçalves *et al*, 2016; Kara *et al*, 2021). However, these approaches are limited by off-target effects, inefficient suppression of target genes, and low specificity, which can confound the identification of crucial modifiers. Pooled CRISPR screens offer improved specificity and have provided meaningful insights into α-Syn pathology (Chen *et al*, 2017; Hu *et al*, 2023; Santhosh Kumar *et al*, 2024; See *et al*, 2021; Vanderperre *et al*, 2023), but are not well suited for scoring morphological phenotypes. In contrast, arrayed CRISPR screens enable high-content phenotyping of individual gene perturbations, combining CRISPR activation (CRISPRa) and ablation (CRISPRo) to systematically map bidirectional regulatory networks (Aguzzi & Kampmann, 2023; Yin *et al*, 2024).

Here, we developed a dual CRISPRa/CRISPRo platform to target mitochondrial, trafficking, and motility (MTM) genes in human cellular models. We performed an image-based arrayed CRISPR screen to quantify pSyn^129+^ aggregates using custom libraries (T.gonfio for CRISPRa; T.spiezzo for CRISPRo), and key hits were validated in pathogenetically relevant human iPSC-derived cortical and dopaminergic neurons. The genes identified in this study provide insights into the organelle-specific genetic nodes of α-Syn pathology.

## Results

### A high-content CRISPR screening platform to identify regulators of α-Syn aggregation

To establish a microscopy-based CRISPR screen for genetic regulators of α-Syn aggregation, we used HEK293 cells stably overexpressing human wild-type (wt) α-Syn (henceforth named HEK^Syn^). Despite being immortalized, HEK293 cells share some biochemical features with neurons (Shaw *et al*, 2002) and provide the scalability and susceptibility to genetic manipulation necessary for arrayed high-throughput screens. Upon addition of exogenous α-Syn pre-formed fibrils (PFFs), HEK^Syn^ cells form intracellular pSyn^129^ aggregates that resemble Lewy body-like inclusions (Luk *et al*, 2009).

We generated HEK^Syn^ clones expressing Cas9 or the tripartite programmable transactivator dCas9-VPR (Chavez *et al*, 2015) for CRISPR ablation (CRISPRo) and CRISPR activation (CRISPRa), respectively (**Figure 1A**). We isolated and expanded five single-cell clones for each of the CRISPRo and CRISPRa lines. One clone per condition was then selected after evaluating (i) stable expression of Cas9 or dCas9-VPR, (ii) stable overexpression of α-Syn, and (iii) the ability to form intracellular α-Syn aggregates upon exposure to α-Syn PFFs. These clones retained the CRISPR machinery, α-Syn expression, and generated pSyn^129^ aggregates after exposure to PFF, confirming their suitability for all subsequent experiments. To modulate gene expression, we used the T.gonfio and T.spiezzo arrayed libraries (Yin *et al*, 2024) which target each gene with four non-overlapping single guide RNAs (qgRNA). We then optimized transfection conditions for high qgRNA delivery efficiency in 384-well format (**Fig. S1A-S1B**). CRISPRa lines showed robust transcriptional upregulation, with mRNA expression levels increasing up to ∼10,000-fold (**Figure 1B),** and Western blotting confirmed elevated expression of a representative target protein **(Figure S1C**). Similarly, CRISPR-based gene ablation in CRISPRo lines with randomly selected qgRNAs resulted in a ∼95% reduction in target gene expression within 5 days after transfection (**Figures 1C and S1D–S1F**). Next, we generated α-Syn PFFs from purified monomeric human wt α-Syn and confirmed their amyloid fibrillar structure by negative-stain transmission electron microscopy (TEM), SDS-PAGE, and Thioflavin T (ThT) fluorescence assays (**Figures 1D and S1G-S1H**). Since smaller fibrils exhibit potent seeding capability (Emin *et al*, 2022), we sonicated the PFFs preparation to obtain a heterogeneous population of fibrillar fragments (average length: 30-100 nm; **Figure 1E**). We then treated HEK^Syn^ cells with sonicated PFFs mixed with the transfection reagent LT1 to facilitate efficient intracellular delivery through lipid-based complex formation. As a proxy for α-Syn aggregation, we quantified pSyn^129+^ inclusions (Anderson *et al*, 2006; Trist *et al*, 2024). Confocal imaging confirmed the presence of pSyn^129^ within the cell body exclusively in PFFs-treated cells (**Figure S1I**).

**Figure 1:**
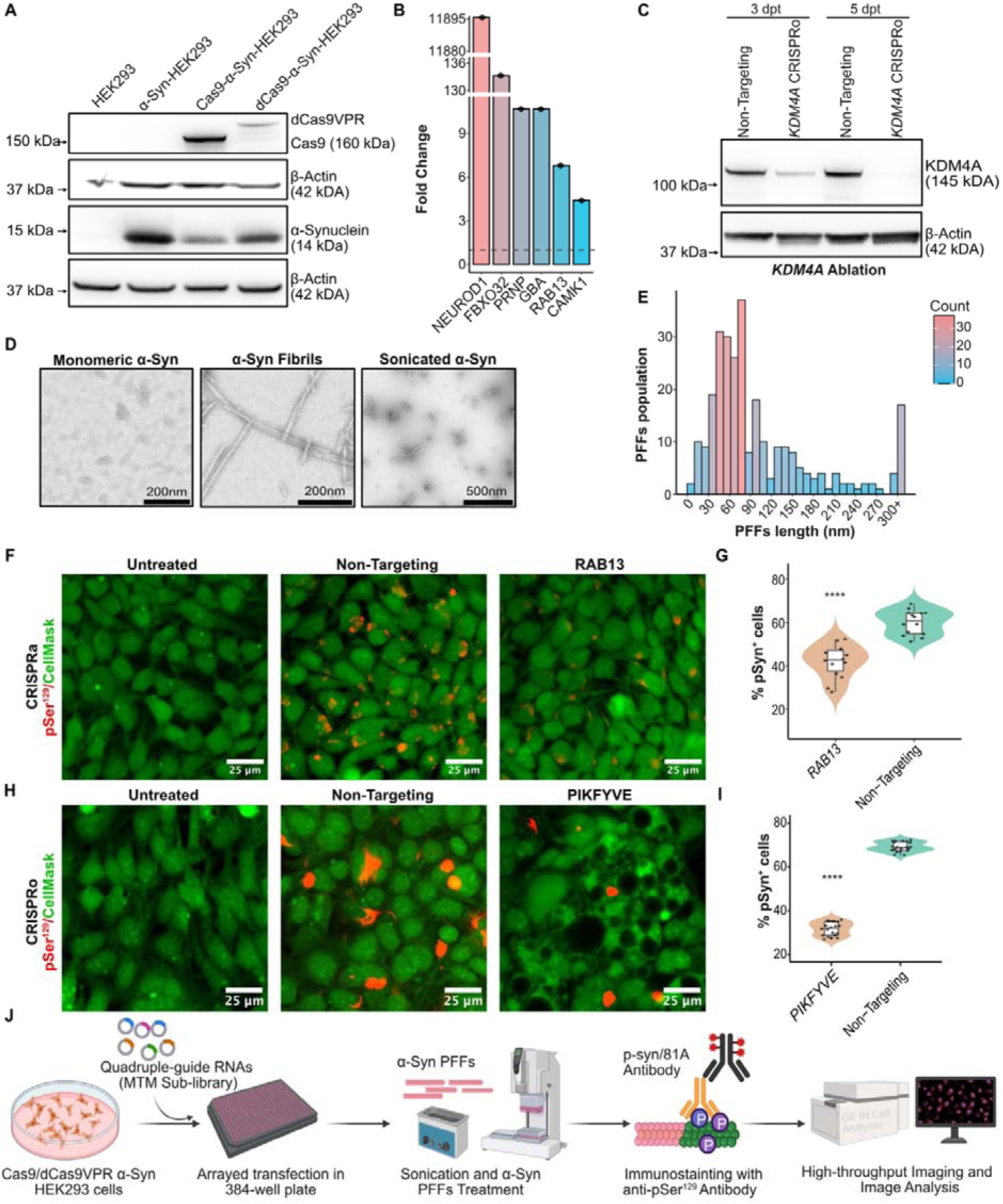
CRISPR screening workflow and α-Synuclein (α-Syn) phosphorylation assay in cellular model. **(A)** Western blot analysis of HEK293 clones overexpressing α-Syn (α-Syn-HEK293/HEK^Syn^) with either dCas9-VPR (CRISPRa) or Cas9 (CRISPRo). (B) RT-qPCR of arbitrarily selected genes for the assessment of dCas9-VPR activity normalized to non-targeting controls. (C) Gene ablation efficiency/Cas9 activity on the cells measured at 3- and 5-days post-transfection (dpt). **(D)** Representative negative stain TEM micrographs of *in vitro* produced α-Syn PFFs. **(E)** Distribution of quantified sonicated PFFs lengths. **(F, H)** Representative immunofluorescence of HEK293 cells showing pSyn aggregates (red) and cytosol (green). **(G, I)** Quantification of p-Ser129^+^ cells normalized to total DAPI count. Inner box plots: median (center line), 75th and 25^th^ percentile (top and bottom edge). **(J)** Schematic representation of the high-content arrayed CRISPR screening assay for phosphorylated α-Synuclein at Ser129 (pSyn) detected using antibody 81A. Welch’s t-test (unequal variance t-test, *P < 0.05, **P < 0.01, ***P < 0.001, ****P < 0.0001).

The absence of cytoplasmic pSyn^129^ signal in HEK293 cells lacking α-Syn overexpression demonstrates that elevated α-Syn levels are essential to drive robust and rapid aggregation. Moreover, it indicates that the 81A antibody selectively recognizes de novo aggregates rather than the recombinant seeds (**Figure S1J**). Collectively, these findings lay the foundation for a platform enabling the systematic interrogation of genetic regulators of α-Syn aggregation.

To monitor the quality and reliability of the screen, we incorporated *RAB13* and *PIKFYVE* as positive control genes, which have been previously reported to modulate pSyn^129^ levels. For the CRISPRa screen, we used *RAB13* which has been shown to reduce α-Syn aggregates upon overexpression (Gonçalves *et al*, 2016). For the CRISPRo screen, we used *PIKFYVE*, whose inhibition (See *et al*, 2021) decreases α-Syn aggregation. High-throughput fluorescence imaging revealed a significant reduction in percentage of pSyn^129+^ cells with activated *RAB13* (**Figures 1F-1G**). Likewise, ablation of *PIKFYVE* resulted in cytoplasmic vacuolation and a decrease in the percentage of pSyn^129+^ cells (**Figures 1H-1I**). These results validate the robustness of our screening platform and establish stringent quality-control metrics for identifying genetic modulators of pSyn^129^.

### Arrayed CRISPRa and CRISPRo screens identify modulators of pSyn^129^

Mitochondrial dysfunction is a key contributor to PD pathology (Thorne & Tumbarello, 2022). Recent reports suggest that the inner and outer mitochondrial membranes act as primary sites for early aggregation events (Choi *et al*, 2022). Hence, we conducted CRISPRa and CRISPRo screens targeting genes involved in mitochondrial homeostasis and function, intracellular trafficking and cytoskeletal reorganization. HEK^Syn^ cells expressing Cas9 or dCas9-VPR were transfected with 2304 or 2428 individual qgRNA plasmids, respectively, as duplicates in separate 384-well plates, selected with puromycin for 48 hours (CRISPRa) or 96 hours (CRISPRo). Cells were then treated for 72 hours with sonicated PFFs delivered via lipofection using the LT1 reagent (**Figure 1J**). Cell nuclei were stained with DAPI, pSyn^129^ aggregates were labeled using the 81A antibody, and the entire cell was uniformly labeled with HCS CellMask™ Deep Red. Images were acquired using automated widefield imaging systems and analyzed by semi-automated machine learning-based pixel classification (ilastik) in combination with object segmentation (CellProfiler) (**Figure S2A**). We created two ilastik probability maps: one for the DAPI-stained nuclei and one for the pSyn^129^ aggregates. These maps were then imported into CellProfiler, where the nuclear map served as a reference to segment cells based on the CellMask channel. The pSyn^129^-aggregate map was overlaid with the segmented cells to determine the number of cells positive for at least one aggregate. We defined the fraction of pSyn^129+^ cells as the number of cells containing at least one aggregate divided by the total number of segmented cells.

The robustness of the primary screens was quantitated by calculating Strictly Standardised Mean Difference (SSMD) scores based on non-targeting qgRNAs (NTG) and moderate-strength positive controls. CRISPRa and CRISPRo screens achieved SSMD values above 1 and 2, respectively (**Figures S2B-S2E**). These metrics indicate a good quality of the screen (Bray & Carpenter, 2017). Next, we evaluated reproducibility between replicates using correlation analyses suited to each dataset’s distribution. Since the CRISPRa data followed a normal distribution, we applied Pearson’s correlation coefficient (r = 0.75, adjusted R² = 0.59; **Figure S2C**). In contrast, the CRISPRo data deviated from normality, so we used Spearman’s rank correlation coefficient (ρ = 0.856; **Figure S2F**). Replicates from both screens correlated strongly, demonstrating technical reproducibility. To visualize the frequency distribution of individual samples for all targeted genes, we plotted histograms of the median values of pSyn^129^⁺ cell fraction. Most genes showed no significant impact on pSyn^129^⁺ levels (**Figures S2D, S2G**). Primary hitlists were compiled by selecting all genes with a mean log2-transformed fold change (log2FC) > 0.58 or < -0.58 for up- or downregulators, and statistical significance (moderated t-test p < 0.01; **Table S1:** CRISPRa**; Table S2;** CRISPRo). The cutoff values for the log2FC compensate for the overall baseline variability among the NTG controls. Moreover, all samples with cell counts lower than half the median of the NTG control were discarded. Both CRISPR screens identified multiple genetic modifiers of pathological pSyn^129^, using the percentage of pSyn^129+^ cells as a readout of aggregated pSyn^129^ levels (**Figures 2A-2B**). Consistent with previous findings, *RAB13* overexpression (CRISPRa screen) reduced pSyn^129^⁺ prevalence, whereas *RAB13* ablation (CRISPRo screen) enhanced pSyn^129^⁺ levels (**Figures 2D-2E**) (Gonçalves *et al*, 2016), further validating the robustness of our assay. To assess the bidirectional regulatory effects of the identified hit genes on pSyn^129^⁺ prevalence, we compared log2FC scores between CRISPRa and CRISPRo screens (**Figure 2C**). Genes with opposing effects clustered in distinct quadrants, with CRISPRa upregulators acting as downregulators in CRISPRo, and vice versa. However, bidirectional effect sizes were modest, suggesting that while such regulation exists, its impact on pSyn^129^⁺ prevalence may be limited.

**Figure 2:**
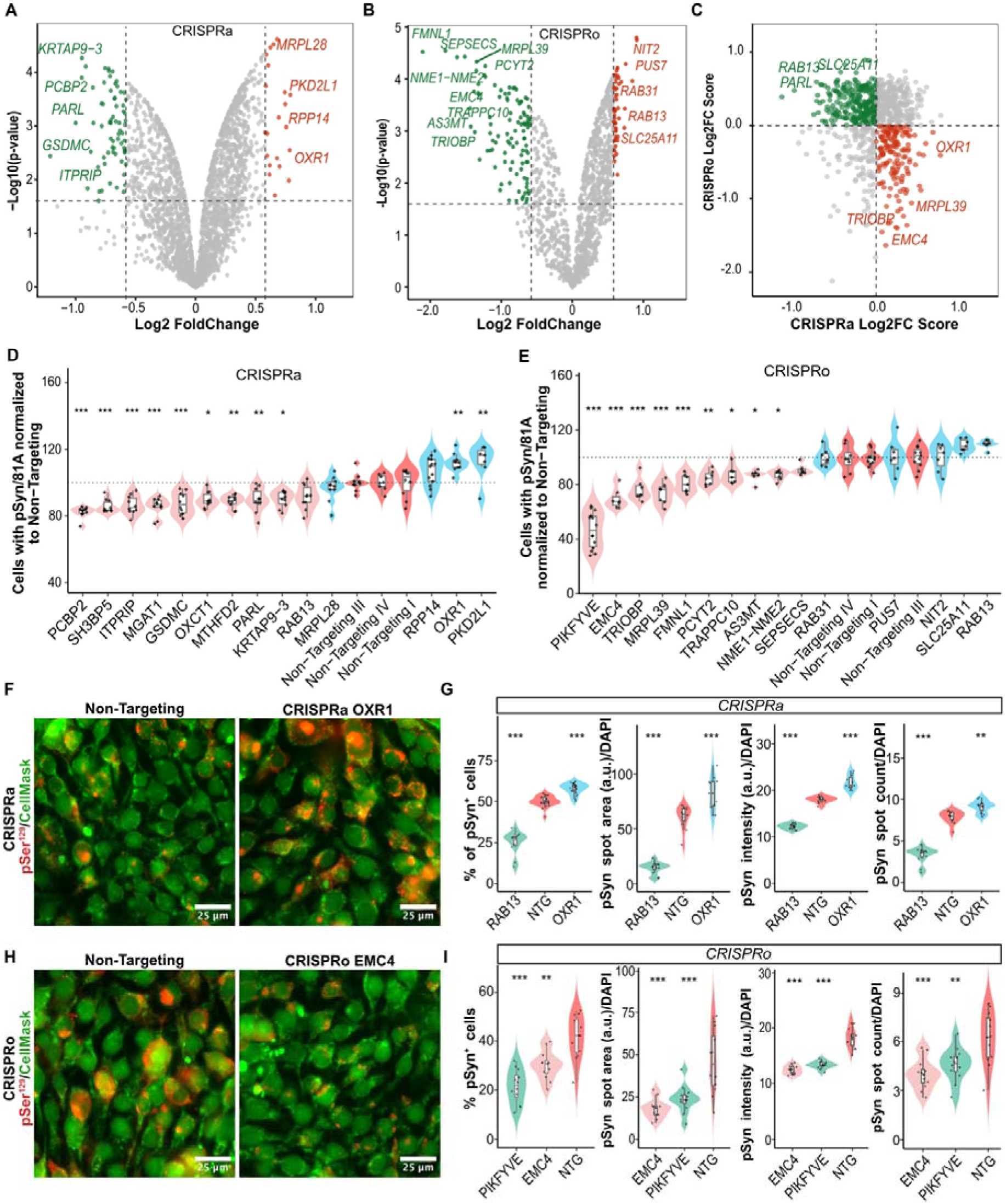
CRISPRa/CRISPRo screens for genetic modulators of synuclein aggregation. **(A-B)** Volcano plots highlighting genes increasing (red) or decreasing (green) pSyn^129^ levels. **(C)** Cross-tabulation of effect sizes in CRISPRa vs CRISPRo screens. Green and red: bidirectional modifiers. **(D-E)** Validation of hits with increased replicates. **(F, H)** pSyn^129+^ aggregates in HEK^Syn^ cells following OXR1 activation (F) or EMC4 ablation (H). CellMask: red; pSyn^129+^ aggregates: yellow; DAPI: blue. **(G, I)** Quantification of pSyn^129+^ cells, aggregate area arbitrary units (a.u.) per DAPI, aggregate intensity (a.u.) per DAPI, and aggregate count (per DAPI) following OXR1 activation (G) or EMC4 (I) ablation. NTG: Non-Targeting. Inner box plots display the median (centre line), the 75th and 25^th^ percentile (top and bottom edge). One-way ANOVA followed by Dunnett’s post hoc test; *P < 0.05, **P < 0.01, ***P < 0.001.

### Secondary screen identified EMC4 and OXR1 as strong modulators of pSyn^129^

To refine our candidate selection and eliminate false positives, we retested all selected hits using additional technical replicates. Each candidate gene was statistically compared to the non-targeting control group using a one-way ANOVA followed by Dunnett’s post hoc test. Most hits maintained the same directional trend observed in the primary screen, with 11 of 13 CRISPRa hits and 9 of 14 CRISPRo hits reaching statistical significance (p < 0.05; **Figures 2D-2E**). In the CRISPRa screen, *PCBP2, SH3BP5, ITPRIP, MGAT1, GSDMC, OXCT1, MTHFD2, PARL,* and *KRTAP9-3* were confirmed as pSyn^129^ downregulators whereas *OXR1* and *PKD2L1* emerged as upregulators (**Figure 2D**). For the CRISPRo screen, *EMC4, TRIOBP, MRPL39, FMNL1, PCYT2, TRAPPC10, AS3MT,* and *NME1-NME2* were confirmed as pSyn^129^ downregulators (**Figure 2E**). *RAB13* was identified as an upregulator alongside *SLC25A11*, although the latter did not reach statistical significance.

To determine whether any of our hits have previously been implicated in PD, we obtained a PD-associated gene set from the Open Targets Platform (https://platform.opentargets.org/). Open Targets dataset compiles genes from GWAS and expression studies, as well as literature-curated and other disease-linked genes. Among our validated hits, only GSDMC (Gasdermin C) overlapped with the genes from the Open Targets dataset (**Figure S2H**) (Jiang *et al*, 2020; Nalls *et al*, 2019); however, its modulation of pSyn^129^ was minimal and was not further investigated. Instead, we focused on the two strongest and most reproducible modifiers: the mitochondrial Oxidation Resistance Gene 1 (*OXR1*) and the ER Membrane Protein Complex Subunit 4 (*EMC4/TMEM85*). Both genes are highly expressed in the human brain, including neurons (https://www.proteinatlas.org/) (Sjöstedt *et al*, 2020). CRISPR-based manipulation of *OXR1* and *EMC4* did not affect cellular viability as indicated by lactate dehydrogenase (LDH) release assay (**Figures S3A–S3B**). pSyn^129^ was assessed with the 81A and EP1536Y antibodies. *OXR1* upregulation caused a significant increase in the fraction of pSyn^129^⁺ cells, total pSyn^129^ spot area, total pSyn^129^ spot intensity, and the number of pSyn^129^ puncta divided by the number of DAPI+ nuclei (**Figures 2F–2G, 3A, and S3D**). This effect was further supported by flow cytometry, which showed a consistent trend, albeit with lower effect size (**Figure S4A-S4B**). Western blotting confirmed increased OXR1 protein levels in CRISPRa-activated cells (**Figure S2K**).

Similarly, ablation of *EMC4* significantly reduced the fraction of pSyn^129^⁺ cells, total pSyn^129^ spot area, total pSyn^129^ spot intensity, and the number of pSyn^129^ puncta (**Figures 2H–2I, 3B, and S3E**). Similar results were obtained by flow cytometry (**Figures S4A-S4C**). Although CRISPR-based ablation of *EMC4* led to a substantial reduction in its mRNA levels, we were unable to achieve complete depletion of the EMC4 protein (**Figures S2I, S2J**). Attempts to isolate stable clonal lines lacking EMC4 entirely were unsuccessful: the few clones that initially survived showed poor viability and failed to expand beyond one or two passages, suggesting that complete *EMC4* ablation may compromise cell fitness. This is consistent with data from the International Mouse Phenotyping Consortium (MGI:1915282), which indicates that homozygous *EMC4* knockout severely compromises viability in mice. Nonetheless, the partial EMC4 depletion was sufficient to significantly reduce pSyn^129^⁺ prevalence.

We then tested the effect of partially reducing *EMC4* expression through a shRNA-based approach, rather than CRISPR. Knockdown of *EMC4* through shRNA recapitulated the phenotypic effects observed in the CRISPRo experiments, including significant reduction in pSyn^129^ positive cells (**Figures 3C and S3C**). This effect was accompanied by a marked reduction in *EMC4* mRNA and protein levels (**Figures 3D, 3E**). We next examined potential bidirectional effects but found that OXR1 ablation and EMC4 activation did not alter pSyn^129^ levels in HEK^Syn^ cells (**Figures S5C, S5D**).

**Figure 3:**
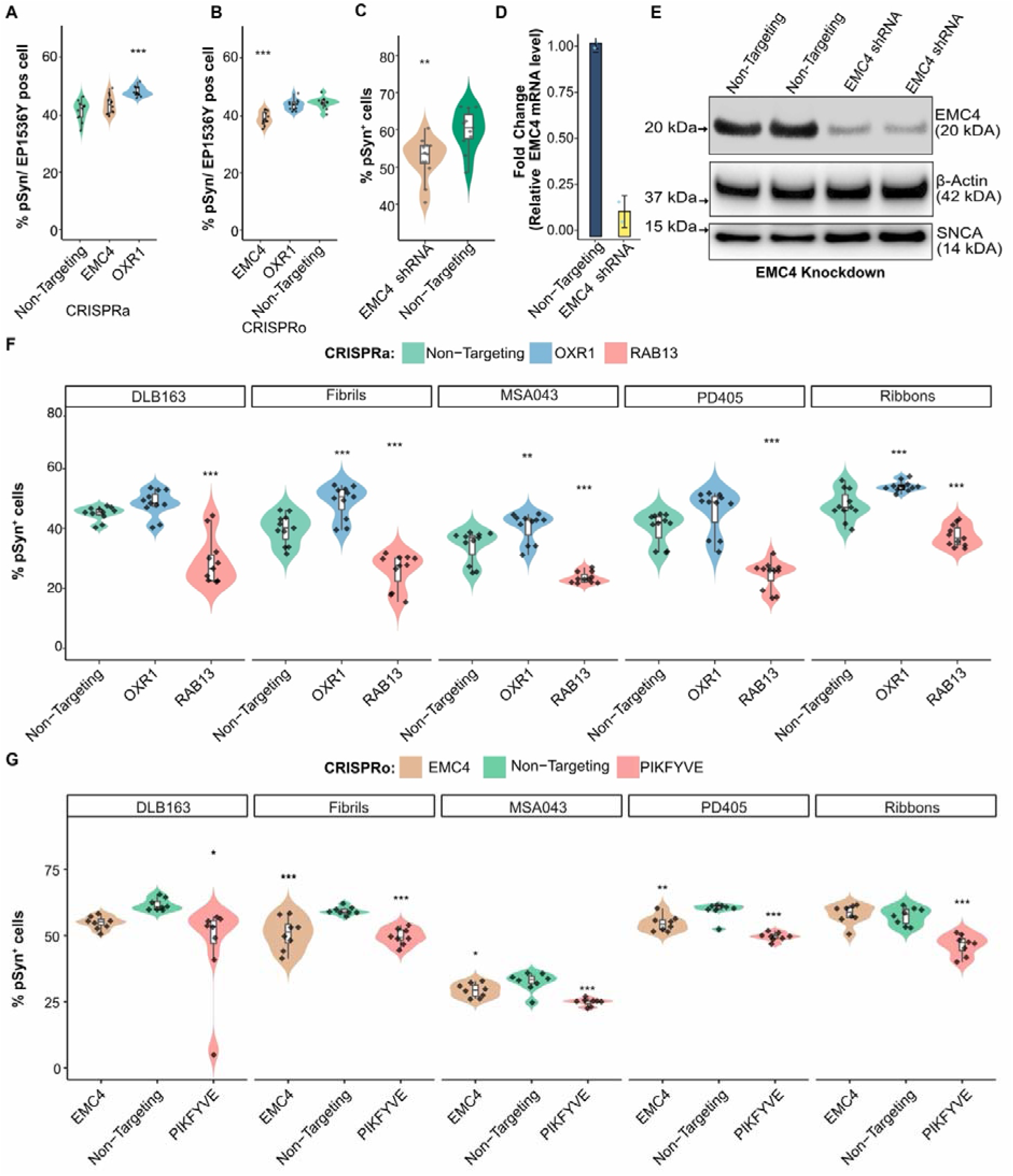
Differential effects of modifiers across different patient-derived and recombinant fibrillar strains and orthogonal hit validation. **(A-B)** pSyn^129^ levels following EMC4 and OXR1 perturbations (EP1536Y antibody). **(C)** shRNA-mediated inhibition of EMC4 and its impact on pSyn^129^ (81A antibody). Welch’s t-test (unequal variance t-test, *P < 0.05, **P < 0.01). **(D)** EMC4 mRNA levels after shRNA-mediated knockdown (RT-qPCR, Mean ± SEM). **(E)** Western blot showing EMC4 protein and SNCA protein after shRNA-mediated inhibition. **(F-G)** Effect of OXR1 activation (f) or EMC4 ablation (G) on pSyn^129^ for human PD, DLB, and MSA patient-derived fibrils as well as recombinant fibrils and ribbons (81A antibody). Inner box plots display the median (centre line), the 75th percentile (top edge), and the 25th percentile (bottom edge). One-way ANOVA followed by Dunnett’s post hoc test. *P < 0.05, **P < 0.01, ***P < 0.001.

### Strain-specific effects of OXR1 activation and EMC4 depletion on pSyn^129^ levels

Synucleinopathies encompass diverse disorders, each defined by structurally distinct α-Syn assemblies. These strains exhibit unique pathogenic properties, reflecting their clinical origins (Bousset *et al*, 2013; Van Der Perren *et al*, 2020). We tested the effect of *OXR1* and *EMC4* perturbation across diverse α-Syn assemblies: patient-derived strains (PD, MSA, DLB) and recombinant polymorphs (fibrils, ribbons). Activation of OXR1 led to a moderate increase in pSyn^129^⁺ prevalence across all strains, with statistically significant effects observed for recombinant polymorphs and MSA-derived fibrils (**Figures 3F and S5A**), suggesting that strain conformation influences OXR1-mediated aggregate accumulation. In contrast, EMC4 depletion significantly reduced pSyn^129^⁺ prevalence across PD, MSA, and fibrillar strains (**Figures 3G and S5A**), supporting its function as a broad-spectrum modulator of α-Syn aggregation.

### OXR1 overexpression impairs ATP synthesis and elevates pSyn^129^ accumulation

OXR1 is involved in several key processes that maintain neuronal homeostasis, including protection from oxidative stress and DNA repair (Volkert & Crowley, 2020). To identify the downstream effectors responsible for its modulation of pSyn^129^, we compared the transcriptome of OXR1-overexpressing HEK^Syn^ cells against HEK^Syn^ cells transduced with NTG CRISPR guides. Differentially expressed genes (DEGs) were defined by stringent filtering criteria (FDR ≤ 0.05, log2FC ≥ 0.5 or ≤ -0.5, p ≤ 0.01; **Figure 4A**; **Table S3)**. *SNCA* mRNA levels were not changed by *OXR1* activation, suggesting that OXR1 modulates pSyn^129^ levels posttranscriptionally or through indirect mechanisms (**Figure 4A**). Next, we individually activated or ablated the top upregulated (*FMOD, CCL8, SALL3, SOCS3, ISX, CRLF1, MYH6*) and downregulated (*B3GNT3, H3Y1, ALDOC, RFPL4A*) genes to determine how each hit influences pSyn^129^ levels in CRISPRa HEK^Syn^ cells exposed to α-Syn PFFs. pSyn^129^ was quantified by flow cytometry following 81A antibody staining (**Figure S6A**). Activation of OXR1-induced genes, *FMOD* (fibromodulin, an extracellular matrix remodeling protein) and *CCL8* (C-C Motif Chemokine Ligand 8, an immune signaling molecule), resulted in an increased pSyn^129+^ prevalence (**Figure 4B**). Similarly, ablation of the *OXR1-*repressed genes *ALDOC* (aldolase C, a key glycolytic enzyme cleaving fructose-1,6-bisphosphate) and *RFPL4A* (ubiquitin protein ligase activity) also increased pSyn^129+^ prevalence (**Figures 4C and S6B**). Additionally, flow cytometry results were further confirmed by immunofluorescence imaging, which showed concordant changes in pSyn^129^ levels for all four genes (**Figures S6C-S6D**). These genes share OXR1 as a regulator and, when perturbed individually, promote pSyn^129^ accumulation.

**Figure 4:**
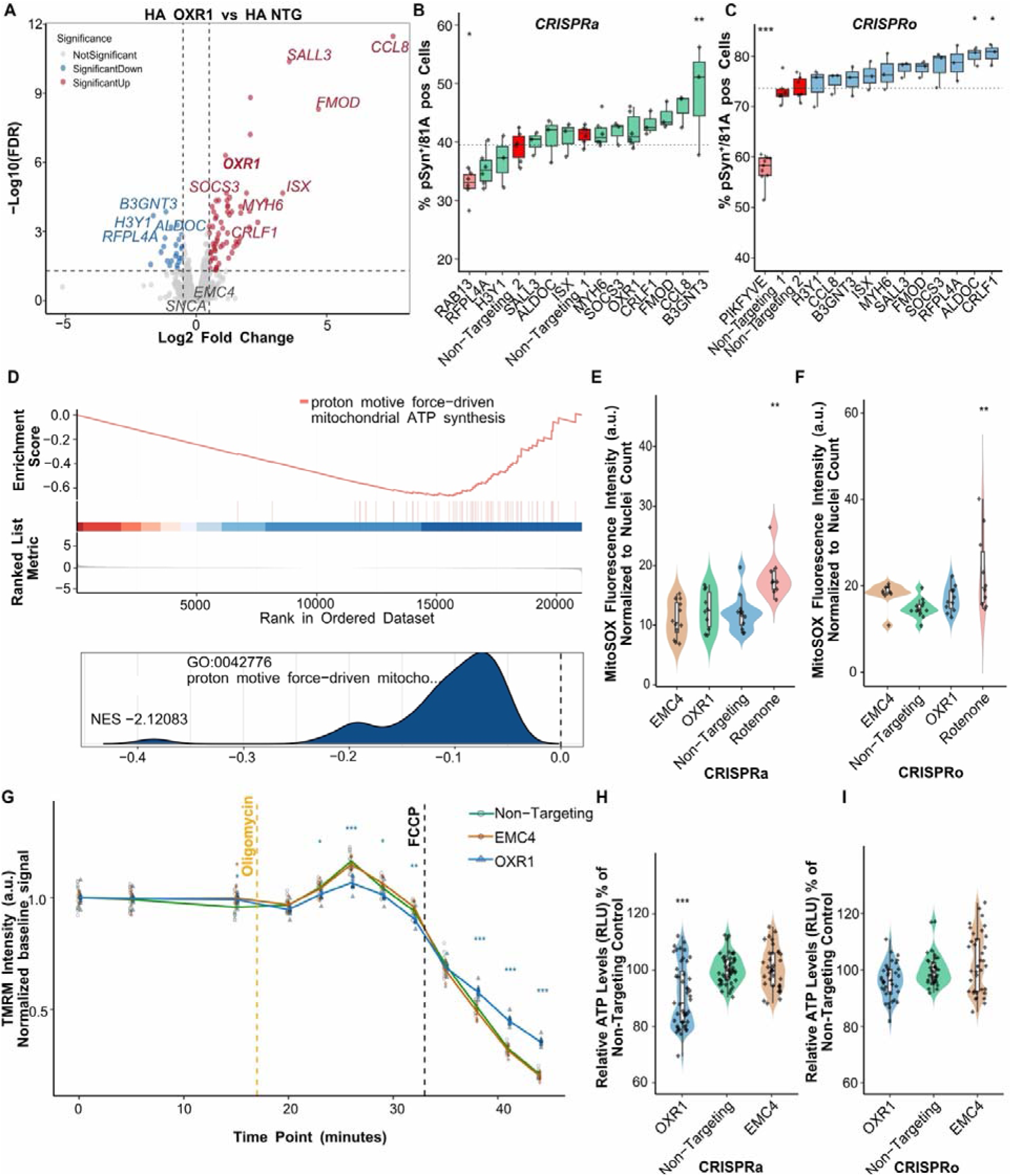
Transcriptional and functional consequences of OXR1 activation. **(A)** Transcripts up (red) and downregulated (blue) by OXR1 activation. NTG: non-targeting control **(B-C)** Individual perturbation of DEGs by OXR1 activation. **(D)** Gene set enrichment analysis (GSEA) highlighting the mitochondria-related pathway with the normalized enrichment score (NES). Candidate terms are based on a false-discovery rate ≤ 0.05; genes are ranked by effect size. Upper panel: running enrichment score plot; lower panel: ridge plot. **(E,F)** Mitochondrial superoxide levels measured by MitoSOX™ Red dye in live cells. Fluorescence intensity was normalized to the total number of cells. (G) Time-resolved measurement of mitochondrial membrane potential (ΔΨm) using TMRM fluorescence under basal conditions and after sequential exposure to oligomycin and FCCP. **(H, I)** Intracellular ATP levels measured using CellTiter-Glo (RLU: Relative Luminescence Units). Percentage of pSyn^+^ cells following CRISPR activation (H) and CRISPR ablation (I). Violin plots represent data distribution. Inner box plots display the median (centre line), the 75th percentile (top edge), and the 25th percentile (bottom edge). One-way ANOVA followed by Dunnett’s post hoc test; *P < 0.05, **P < 0.01, ***P < 0.001.

Gene Set Enrichment Analysis (GSEA) of RNA-seq data from HEK^Syn^ cells overexpressing OXR1 showed a significant enrichment of downregulated genes associated with the proton motive force-driven mitochondrial ATP synthesis pathway (NES = -2.12083) (**Figures 4D and S7A**). The running-enrichment curve indicates that genes in this pathway are significantly enriched among the most down-regulated transcripts (**Figure 4D**, upper panel). Key components of this pathway include *ATP5F1A, ATP5F1B* (ATP synthase subunits), *NDUFS2, NDUFS4* (Complex I), and *MT-ATP6* (mitochondrial ATP synthase). The accompanying ridge-density plot (**Figure 4D**, lower panel) shows the distribution of pathway genes shifted toward negative log2FC values, confirming collective repression in OXR1-overexpressing cells. Since these genes are linked to mitochondrial function, we examined whether OXR1 overexpression alters two key indicators of mitochondrial health: reactive oxygen species (ROS) production and membrane potential. We used MitoSOX™ Red, a fluorescent probe that selectively detects superoxide in mitochondria, to determine if OXR1-induced changes might elevate oxidative stress. We found no significant difference in MitoSOX fluorescence between OXR1-overexpressing HEK^Syn^ and HEK^Syn^ cells transduced with NTG guides (**Figures 4E–4F and S7B–S7C)**. Next, we evaluated mitochondrial membrane potential (ΔΨm) using TMRM, a dye accumulating in polarized mitochondria whose fluorescence intensity reflects membrane potential. To disrupt mitochondrial polarization, we treated cells with oligomycin to inhibit ATP synthase, followed by FCCP to induce complete depolarization (Vianello *et al*, 2023). Live-cell imaging with TMRM showed minimal impact of OXR1 activation on mitochondrial membrane potential (ΔΨm) under basal conditions (**Figures 4G and S7D**). A transient increase in ΔΨm at 26 minutes (***p < 0.001) did not persist after FCCP-induced depolarization, suggesting no sustained effect on mitochondrial polarization. These findings suggest that although OXR1 overactivation downregulates components of the ATP synthase machinery, it does not markedly alter mitochondrial ROS production and exerts only a modest, transient effect on basal membrane potential. Since *OXR1* overexpression downregulated components of ATP synthase, we measured total ATP levels using CellTiter-Glo to test whether overall cellular ATP production was affected (**Figures 4H-4I**). *OXR1* activation reduced total ATP levels by ∼15% (Dunnett’s post-hoc test, p = 2.9 × 10⁻□) compared to non-targeting controls (**Figure 4H**), suggesting a functional role in mitochondrial ATP synthesis. However, *OXR1* ablation did not impact mitochondrial ATP production (**Figure 4I**). Together, these findings show that OXR1 overexpression modestly reduces ATP levels, and mitochondrial membrane potential, while additional regulation by *FMOD*, *CCL8*, *ALDOC*, and *RFPL4A* underscores the interplay between mitochondrial function and gene-specific modulators in driving pSyn^129^ accumulation.

### OXR1 activation exacerbates pSyn^129^ pathology in human iPSC-derived cortical neurons

To test our hits in a disease-relevant neuronal model, we assessed the effect of OXR1 activation in neurons derived from human iPSC line with an inducible NGN2 (neurogenin 2) cassette, and CRISPRa machinery dihydrofolate-reductase destabilising domain (DHFR) fused with transactivator dCas9-VPH (Tian *et al*, 2021). Following the lentiviral delivery of CRISPRa qgRNAs, the iPSCs were differentiated into integrated, isogenic, inducible (i3) glutamatergic cortical neurons (Fernandopulle *et al*, 2018; Wang *et al*, 2017). CRISPRa activity was then triggered by adding trimethoprim (TMP), which stabilises the DHFR-dCas9-VPH fusion. The resulting iPSC-derived cortical neurons were treated with α-Syn PFFs for four weeks (**Figure S8A**). Immunofluorescence staining revealed pSyn^129^ accumulation in both soma and neurites, exclusively within MAP2-positive regions (**Figure 5A**). OXR1 activation significantly increased the pSyn^129+^ puncta area in both compartments (**Figure 5B**). Western blot analysis confirmed OXR1 upregulation in CRISPRa iPSC-derived cortical neurons (**Figure 5C**).

**Figure 5:**
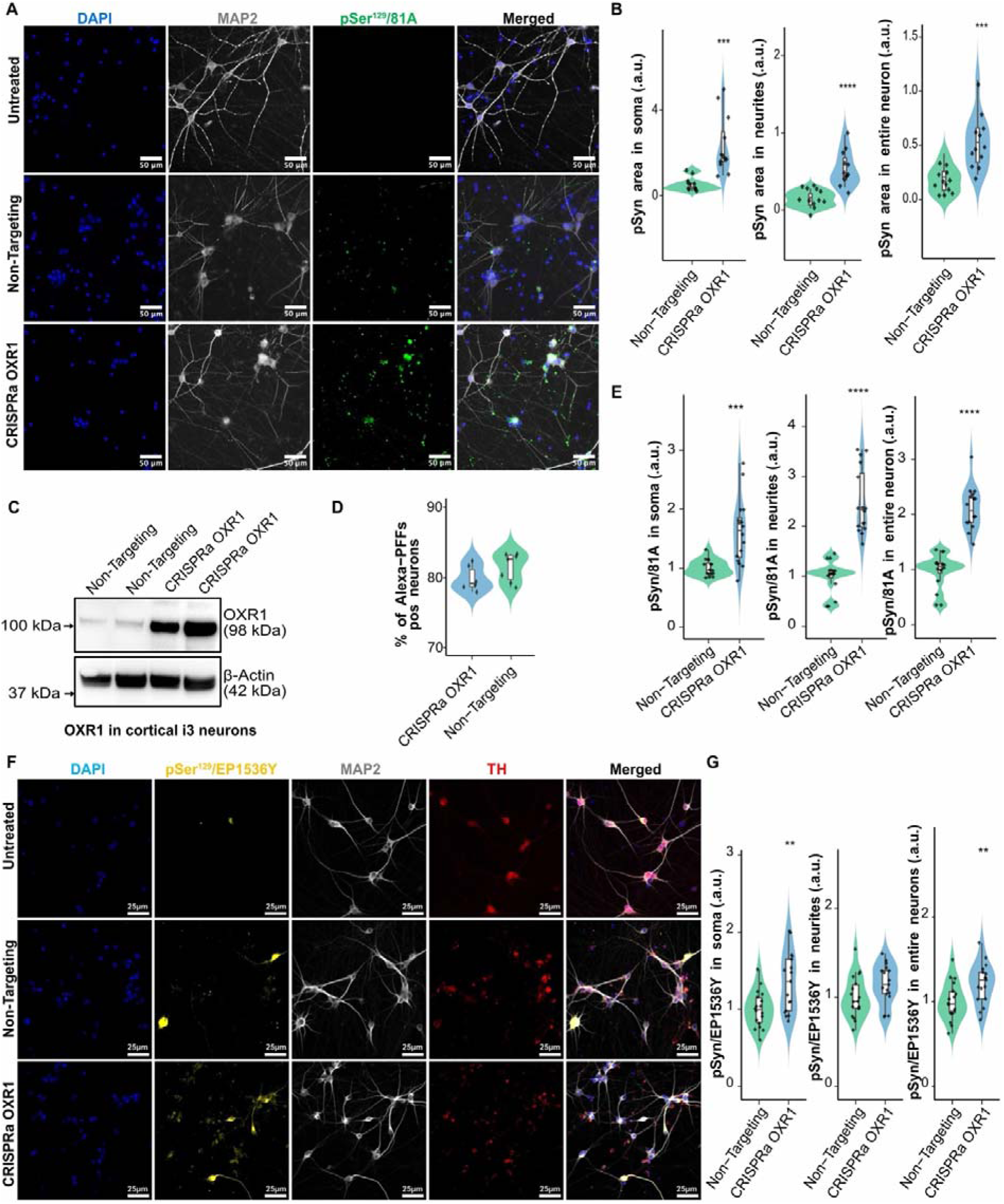
OXR1 activation modulates phosphorylated **α**-Synuclein at Ser129 (pSyn^129^) in human iPSC-derived cortical and dopaminergic (iDA) neurons. **(A)** iPSC-derived cortical neurons stained with DAPI (blue), MAP2 (grey), and pSyn^129^ (81A, green). **(B)** Quantification of pSyn^129^ spot area in soma (left) neurites (middle) and whole neurons (right) per total MAP2 area. **(C)** Immunoblot confirming OXR1 expression in iPSC derived cortical neurons. **(D)** Flow cytometry of Alexa Fluor 488-labelled PFFs uptake in iPSC derived cortical neurons. **(E)** pSyn^129^ levels in iDA neurons in soma, neurites, and whole neurons. **(F)** iDA neurons stained with DAPI (blue), MAP2 (grey), EP1536Y (yellow, pSyn^129^ aggregates), and tyrosine hydroxylase (TH; red, dopaminergic marker). **(G)** Quantification of pSyn^129^ spot area in soma, neurites and in entire iDA neurons normalized to MAP2 area. Box plots: median (center line), 75th), and 25th percentile (top and bottom edges). Welch’s t-test (unequal variance t-test); *P < 0.05, **P < 0.01, ***P < 0.001.

Next, we investigated whether *OXR1* activation enhances PFF internalization. We quantified Alexa Fluor 488-labelled PFFs in iPSC-derived cortical neurons treated for 6 hours via flow cytometry. No differences were observed between *OXR1*-activated iPSC-derived cortical neurons and control (NTG) iPSC-derived cortical neurons (**Figures 5D and S8C**), ruling out PFF uptake modulation as the cause of enhanced pSyn^129^ accumulation. Finally, Western blot analysis further confirmed that OXR1 activation does not alter endogenous SNCA expression (**Figure S8B**), reinforcing that increased pSyn^129^ accumulation is not due to elevated α-Syn levels.

### OXR1 activation promotes pSyn^129^ accumulation in human iPSC-derived dopaminergic neurons

As pSyn^129^ accumulation mostly affects the dopaminergic neurons of the substantia nigra, we differentiated NGN2 dCas9VPH-expressing iPSCs into dopaminergic neurons (iDA) using a rapid NGN2-based differentiation protocol (**Figure 5F**), which yields a homogeneous population of tyrosine hydroxylase (TH)-positive neurons with mature dopaminergic features (Sheta *et al*, 2023). To quantify pSyn^129^ accumulation in OXR1-overexpressing iDA neurons, we used the antibodies 81A and EP1536Y. Notably, 81A staining showed increased pSyn^129^ accumulation in both the soma and neurites, leading to an overall increase in total neuronal pSyn^129^ burden (**Figures 5E and S9A**). In contrast, EP1536Y staining revealed a significant increase in pSyn^129^ signal in neuronal somata (**Figures 5F-5G**), while neuritic levels exhibited a similar trend that did not reach statistical significance (**Figure 5G**).

### EMC4 ablation enhances lysosomal but not proteasomal degradation

We next sought to clarify the mechanism by which EMC4 ablation reduces pSyn^129^ levels. To identify the downstream mechanisms by which EMC4 ablation decreases pSyn^129^, we sequenced mRNA of EMC4-ablated HEK^Syn^ cells and compared to that of CRISPRo HEK^Syn^ cells transduced with NTG CRISPR guides. DEGs were selected following the same criteria used for OXR1 upregulation (FDR ≤ 0.05, log2FC ≥ 0.5 or ≤ -0.5, p ≤ 0.01) (**Figure 6A**; **Table S4**). Several genes were significantly upregulated, including *ADCYAP1R1, H2BC11, H4C14, IFI30, KCNN2, KLHL38, OTOP2, PIK3R2,* and *PSMB9*. Downregulated genes included *CGA, DNAJC17, EMC4, HYPK, MYO5C, RORA, SLC27A2,* and *TPM1*. Notably, neither α-Syn mRNA (**Figure 6A**) nor α-Syn protein levels changed upon EMC4 depletion (**Figure 3E**), indicating that EMC4 depletion modulates pSyn^129^ levels independently of endogenous SNCA expression.

**Figure 6:**
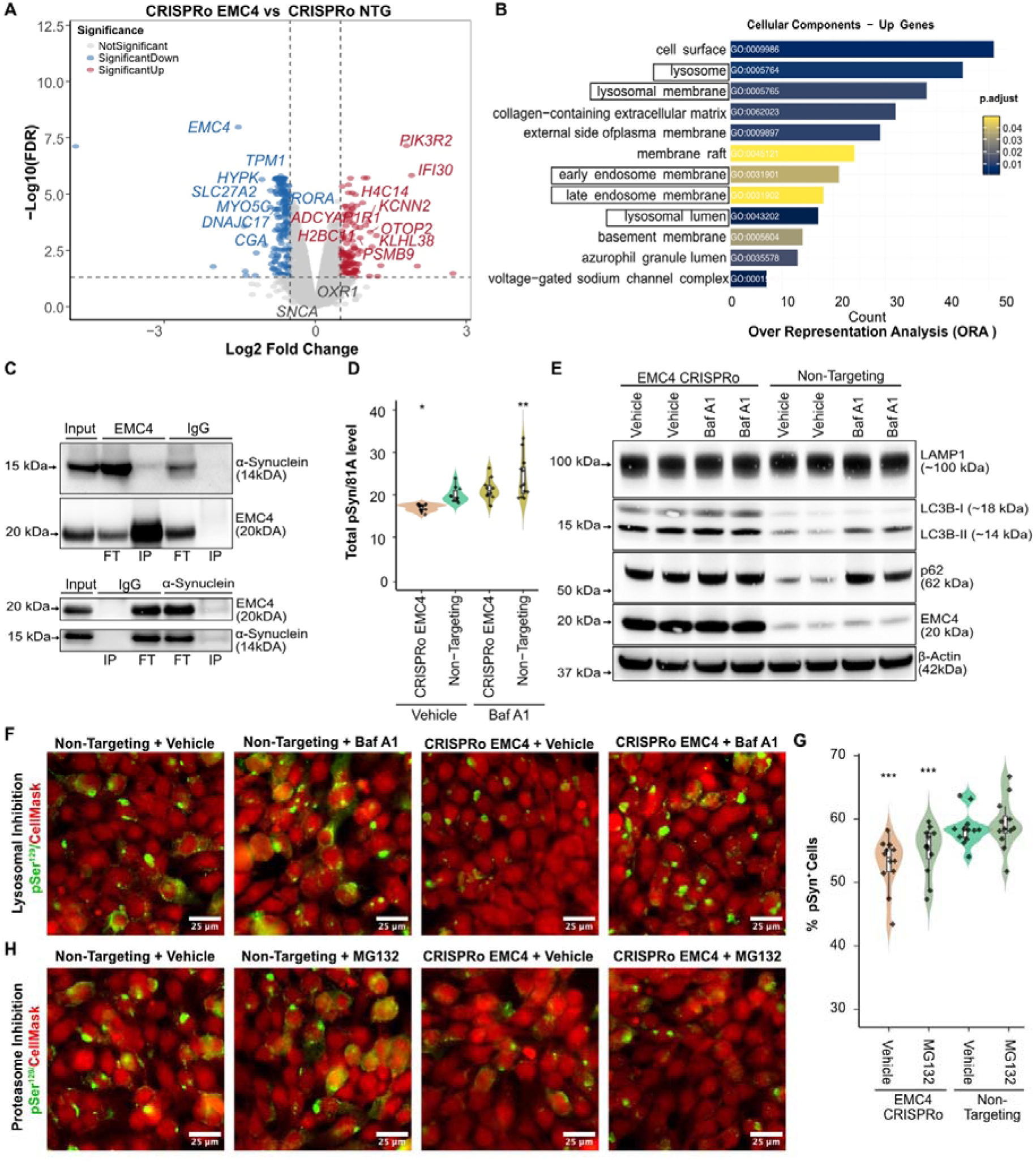
EMC4 ablation and lysosomal function. **(A)** RNA sequencing showing DEGs between CRISPRo EMC4 and CRISPRo non-targeting (NTG) controls. **(B)** Hypergeometric ORA of cellular compartments with FDR ≤ 0.05 associated with upregulated genes. **(C)** Bidirectional co-immunoprecipitation (Co-IP) of EMC4 and α-synuclein (α-Syn). Top: Western blots of EMC4 immunoprecipitates probed for EMC4 or α-Syn. Bottom: Reciprocal Co-IP showing α-Syn immunoprecipitation and detection of both EMC4 and α-Syn. Input lanes: lysate prior to immunoprecipitation, FT (flow-through) and IP (immunoprecipitated) fractions. **(D)** Quantification of lysosomal inhibition (Baf A1) effects on total phosphorylated α-Synuclein at Ser129 (pSyn129) spot intensity levels, normalised to the total cell number in CRISPRo EMC4 cells. DMSO was used as a vehicle control. **(E)** Western blot analysis of lysosomal and autophagy markers, including LAMP1, LC3B-I/II, and p62, in CRISPRo EMC4 and CRISPRo NTG cells treated with vehicle, Baf A1. **(F)** Representative immunofluorescence images showing the effect of lysosomal inhibition with Baf A1. **(G)** Quantification of pSyn129^+^ cells under proteasomal inhibition (MG-132) in EMC4 ablation and NTG conditions. **(H)** pSyn129^+^ cells under proteasomal inhibition (MG-132) in CRISPRo EMC4 and NTG conditions. (Red: HCS CellMask; Green: pSyn129/81A). Inner box plots display the median (centre line), the 75th percentile (top edge), and the 25th percentile (bottom edge). One-way ANOVA followed by Dunnett’s post hoc test; *P < 0.05, **P < 0.01, ***P < 0.001.

Next, we tested the top 8 DEGs (upregulators and downregulators) for their effect on pSyn^129^ accumulation. Most of them were associated with lysosomal or proteostatic pathways. Several candidates influenced pSyn^129^ prevalence upon individual perturbation, with some exhibiting bidirectional effects under opposite perturbations. Notably, overexpression of the lysosomal thiol reductase *IFI30* significantly reduced pSyn^129^ load (**Figure S10A**). Thiol reductases help unfold proteins destined for lysosomal degradation. Hence, *IFI30* overexpression may facilitate the disassembly of pSyn^129^ aggregates prior to lysosomal digestion. Additionally, genetic ablation of *DNAJC17,* encoding an essential chaperone, decreased pSyn^129^ accumulation, while its activation promoted pSyn^129^ accumulation (**Figures S10A–S10D**). *DNAJC17* mRNA levels decreased following EMC4 ablation, whereas EMC4 levels remained unaffected by *DNAJC17* depletion, suggesting that *DNAJC17* may act downstream of *EMC4* (**Figures S10E–S10G**). Combined ablation of *EMC4* and *DNAJC17* showed no additive effects, indicating that they may share the same pathway (**Figure S10H**).

We next sought to identify broader pathway-level changes underpinning EMC4’s mechanism of action. Hypergeometric over-representation analysis (ORA) revealed significant enrichment of lysosome-associated cellular components (lysosome, membrane/lumen), as well as endosomal transport pathways and dysregulation of both early and late endosomal compartments (**Figure 6B**). Moreover, we observed upregulation of ER stress response pathways (**Figures S11A–S11B**) and genes associated with ER homeostasis including *HSPA5*, *DNAJC3*, *DNAJB9*, and *PIK3R2* (**Figure S11C**). Since our transcriptomic profiling highlighted an enrichment of lysosome-associated processes (**Figure 6B**), and given that the EMC complex is also implicated in ER-associated degradation (ERAD), we hypothesized that EMC4 depletion might reduce pSyn^129^ levels by promoting its degradation via these pathways. To test this, we first inhibited lysosomal activity using Bafilomycin A1 (BafA1), a selective inhibitor of lysosomal acidification. The decrease in pSyn^129^ levels following EMC4 depletion was suppressed by BafA1 treatment (**Figures 6D–6F**), indicating that the reduction was lysosome-dependent. EMC4-depleted HEK^Syn^ cells exhibited reduced levels of LC3B-II (**Figure 6E**), a marker of autophagic flux, and p62/SQSTM1, a cargo receptor for autophagic degradation, consistent with increased lysosomal clearance and turnover. Treatment with BafA1 restored both LC3B-II and p62 levels. Hence, we speculate that pSyn^129^ reduction upon EMC4 ablation proceeds through an increase in the autophagic flux and enhanced lysosomal degradation. No alterations in LAMP1 levels were observed, suggesting that EMC4 ablation does not broadly impact lysosomal biogenesis. Secondly, to determine whether *EMC4* ablation also enhances proteasomal degradation, we treated cells with the proteasomal inhibitor MG132. Unlike BafA1, MG132 treatment failed to restore pSyn^129^ levels (**Figures 6G–6H**), indicating that *EMC4* depletion does not promote proteasomal clearance of pSyn^129^. Interestingly, co-immunoprecipitation (Co-IP) analysis revealed a physical interaction between EMC4 and α-Syn. Immunoprecipitation of EMC4 co-precipitated α-Syn, and vice versa, immunoprecipitation of α-Syn enriched EMC4 in the pull-down fraction (**Figure 6C**). Specificity was confirmed using IgG controls, which showed no interaction. These results establish EMC4 as a regulator of lysosome-mediated aggregated α-Syn proteostasis and highlight ER stress–lysosomal crosstalk as a key pathway in synucleinopathies.

### EMC4 depletion reduces pSyn^129^ levels in iPSC-derived cortical neurons

To investigate the effects of *EMC4* depletion in iPSC-derived cortical neurons, we used shRNA to deplete its expression (**Figures 7A–7B**). shRNA-mediated EMC4 depletion resulted in a significant reduction in pSyn^129^ levels, consistent with the effect observed in HEK cells. No differences were detected in PFF internalization between EMC4-depleted and scrambled-shRNA iPSC-derived cortical neurons (**Figures 7C and S12A**), confirming that the reduced pSyn^129^ accumulation cannot be explained by altered PFF uptake. Efficient knockdown of EMC4 was confirmed by RT-qPCR (**Figure 7D**) and by immunoblotting, which demonstrated an ∼75% reduction in EMC4 protein (**Figure 7E**). Consistent with findings in HEK^Syn^ cells, *EMC4* knockdown in iPSC-derived cortical neurons also led to reduced LC3B-II levels, while LAMP1 expression remained unchanged (**Figure 7E**), supporting engagement of the autophagy-lysosome pathway. These data, obtained in disease-relevant neuronal models, validate the lysosome-dependent mechanism of pSyn^129^ clearance observed in *EMC4*-ablated HEK^Syn^ cells.

**Figure 7:**
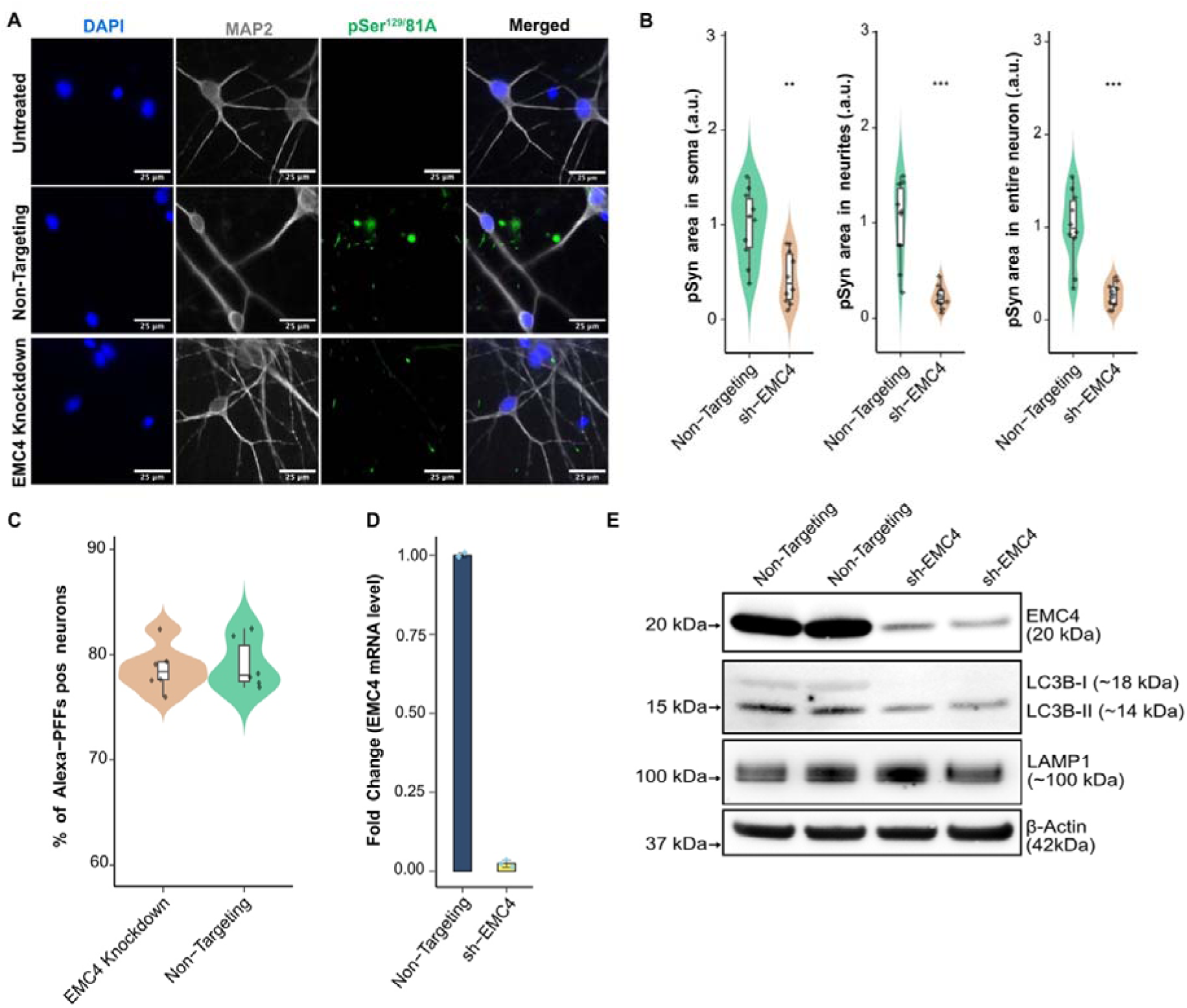
EMC4 depletion modulates phosphorylated α-Synuclein at Ser129 (pSyn^129^) levels in human iPSC-derived cortical neurons. **(A)** Neurons subjected to EMC4 shRNA or to control conditions. DAPI (blue), MAP2-labeled neurons (grey), and 81A stained pSyn^129^ aggregates (green). **(B)** Quantification of pSyn^129^ spot area: (left) in soma, (middle) neurites and (right) whole neurons normalized to total MAP2 area. **(C)** Flow cytometry analysis of fluorescent PFF uptake by neurons. **(D)** RT-qPCR analysis of EMC4 mRNA levels in knockdown neurons. Bar plots data are presented as mean ± SEM. **(E)** Western blot analysis of lysosomal and autophagy markers in cortical neurons after shRNA-mediated knockdown of EMC4. Violin plots represent data distribution. Box plots display the median (center line), the 75th percentile (top edge), and the 25th percentile (bottom edge). Welch’s t-test (unequal variance t-test), *P < 0.05, **P < 0.01, ***P < 0.001.

## Discussion

Strong genetic evidence indicates that α-Syn aggregation is a major contributor to PD, MSA, and DLB. Hence, the identification of targetable modifiers of this process may help devise causal treatments for these ailments (Calabresi *et al*, 2023; Neupane *et al*, 2023; Oliveira *et al*, 2021). Yet, few such modifiers are known. We reasoned that large-scale interrogations of appropriate cellular models of disease may help discover wholly new actors, including those that may not be identifiable by studies of human genetics. We therefore deployed arrayed dual CRISPR activation/ablation screens targeting all genes associated with mitochondrial dynamics and intracellular trafficking, and assessed their impact on pSyn^129^ accumulation, a key hallmark of synucleinopathies.

In contrast to pooled CRISPR screens, image-based arrayed CRISPR screens provide single-cell resolution for thousands of cells, enabling the statistically robust detection of subtle phenotypes. However, they can suffer from replicability issues due to imaging artefacts, signal fluctuations and missegmentation during image analysis. Furthermore, despite the high specificity of CRISPR-based gene perturbations, off-target effects and false positives remain challenging in large-scale screens (Bock *et al*, 2022). We mitigated these limitations by using qgRNAs per gene to ensure targeting redundancy. Additionally, we implemented multiple orthogonal validation strategies, robust image analysis pipelines, as well as multiple distinct cell-based models (HEK293 cells and iPSC-derived neurons) to ensure biological relevance. HEK293 cells are often described as epithelial, yet they likely arose from neuroendocrine cells (Shaw *et al*, 2002), express specific neuronal markers, and are responsive to neuronal stressors. Though imperfect, these neuron-like characteristics, in combination with their rapid cell cycles and ease of culture, make HEK293 cells suitable for genetic screens related to neurological diseases.

To maximize discovery of novel modulators, we focused our unbiased screens on genes implicated in MTM function or proteostasis, without further prioritization. This approach repeatedly identified the mitochondrial *OXR1* and ER-associated *EMC4* proteins as regulators of pSyn^129^. OXR1 activation exacerbated α-Syn pathology via compromised mitochondrial function, whereas EMC4 depletion facilitated α-Syn clearance through lysosomal degradation independently of proteasomal activity.

Mitochondrial dysfunction is a hallmark of PD and is linked to the accumulation of pSyn^129^. Recent studies suggest a self-reinforcing cycle: pSyn^129^ accumulation impairs mitochondrial Complex I and increases ROS, exacerbating pSyn^129^ accumulation (Ganjam *et al*, 2019; Geibl *et al*, 2024), while mitochondrial malfunction and ATP depletion promote α-Syn misfolding (Risiglione *et al*, 2021). This bidirectional crosstalk creates a pathological loop, yet the initial trigger—mitochondrial insult or α-Syn toxicity—remains unclear (Gao *et al*, 2022). We observed that OXR1 activation coincided with elevated pSyn^129^ levels and was associated with reductions in mitochondrial membrane potential and ATP synthesis. This may impair ATP-dependent proteostasis mechanisms, such as chaperone-assisted refolding and proteasome-mediated degradation of misfolded proteins (Goldberg, 2003). As a result, α-Syn misfolds and accumulates. This is consistent with prior evidence of mitochondrial dysfunction and ATP deficits exacerbating α-syn pathology (Chinta & Andersen, 2008; Rocha *et al*, 2022; Sherer *et al*, 2003).

Besides protecting neurons against oxidative stress (Volkert & Crowley, 2020), OXR1 has been implicated in retromer maintenance (Wilson *et al*, 2024) and epigenetic regulation in neurodevelopment (Lin *et al*, 2023), suggesting a broader role in protein homeostasis. *OXR1* expression is disease- and context-dependent: while it is downregulated in PD patient brains (Zhang *et al*, 2005), it is upregulated in ALS human neuronal models (Hruska-Plochan *et al*, 2024), suggesting that its function may shift depending on cellular stress conditions. Importantly, *OXR1* loss in humans is associated with severe neurological defects and premature death (Wang *et al*, 2019). Exosomal miR-137 downregulates OXR1, worsening oxidative stress in a PD mouse model (Jiang *et al*, 2019) and underscoring the critical role of OXR1 in neurodevelopment. Thus, while OXR1 activation may enhance neuronal survival at physiological levels, supraphysiological overexpression (as achieved here via CRISPRa) could overwhelm mitochondrial buffering capacity, shifting from protective to detrimental outcomes. These findings suggest that while *OXR1* activation promotes pSyn^129^ accumulation, its neuroprotective roles must be carefully considered in therapeutic strategies, as indiscriminate inhibition could have unintended consequences on neuronal function. While *OXR1* is essential for oxidative stress resistance, we found that its overexpression disrupts mitochondrial homeostasis, suppressing proton motive force genes (*ATP5F1A/B, NDUFS2, NDUFS4*) and driving pSyn^129^ accumulation. Studies suggest that excessive antioxidant activity can induce reductive stress, which in turn disrupts mitochondrial metabolism and proteostasis (Ma *et al*, 2020; Xiao & Loscalzo, 2020). Hence, reductive stress— rather than oxidative stress alone—may contribute to α-Syn pathology, potentially through impaired NAD+/NADH homeostasis, ATP depletion, or dysregulated chaperone-mediated refolding.

We observed that *OXR1* activation preferentially increases α-Syn aggregates phosphorylation (EP1536Y) in neuronal somata, suggesting that mitochondrial dysfunction exacerbates α-Syn phosphorylation in later-stage aggregates. This finding aligns with previous reports that early α-Syn aggregates initially form in neurites before redistributing to the soma, where they accumulate into larger phosphorylated inclusions (Mahul-Mellier *et al*, 2020). Although our PFF-based model reliably induces pSyn^129^ inclusions, it does not fully capture the complexity of patient-derived Lewy bodies, which often incorporate lipids and membranous organelles (Shahmoradian *et al*, 2019; Bayati *et al*, 2024). Nevertheless, seeded PFF aggregates in neuronal culture can partly recapitulate organelle and lipid incorporation, indicating some overlap with the pathology observed in patient brain tissue (Mahul-Mellier *et al*, 2020).

Antibody specificity remains a critical consideration in α-Syn aggregation studies. While the 81A antibody, a widely used pan-pSyn^129^ marker, has been reported to cross-react with phosphorylated neurofilaments in pathological inclusions (Rutherford *et al*, 2016), we observed no detectable 81A signal in untreated control conditions, confirming no off-target binding in our experimental system.

Concordant results obtained with EP1536Y—a highly selective antibody for pSer^129^ α-Syn (Lashuel *et al*, 2022)—strengthen the validity of our findings, as both antibodies consistently reflected α-Syn pathology across models.

Although Ser^129^ phosphorylation dominates PD pathology, it remains unknown whether our identified genetic modifiers also affect other phosphorylation sites (e.g., Tyr125, Ser87, and Tyr39) reported to be critical for α-Syn membrane binding (Srinivasan *et al*, 2021). The strain-specific sensitivity of MSA and fibrillar α-Syn strains to OXR1 activation suggests a structure/pathology relationship. Further structural characterization of α-Syn strains may clarify this selectivity and provide insights into differential aggregation mechanisms. Notably, MSA-derived α-Syn strains are often reported to exhibit more aggressive prion-like propagation than PD- or DLB-derived strains (Shahnawaz *et al*, 2020), potentially amplifying the mitochondrial perturbations triggered by *OXR1* activation and thereby intensifying pSyn^129^ accumulation.

The EMC complex maintains ER homeostasis, and its malfunction leads to the accumulation of misfolded proteins and unfolded protein response (UPR) activation (Shurtleff *et al*, 2018). Aberrantly folded proteins are typically cleared through ERAD or ER-to-lysosome-associated degradation (ERLAD) (Fasana *et al*, 2024). Notably, while EMC4 plays an important role in EMC functionality, its depletion does not compromise other EMC subunits’ stability or abundance, indicating that EMC4 is not structurally essential for the complex (Shurtleff *et al*, 2018). In PD, α-Syn aggregates localize to the ER in both human postmortem brains and mouse models (Colla *et al*, 2012b). α-Syn also interacts with ER chaperones, leading to impaired ER-associated proteostasis (Colla *et al*, 2012a). The finding that *EMC4* ablation reduces pSyn^129^ burden via enhanced lysosomal degradation reinforces the emerging role of ER-lysosome crosstalk in α-Syn clearance and offers a mechanistic bypass for ERAD-compromised neurons in synucleinopathies. These findings align with recent reports demonstrating the involvement of lysosomal degradation (Gao *et al*, 2025), and ER-phagy, a selective form of autophagy that degrades ER components to maintain ER homeostasis (Kim *et al*, 2023) in α-Syn aggregate removal. They are further supported by evidence that enhancing ER proteostasis and protein trafficking in patient-derived neurons can synergistically reduce α-Syn pathology (Stojkovska *et al*, 2022). Lysosomal dysfunction is intimately linked to α-Syn pathology, as exemplified by *GBA1*, which encodes the lysosomal enzyme glucocerebrosidase and constitutes one of the strongest genetic risk factors for Parkinson’s disease (Mazzulli *et al*, 2011). Loss-of-function mutations in *GBA1* impair lysosomal clearance of α-Syn, thereby exacerbating its aggregation and toxicity. Although *GBA1* was not included in our sublibrary, our identification of *EMC4* as a lysosomal regulator underscores that multiple, potentially convergent pathways can influence α-Syn proteostasis.

We found that *EMC4* depletion downregulates *DNAJC17*, a proteostasis-associated chaperone, and its ablation reduces pSyn^129^⁺ prevalence. While *DNAJC17* has been implicated in nuclear mRNA processing and splicing (Pascarella *et al*, 2018), its direct role in pSyn^129^ clearance whether as a mediator of lysosomal degradation or an epiphenomenon, remains unresolved. While our findings highlight EMC4 as a potent regulator of α-Syn pathology, existing large-scale transcriptomic and proteomic studies in PD and MSA do not explicitly report EMC4 dysregulation. However, this does not exclude the possibility of subtle or region-specific changes in EMC4 expression or function that may fall below detection thresholds in bulk analyses.

In conclusion, our work defines mitochondrial OXR1 (activation-driven enhancer) and ER-associated EMC4 (ablation-dependent suppressor) as important regulators of α-Syn proteostasis. The polygenic and multifactorial nature of synucleinopathies demands a shift toward combinatorial strategies targeting mitochondrial resilience, ER-lysosome coordination, and post-translational modification networks, some of which are being identified in the current study. By leveraging advanced models such as patient-derived fibrillar strains, human iPSC-derived neurons, and CRISPR tools to target these pathways, we may disrupt self-reinforcing cycles of proteostatic collapse in Parkinson’s disease and possibly in other synucleinopathies.

## Methods

### Generation and maintenance of HEK Cell Lines

The HEK293 QBI cell line, which stably expresses wild-type α-synuclein (α-Syn), referred to as HEK^Syn^ was used in this study and kindly provided by Prof. Kelvin C. Luk (Luk *et al*, 2009) (Kelvin=C.=Luk laboratory, University of Pennsylvania). Cells were maintained in DMEM (Catalog #31053-036, Thermo Fisher Scientific) supplemented with 10% FBS (Hyclone, Heat Inactivated, Catalog #SV30160.03HI, GE Healthcare BioSciences, Austria GmbH), 1% GlutaMax (Catalog #35050-038, Thermo Fisher Scientific), and 1% Penicillin/Streptomycin (Catalog #15070063, Thermo Fisher Scientific). To ensure the continuous expression of α-Syn, the culture medium was supplemented with Geneticin (0.4 mg/mL, Catalog #10131035, Thermo Fisher Scientific). Routine culture and expansion were performed in T75 flasks (TPP, Trasadingen, Switzerland) under standard conditions.

### Generation of CRISPR activation and CRISPR ablation cell lines

To establish CRISPR-compatible cell lines, HEK293 QBI α-Syn (referred as HEK^Syn^) cells were transfected with dCas9-VPR or Cas9. The dCas9-VPR plasmid (pXPR_120, Catalog #96917, Addgene) was introduced using Lipofectamine 2000 (Catalog #11668027, Thermo Fisher Scientific), following the manufacturer’s protocol. After transfection, cells were subjected to antibiotic selection with Blasticidin S HCl (10 μg/mL, Catalog #A1113903, Thermo Fisher Scientific) to establish a stable dCas9-expressing cell population.

Similarly, Cas9-expressing cells were generated via lentiviral transduction using lentiCas9-Blast (Catalog #52962, Addgene), followed by Blasticidin selection. To maintain the genetic modifications, the culture medium for dCas9 and Cas9-expressing cells was supplemented with Blasticidin (10 μg/mL) alongside Geneticin (0.4 mg/mL).

For monoclonal cell line generation, polyclonal CRISPR cell lines underwent limiting dilution in 96-well flat-bottom plates (Catalog #7000209, TPP92096, TPP) to isolate single-cell clones. Wells containing a single viable cell were monitored for colony formation, and successfully expanded clones were validated for Cas9 or dCas9-VPR expression via molecular and phenotypic assays. The Cas9-expressing Cl-7-Cas9-aSyn HEK and dCas9-VPR-expressing Mo-4-dCas9-aSyn HEK clones were designated as the CRISPRa and CRISPRo cell lines, respectively, in accordance with the experimental workflow.

Cells were passaged at 80–90% confluency using 0.25% Trypsin-EDTA (Catalog #25200056, Thermo Fisher Scientific) and cryopreserved in Bambanker™ (Catalog #BB01, LuBio Science). Cryovials were gradually cooled to -80°C before long-term storage in liquid nitrogen. Mycoplasma testing was routinely conducted using the LookOut® Mycoplasma PCR Detection Kit (Catalog #MP0035, Sigma-Aldrich) according to the manufacturer’s instructions.

### Transfection efficiency

Transfections were performed using ViaFect Transfection Reagent (Catalog #E4981, Promega), which is optimized for HEK cells. Plasmid DNA encoding BFP (T.gonfio) or a GFP plasmid of similar size (Addgene Plasmid #48138) was mixed with ViaFect in Opti-MEM™ (Catalog # 31985062, Thermo Fisher) following the manufacturer’s instructions. Forty-eight hours post-transfection, efficiency was evaluated by quantifying the percentage of GFP- or BFP-positive cells using fluorescence microscopy and flow cytometry.

### Expression and purification of recombinant α-Syn

Human wt full-length α-Syn (NM_000345) was cloned into the ampicillin-resistant bacterial expression vector pRK172 and transformed into competent BL21(DE3) RIL E. coli cells. Transformed bacteria were plated on LB/Amp agar plates and incubated overnight at 37°C. For preculture preparation, a single colony from the overnight plate was transferred to SOC medium (Catalog #15544034, Invitrogen™) and incubated for 6 hours at 37°C, 200 rpm. The preculture was then used to inoculate Terrific Broth (TB) medium supplemented with ampicillin in baffled flasks, followed by overnight incubation at 37°C with shaking to induce protein expression.

For cell harvesting, overnight cultures were transferred into centrifuge bottles, pelleted by centrifugation, and the supernatant was discarded. Cell pellets were stored at -20°C until further processing. For α-Syn purification, the pellets were thawed and lysed via freeze-thaw cycles using liquid N₂ and tap warm water. The lysates were cleared by centrifugation, and the supernatant was treated with streptomycin sulfate to precipitate DNA, followed by ammonium sulfate precipitation to isolate α-Syn. The pellet was resuspended and dialyzed against ion exchange chromatography (IEC) start buffer for desalting.

Purification was performed in two steps: Anion exchange chromatography (AEC) for initial purification and Size exclusion chromatography (SEC) to achieve high purity and monomeric α-Syn. The purified α-Syn was concentrated to ∼700 µM (∼10 mg/mL) using Amicon Ultra centrifugal filter devices (10 kDa MWCO, Catalog #UFC503024, Millipore). The protein was aliquoted into 250 µL volumes and stored at -20°C. All steps were performed at room temperature following BSL-2 biosafety guidelines.

### Preparation and sonication of α-Syn preformed fibrils

Purified monomeric α-Syn was diluted to 345 µM (4.98 mg/mL) in PBS (300 µL/aliquot) and incubated in screw-cap tubes at 60°C with continuous agitation (1,000 rpm, 72 hours) in a Thermomixer with a heated lid to induce fibrillation. Fibrillation progression was monitored at 24, 48, and 72 hours by centrifuging 20 µL samples (15,200 rpm, 60 minutes, 25°C), separating supernatants and pellets for storage at -20°C. Supernatants and pellets were analyzed via SDS-PAGE (12% Bis-Tris gel, MOPS buffer) after denaturation (95°C, 10 minutes) and Coomassie staining (Instant Blue) to quantify α-Syn. Aliquoted preformed fibrils (PFFs) were stored at -80°C to preserve integrity. For experiments, PFFs were thawed, diluted to 0.5 mg/mL (monomer equivalent) in PBS, and sonicated (10-minute cycles: 30 seconds on/30 seconds off) using a high-powered water bath sonicator filled with ice-cooled water to ensure consistent fragmentation. This standardized protocol was consistently followed to ensure reproducibility across all experimental runs.

### Characterization by Transmission Electron Microscopy (TEM)

α-Syn monomers, PFFs, and sonicated PFF ultrastructures were analyzed by TEM. Samples were diluted to 0.5–1 mg/mL in PBS (pH 7.4), applied to mesh copper grids for 1–2 minutes, and excess liquid blotted with filter paper. Grids were negatively stained with 2% uranyl acetate (Electron Microscopy Sciences, Catalog number: 22400-2) for 30 seconds, rinsed three times with double-distilled water, and air-dried at room temperature. TEM imaging was performed on a FEI Talos 120 TEM (Thermo Fisher Scientific) at the Center for Microscopy and Image Analysis (University of Zurich), using magnifications of 6,000× to 120,000× to capture an overview and structural features of α-Syn fibrils.

### Thioflavin T (ThT) Fluorescence Assay

Fibril formation was monitored using ThT (Sigma-Aldrich, Catalog number: T3516). A 10 mM ThT stock was prepared in ultrapure water, filtered (0.22 μm syringe filter), and diluted to 25 μM in PBS. For each time point (0, 1, 3, 12, 24, 48, 72 hours), 5 μL of fibril suspension was added to 195 μL ThT solution in a black-walled 96-well plate (Greiner Bio-One, Catalog number: 655209). After 5 minutes of incubation (25°C), fluorescence was measured using a FLUOstar Omega Microplate Reader (BMG Labtech) at 450 nm excitation/480 nm emission.

### Preparation of fibrillar and patient-derived fibrils α-Syn strains

The generation of the two α-Syn fibrillar strains (fibrils and ribbons, 350 μM) (Bousset *et al*, 2013), and patient-derived strains (PD, DLB, MSA; 100 μM) (Van Der Perren *et al*, 2020), their fragmentation and storage at -80°C have been extensively described. For use, aliquots were thawed in a 37°C water bath (3 minutes), equilibrated to room temperature, and diluted to 0.5 mg/mL in PBS. No refreezing was permitted to preserve fibril integrity.

### LT1-mediated PFF delivery

Sonicated PFFs (0.5 mg/mL) were complexed with TransIT™-LT1 Transfection Reagent (Catalog #MIR 2306, Mirus Bio) at a 1:3 (v / v) ratio in Opti-MEM™ (Thermo Fisher Scientific, Catalog number: 31985070) and incubated for 15 minutes (25°C). Cells were washed with PBS, treated with the PFF-LT1 complex (final PFF concentration: 7.5 µg/mL), and maintained in DMEM complete medium (without penicillin/streptomycin).

### CRISPR activation and ablation primary screen

The screening assay was conducted in a high-throughput 384-well format. CRISPRa cells (3,000 cells/well) were seeded in PDL-coated 384-well PhenoPlates (Catalog #6057300, PerkinElmer) using the BioTek MultiFlo FX Multimode Dispenser (Agilent). Transfection was performed using a custom genome-wide CRISPRa library (T.gonfio library; 2,428 genes related to mitochondria, trafficking, and motility, with four non-overlapping guide RNAs [qgRNAs] per gene) (Yin *et al*, 2024).

Each well received 60 ng plasmid DNA complexed with 0.3 µL ViaFect Transfection Reagent (Promega, #E4981) in Opti-MEM™ I Reduced Serum Medium (Thermo Fisher Scientific, #31985070). Non-targeting (scrambled) qgRNAs and *RAB13-*positive controls were included. Post-transfection, cells underwent 48-hour puromycin selection (0.6 µg/mL, Thermo Fisher Scientific, #A1113803) before treatment with 7.5 µg/mL sonicated PFFs complexed with TransIT™-LT1 Transfection Reagent (Mirus Bio, #MIR2306). After 72 hours, cells were fixed and immunostained for phosphorylated α-Syn at Ser^129^ (pSyn^129^). Imaging was conducted using the GE IN Cell Analyzer 2500HS high-content microscopy system.

For ablation screen, CRISPRo cells (2,000 cells/well) were plated and transfected with a custom CRISPRo library (T.spiezzo library; 2,304 genes, 4 gRNAs per gene). Although alternative transcription start sites can sometimes inflate CRISPRa libraries, in this study the difference in library size mainly reflected sublibrary arrangement, with some “tail genes” continuing mid-plate from previous collections, ensuring complete coverage but resulting in a slightly larger CRISPRa library. Non-targeting controls and *PIKFYVE*-positive controls were included. The transfection and PFF treatment protocols mirrored CRISPRa, but puromycin selection lasted 96 hours to ensure stable ablation. pSyn^129^ inclusion analysis followed the same immunostaining, imaging, and analysis workflows. Imaging was conducted using the ImageXpress Confocal HT.ai (Molecular Devices). Both screenings were performed in duplicates, with each sample plated in two independent wells to ensure technical reproducibility.

### iPSCs culture and maintenance

The maintenance of iPSCs, neuronal differentiation, and maturation followed the protocols described here (Fernandopulle et al, 2018) and Tian et al (Tian et al, 2021). WTC11 human iPSCs harbouring a trimethoprim (TMP)-inducible CRISPR activation system (DHFR–dCas9–VPH) were kindly provided by the Kampmann Laboratory (University of California, San Francisco). iPSCs were cultured on Matrigel-coated (Corning® Matrigel® hESC-Qualified Matrix, LDEV-free, Catalog #54277, Corning) cell culture dishes with daily media changes in STEM Flex Medium (StemFlex™ Medium, Catalog #A3349401, Thermo Fisher Scientific). Additionally, a fresh 10 µM ROCK inhibitor (Y-27632, Catalog #1254/10, Tocris Bioscience) was added to the medium. Cells were dissociated using Accutase (StemPro® Accutase® Cell Dissociation Reagent, Catalog #A1110501, Thermo Fisher Scientific). For long-term storage, cells were cryopreserved in CryoStor® (Catalog #C2874, Sigma-Aldrich). Routine mycoplasma testing was performed to ensure culture integrity. CRISPRa guide RNAs were introduced into iPSCs using lentiviral transduction at an MOI of 0.3, followed by puromycin selection to generate CRISPRa Non-targeting and CRISPRa OXR1 lines before differentiation.

### Differentiation and culture of iPSCs into i3 cortical neurons

The iPSCs were differentiated into glutamatergic (i3) cortical neurons in Matrigel-coated dishes using an induction medium composed of DMEM/F-12, HEPES (Catalog #31330095, Invitrogen), supplemented with N2 supplement (N2 Supplement, 100X, Catalog #17502-048, Invitrogen), non-essential amino acids (NEAA MEM, Catalog #10370047, Thermo Fisher Scientific), and GlutaMAX Supplement 100X (Gluta-MAX MEM, Catalog #41090028, Thermo Fisher Scientific). Additionally, 2 μg/mL doxycycline (Catalog # D9891-10G, Sigma) was freshly added to the medium daily.

Differentiated neurons at day 3 *in vitro* (DIV 03) were further matured in a neuronal maturation medium. Culture plates were coated with Poly-D-Lysine (Catalog #A3890401, Thermo Fisher Scientific), Poly-L-Ornithine (Catalog #P4957, Sigma-Aldrich), and mouse Laminin (Natural, Catalog #23017015, Thermo Fisher Scientific). The maturation medium consisted of BrainPhys™ Neuronal Medium (Catalog #05790, StemCell Technologies), supplemented with recombinant human NT-3 (Neurotrophin-3, Catalog #450-03, Peprotech), recombinant human/murine/rat BDNF (Brain-Derived Neurotrophic Factor, Catalog #450-02, Peprotech), serum-free B-27™ Supplement 50X (Catalog #A3582801, Thermo Fisher Scientific), and 100 nM TMP (Trimethoprim, Catalog #195527, Biomedicals Europe) for dCas9-VPH activation.

At DIV04, iPSC-derived cortical neurons were treated with sonicated preformed fibrils PFFs at a concentration of 7.5 μg/mL, utilizing half-conditioned medium to minimize medium turnover effects. The following day, the medium was replaced, and every 2–3 days, iPSC-derived cortical neurons were replenished with TMP-containing neuronal maturation medium for a total duration of 4 weeks. For the shRNA based knockdown experiment, the EMC4 human shRNA Plasmid Kit and a 29-mer scrambled shRNA cassette (non-targeting control) (Catalog #TL300940, OriGene) were introduced using lentiviral transduction at a multiplicity of infection (MOI) of 2 on Day 4 *in vitro* (DIV 04). After 24 hours, the medium was fully replaced with half-conditioned media with PFFs.

### Differentiation of iPSCs into iDA neurons

iPSCs were differentiated into induced dopaminergic (iDA) neurons using an adapted protocol from Sheta et al., 2023 (Sheta *et al*, 2023). Cells were plated on Matrigel-coated dishes and maintained in induction medium based on DMEM/F-12 with HEPES **(Table S5).** The differentiation medium was freshly prepared and applied, and cells were left undisturbed for the first three days (DIV 0–3) to facilitate initial neuronal induction. On DIV 3, differentiated iPSCs were detached and replated onto Poly-L-Ornithine (PLO)- and Laminin-coated plates in Neurobasal-based PreDOPA medium **(Table S5)** to initiate dopaminergic differentiation. Cells were maintained in PreDOPA medium for three days (DIV 3–6) without disturbance to allow cell attachment and early differentiation into the dopaminergic lineage. Following the PreDOPA stage (DIV 6 onward), cells were transitioned to iDA dopaminergic neuron media, which was based on Neurobasal Medium **(Table S5).** To maintain optimal neuronal differentiation conditions, a half-media change was performed every 5 days.

### iDA neuronal culture and PFF treatment assay

Neurons were treated with sonicated α-Syn PFFs after 5 days in iDA dopaminergic neuron media at a concentration of 7.5 μg/mL, utilizing half-conditioned medium. The following day, the medium was replaced, and every 5 days, neurons were replenished with TMP-containing neuronal maturation medium for a duration of 4 weeks. Imaging was performed using the Operetta CLS High-Content Analysis System (PerkinElmer). Image analysis was conducted using the CellProfiler pipeline as previously described. To account for batch-to-batch variability of PFFs, data from different PFF batches were normalized to the corresponding non-targeting control (NTG) conditions, enabling conversion into effect size for comparative analysis.

### Immunostaining

Cells were washed once with Tris-Buffered Saline (TBS) and fixed with 4% paraformaldehyde (PFA, methanol-free) (Thermo Fisher Scientific, Catalog #043368.9M) for 15 minutes at room temperature (RT). After three washes with TBS, cells were permeabilized with 0.1% Triton X-100 (Catalog #9036-19-5, Sigma) in TBS for 10 minutes, followed by another three washes with TBS. To minimize non-specific binding, cells were incubated for 1 hour at RT in 4% bovine serum albumin (BSA) in TBS. For primary antibody staining, cells were incubated overnight at 4°C with anti-α-Syn (phospho S^129^; clone 81A) [81A] (Catalog #825701, BioLegend) diluted 1:2500 in 0.5% BSA/TBS. The following day, after three washes with TBS, cells were incubated for 1 hour at RT with the secondary antibody, diluted 1:400 in 0.5% BSA/TBS. For whole-cell labeling, HCS Cell Mask™ Deep Red Stain (Catalog #H32721, Thermo Fisher Scientific) was diluted 1:5000 in PBS, and nuclei were counterstained with DAPI (1:10,000 dilution, Sigma-Aldrich, Catalog #D9542) for 10 minutes. All washing steps were performed using the BioTek 405 TS Washer (Agilent) to ensure consistency across experiments. Unless otherwise specified, 81A was used for pSyn^129^ immunostaining. For experiments requiring alternative validation, phospho-α-Syn (pS^129^) antibody [EP1536Y] (Catalog #ab51253, Abcam) was employed. Additional antibodies and their specific dilutions are listed in **Table S6.**

### Imaging and Image analysis

Image acquisition was performed using a GE IN Cell Analyzer 2500HS widefield system (10X or 20X or 40X objectives) or an ImageXpress Confocal HT.ai (Molecular Devices; 10X or 20X objective). For nuclear and pSyn^129^ signal segmentation, ilastik 1.4.0 was used, employing a random forest classifier algorithm trained manually on raw images. Probability maps generated in ilastik were exported to CellProfiler 4.2.1 for further analysis. Cell segmentation was performed using the Propagation algorithm with the nucleus as a reference, while object segmentation was achieved using the minimum cross-entropy algorithm in CellProfiler. Quantitative assessments included total cell counts, the percentage of pSyn^129+^ cells, total pSyn^129^ spot area and total pSyn^129^ signal intensity. Raw images were used for pSyn^129^ spot intensity calculations, without applying pretrained ilastik models. For confocal 3D imaging, a FluoView FV10i Confocal Laser Scanning Microscope (Olympus) was used. Images were adjusted uniformly, and pseudo-coloring was applied to generate representative images using ImageJ.

### Neuronal Image analysis

Neuronal image analysis was conducted using CellProfiler to quantify the pSyn^129^ spots in the neurite and soma regions separately. Segmentation was applied to delineate these compartments. Feature enhancement and suppression techniques were applied to refine segmentation and associate pSyn^129^ spots with their respective neuronal compartments. The detected pSyn^129^ signal was normalized to the MAP2-positive area to account for morphological variations. pSyn^129^ spots outside defined neurite and soma regions were excluded to maintain specificity.

### Labeling and Treatment of Fibrils

Pre-formed fibrils (PFFs) were labeled with Alexa Fluor™ 488 (Alexa Fluor™ 488 NHS Ester Succinimidyl Ester, Catalog #A20000, ThermoFisher Scientific) following the manufacturer’s protocol. PFFs (2 mg/mL) were sonicated and incubated with Alexa Fluor™ 488 for 1 hour at room temperature with continuous shaking at 300 rpm. Labeled fibrils were then transferred to a dialysis membrane (Slide-A-Lyzer™ MINI Dialysis Device, 10K MWCO, 0.1 mL, Catalog #69570, ThermoFisher Scientific) and dialyzed against PBS with magnetic stirring. The PBS buffer was replaced three times over 24 hours to remove unbound dye. After dialysis, labeled PFFs were collected, aliquoted, and stored at -80°C until use. For neuronal uptake experiments, fluorescently labeled α-Syn PFFs were incubated with neurons for 6 hours, and uptake was quantified using flow cytometry.

### Lentiviral production and titration

HEK293T cells were plated on poly-D-lysine (PDL)-coated dishes to enhance cell adherence and growth. Once cells reached 50–60% confluency, they were transfected using Lipofectamine 3000 (Catalog #L3000008, Thermo Fisher) following the manufacturer’s protocol. The transfection cocktail included transfer plasmids carrying CRISPR guides, along with packaging plasmids: psPAX2 (Addgene #12260) and VSV-G (Addgene #8454) to facilitate lentiviral particle production. Six hours post-transfection, the medium was replaced with Virus Harvesting Medium, composed of DMEM supplemented with 10% fetal bovine serum (FBS) and 10 mg/mL bovine serum albumin (BSA) to optimize viral particle collection. The viral supernatant was carefully harvested and stored appropriately. For viral titration, target cells were seeded at 2.5 × 10= cells per well in PDL-coated 24-well plates, followed by the addition of serial dilutions of the harvested virus. Three days post-infection, intracellular flow cytometry staining and analysis was performed to determine the transduction efficiency and quantify the viral titre. The titre, expressed as transducing units per millilitre (TU/mL), was calculated based on the proportion of fluorescent-positive cells.

### Intracellular flow cytometry staining

Cells were trypsinized, pelleted, and washed with TBS before fixation and permeabilization using Cyto-Fast™ Fix/Perm Buffer (Catalogue #426803, BioLegend) for 20 minutes at room temperature (RT). Following fixation, cells were washed twice with 1X Cyto-Fast™ Perm Wash solution before incubation with Alexa Fluor® 594 81A antibody (Catalogue #825708, BioLegend) for 20 minutes in the dark at RT. After staining, cells were washed again with 1X Cyto-Fast™ Perm Wash solution, followed by a final wash with Cell Staining Buffer (Catalogue #420201, BioLegend). The cells were then resuspended in Cell Staining Buffer and acquired on an LSR II Fortessa 4L flow cytometer for the quantification of intracellular α-Syn phosphorylation (pSyn^129^).

For data analysis, flow cytometry data were exported and analyzed in FlowJo v10.1. Gating was performed to exclude debris and doublets, ensuring an accurate representation of single, viable cells. CRISPR-expressing cells were identified by BFP positivity (qgRNA expressing cells), and within this population, the percentage of Alexa Fluor® 594-positive cells was quantified as a measure of 81A positive cells.

### Gene expression analysis using quantitative Real-Time PCR (qRT-PCR)

Gene expression levels were analyzed using quantitative real-time PCR (qRT-PCR). Total RNA was extracted using TRIzol™ Reagent (Catalog #15596018, Thermo Fisher Scientific) or the RNeasy Mini Kit (Catalog #74104, Qiagen), following manufacturer protocols. RNA purity and concentration were assessed using a NanoDrop spectrophotometer (Catalog #ND-1000, Thermo Fisher Scientific). cDNA synthesis was performed with the QuantiTect reverse transcription kit (Catalog #205311, Qiagen) using 1 µg of RNA per reaction. qRT-PCR was carried out with the FastStart Universal SYBR Green Master Mix (Catalog #491385000, Sigma-Aldrich) on a ViiA 7 Real-Time PCR System (Catalog #4453536, Thermo Fisher Scientific) under the following thermal cycling conditions: initial denaturation at 95°C for 10 minutes, followed by 40 cycles of 95°C for 15 seconds and 60°C for 1 minute, with a post-amplification melting curve analysis to verify amplicon specificity. Relative gene expression was quantified using the 2^−ΔΔCT method, normalized to the housekeeping gene β-actin, and compared to control conditions. Primer sequences are listed in **Table S7.**

### Western Blot

Cells were lysed using either lysis buffer (50 mM Tris pH 8, 150 mM NaCl, 1% Triton^®^ X-100) or RIPA buffer (RIPA 10X, Catalog #9806S, Cell Signaling Technology), supplemented with protease inhibitors (Complete EDTA-free Protease Inhibitor Cocktail, Catalog #11873580001, Sigma-Aldrich) and phosphatase inhibitors (PhosSTOP, Catalog #4906845001, Roche). Protein concentrations were determined using the Bicinchoninic Acid (BCA) Protein Assay (Pierce™ BCA Protein Assay Kit, Catalog #A55864, Thermo Scientific™) according to the manufacturer’s instructions. Lysates were resolved by SDS-PAGE and transferred onto PVDF membranes (iBlot™ 2 Transfer Stacks, PVDF, Catalog #IB24002, Thermo Scientific™) using the Invitrogen™ iBlot™ 2 Gel Transfer Device. Membranes were blocked with 5% SureBlock (Catalog #SB232010-500G, LubioScience) in PBST (0.1% Tween, Catalog #9005-64-5, Sigma, in 1X PBS) for 1 hour at room temperature. Primary antibody incubation was performed overnight at 4°C with shaking at optimized dilutions, followed by three 10-minute washes in PBST (PBS + 0.2% Tween). All the antibodies used for western blot with specific dilutions are listed in **Table S6.**

For secondary detection, membranes were incubated for 45 minutes at room temperature with HRP-conjugated respective secondary antibodies at 1:10,000 dilution. After three 10-minute washes in PBST, signals were developed using Classico, Crescendo, or Forte ECL solutions (Immobilon Western HRP substrate, Merck Millipore) depending on signal intensity. β-actin was used as a loading control, where membranes were blocked for 1 hour, followed by a 30-minute incubation with β-actin HRP-conjugated antibody (anti-actin HRP, Catalog #A3854-200UL, Sigma-Aldrich). Signal acquisition was performed immediately after three 10-minute washes in PBST.

### Co-Immunoprecipitation (Co-IP)

CRISPRo HEK cells were washed with PBS and lysed in CHAPS lysis buffer (25 mM Tris, 150 mM NaCl, 1 mM EDTA, 1% CHAPS, 5% glycerol, pH 7.4) supplemented with protease and phosphatase inhibitors. Protein concentration was determined using the BCA assay, and 1 mg of total lysate was diluted in CHAPS lysis buffer to a final volume of 500 μL per tube. For immunoprecipitation, Recombinant Anti-EMC4 antibody (Catalogue #ab184544, Abcam) was conjugated to Dynabeads™ Protein G (Catalogue #10003D, Thermo Fisher Scientific) for 1 hour at 4°C on a spinning wheel at 30 rpm. Normal Rabbit IgG (Catalogue #12-370, Merck) was used as an isotype control. Lysates were pre-cleared (Input Lysate) to remove nonspecific binding, then incubated overnight at 4°C with antibody-conjugated beads under constant rotation (30 rpm). After incubation, tubes were placed on a magnetic rack, and the cleared supernatant (flow-through, FT) was collected. The supernatant from the EMC4 IP was referred to as EMC4 FT, and the flow-through from the IgG control was referred to as IgG FT. Beads were then washed four times with 200 μL of CHAPS lysis buffer per tube to remove unbound proteins. Immunoprecipitated proteins were eluted in 20 μL of 2X SDS Loading Buffer (LB) with DTT (DL-Dithiothreitol, Catalog #10708984001) and subjected to SDS-PAGE. After electrophoresis, proteins were transferred onto PVDF membranes and immunoblotted with mouse anti-α-Syn antibody (Syn 211, Catalogue #AHB0261, Thermo Fisher Scientific). Membranes were subsequently stripped and re-probed for EMC4 to confirm successful immunoprecipitation.

### Lactate dehydrogenase (LDH) Release Assay

Lactate dehydrogenase (LDH) release was measured using the LDH-Glo™ Cytotoxicity Assay (Catalog #J2380, Promega) according to the manufacturer’s instructions. LDH storage buffer was freshly prepared at a final concentration of 200 mM Tris-HCl (pH 7.3), 10% Glycerol, and 1% BSA, stored at 4°C, and used for diluting samples. Following experimental treatments, 5 µL of supernatant was carefully collected from each well to avoid disturbing adherent cells and transferred to a white-walled opaque plate. To determine maximum LDH release, a subset of wells was treated with 0.2% Triton X-100, serving as a positive control for complete cell lysis. Samples were diluted in LDH storage buffer. LDH detection enzyme mix was combined with the Reductase Substrate and thoroughly mixed with the diluted samples in LDH buffer (1:1) in an opaque plate. Plates were incubated at room temperature in the dark for 60 minutes, after which luminescence was measured using an EnVision plate reader (PerkinElmer). LDH release values were normalized and expressed as relative cytotoxicity.

### MitoSOX-based detection of mitochondrial ROS

Mitochondrial reactive oxygen species (ROS) were measured using MitoSOX™ (Catalog #M36007, Thermo Fisher Scientific). As a positive control, cells were treated with 2 µM rotenone (Catalog #R8875-1G, Sigma-Aldrich) for 2 hours to induce ROS. A 5mM MitoSOX stock solution was prepared by dissolving 13 µL of anhydrous DMSO into the supplied aliquot. The working solution was prepared by diluting MitoSOX to a final concentration of 1 µM in colorless Opti-MEM medium. Cells were incubated with MitoSOX and Hoechst 33342 (Catalog #62249, Thermo Fisher Scientific) for 30 minutes at room temperature (RT), protected from light. After incubation, cells were gently washed 3× with pre-warmed medium. Live-cell imaging was performed at 20× or 40× magnification using the GE IN Cell Analyzer.

### Mitochondrial membrane potential measurement

Mitochondrial membrane potential (ΔΨm) was assessed using Image-iT™ TMRM Reagent (Catalog #I34361, Thermo Fisher Scientific). Nuclear labeling was performed using Hoechst 33342 (Catalog #62249, Thermo Fisher Scientific) at a 1:2000 dilution from a 10 mg/mL stock solution. Cells were incubated with 50 nM TMRM for 30 minutes at 37°C to establish baseline ΔΨm. Regions of interest (ROI) were selected to ensure coverage of at least 1,000 cells per condition. Following baseline measurement: Oligomycin (2 µM, Catalog #ab141829, Abcam) was added to inhibit ATP synthase (Complex V), initially increasing ΔΨm due to reduced proton leakage. TMRM fluorescence was recorded every 3 minutes for 15 minutes to monitor changes in mitochondrial polarization. FCCP (2 µM, Catalog #ab120081, Abcam) was then added to collapse ΔΨm, inducing complete mitochondrial depolarization. TMRM fluorescence was measured every 3 minutes for an additional 15 minutes to assess mitochondrial uncoupling. Images were consistently captured from the same ROI throughout the experiment. ΔΨm changes were analyzed by comparing TMRM fluorescence intensity before and after treatments to quantify mitochondrial depolarization.

### Lysosomal and proteasomal inhibition

MG132 (Catalog #C2211, Sigma-Aldrich, stock solution 5 mM) and Bafilomycin A1 (BafA1, Catalog #BML-CM110, Enzo Life Sciences, stock solution 1 mM) were prepared in DMSO and stored at – 20°C. For proteasomal inhibition, cells were treated with MG132 (5 µM) for 48 hours, with media replenished every 24 hours. For lysosomal inhibition, cells were treated with BafA1 (50 nM) for 18 hours following α-Syn PFF addition. Equivalent volumes of DMSO were used as vehicle controls. After treatments, cells were either lysed for Western blot analysis or fixed for immunostaining and imaging.

### Bulk RNA sequencing

Total RNA was extracted using RNeasy Mini Kit following the manufacturer’s protocol. RNA integrity and purity were assessed using a Qubit® Fluorometer (Life Technologies) and a Fragment Analyzer (Agilent). Samples with a 260/280 nm ratio between 1.8 and 2.1 and a 28S/18S ratio between 1.5 and 2 were considered suitable for sequencing. RNA samples (100–1000 ng) underwent poly(A) enrichment and were reverse transcribed into double-stranded cDNA using the TruSeq Stranded mRNA Kit (Illumina).

cDNA libraries were enzymatically fragmented, end-repaired, adenylated, and ligated with TruSeq adapters containing unique dual indices (UDI). Adapter-ligated fragments were enriched by PCR amplification, and final libraries were assessed for quality and quantity using a Qubit® Fluorometer and Fragment Analyzer. The final product had an average fragment size of ∼260 bp and was normalized to 10 nM in Tris-Cl (10 mM, pH 8.5, 0.1% Tween-20) before sequencing. Sequencing was performed on an Illumina NovaSeqX platform using a paired-end 2 × 150 bp strategy. Raw reads were pre-processed using fastp (Chen *et al*, 2018) with the following parameters: *-- trim_front1 1 --cut_tail 20 --trim_poly_x --poly_x_min_len 10 --length_required 18*.

Transcript quantification was performed with kallisto using the GRCh38.p13 reference transcriptome (Annotation Release 42, downloaded on 2023-01-30, doi:10.1038/nbt.3519). Differential gene expression analysis was conducted using EdgeR (Robinson *et al*, 2010). All computational analyses, including Gene Set Enrichment Analysis (GSEA) and Over-Representation Analysis (ORA), were executed via the SUSHI platform at the Functional Genomics Center Zurich (Hatakeyama *et al*, 2016).

### Statistical analysis

Statistical tests were performed using R Studio (R). For primary screening data, moderated t-tests (empirical Bayes) were performed using the limma package in R. For correlation test, data were initially evaluated for normality using the Shapiro–Wilk test (α = 0.05). Data that passed the normality assumption were analyzed with Pearson’s correlation and data violating normality assumptions Spearman’s correlation was used. For multiple comparisons, one-way ANOVA followed by Dunnett’s post hoc test was applied, while two-tailed Student’s t-test or Welch’s t-test (for unequal variance) was used for two-group comparisons. All experiments included at least three biological replicates, with 3–4 technical replicates per condition. iPSC-derived neuronal experiments were based on three independent differentiations. Detailed statistical tests and significance values are reported in the figure legends.

### Data availability

All data supporting the findings of this study are included in this published article and its Supplementary Information files. Raw microscopy images generated during this study have been deposited in FigShare (https://figshare.com/s/ce7273dfd93096f9c01e; will be made public upon publication) and Zenodo (DOI: 10.5281/zenodo.15358052). Bulk RNA sequencing data will be available through the Gene Expression Omnibus (GEO) under accession number GSE295558 upon publication, in accordance with NIH data sharing policies. Source data are provided with this paper.

### Code availability

This study did not generate novel computational code. Data analysis and visualization (e.g., statistical tests, violin plots, volcano plots) were performed using standard functions in R (version 4.3.1) with publicly available packages (e.g., ggplot2, dplyr). The CellProfiler pipeline used to analyze pSyn^129^ staining in HEK cells, and neurites, and soma is available from the corresponding author upon reasonable request.

## Acknowledgments

We thank Prof. Martin Kampmann for providing iPSCs-dCas9 VPH and Prof. Kelvin C. Luk for HEK cells. We acknowledge Araneya Sivanantharajah for technical assistance in the primary screening. We thank Rafaela Ribeiro for performing Western blotting for lysosomal inhibition studies. We thank Dr. Kathi Ging, Dr. Chiara Trevisan, and Dr. Lukas Frick for discussions on primary screen setup and data analysis. We are grateful to Johannes Riemann from the Centre for Microscopy and Image Analysis, University of Zurich, for TEM imaging of fibrils, Dr. Elif Köksal for assistance with synuclein monomer purification, Dr. Tibor Hortobagyi for discussions on hit validation, Giovanni Mariutti, Dr. Simone Hornemann, Dr. Andrea Armani, and Dr. Davide Caredio for general discussions, Federico Baroni, Ilan Margalith, Schwarz Petra and Dr. Athena Economides for technical support and discussions, Roxanne Larivière for iDA-related discussions and inputs, Dr. Maria Domenica Moccia, Dr. Hubert Rehrauer, and Catharine Aquino (Functional Genomics Center Zurich) for preparing sequencing libraries, bulk RNA sequencing, quality control, and RNA-seq data analysis. Schematic figures were created using BioRender.com. A.A. is supported by a Distinguished Scientist Award of the NOMIS Foundation and grants from the GELU Foundation, the Swiss National Science Foundation (SNSF grant ID 179040, grant ID 207872, grant ID 227951, Sinergia grant ID 183563), the Human Frontiers Science Program (grant ID RGP0001/2022), the Michael J. Fox Foundation (grant ID MJFF-024255), the CJD Foundation, and a donation from the estate of Dr. Hans Salvisberg. E.A.F. is supported by grants from The Michael J. Fox Foundation for Parkinson’s Research (MJFF), the Canadian Institutes of Health (CIHR) grant (PJT-195804) and a Canada Research Chair (Tier 1) in Parkinson Disease.

## Author Contributions

**S.N.** designed, performed or contributed to all experiments, analyzed data (including statistical, image analyses, data visualisation), and wrote the manuscript. **L.N.** performed co-immunoprecipitation (Co-IP) experiments, and αSyn uptake assays in iPSC-derived neurons; assisted in cell culture, immunostaining and imaging. **L.M.** contributed to intracellular flow cytometry-based screening of RNA-seq DEGs, lentivirus production, CTG viability assays, and Western blotting. **T.G.** designed dopaminergic neuron experiments, performed assays in dopaminergic neuronal models, and provided technical expertise. **N.K.** assisted in imaging and analysis of induced dopaminergic (iDA) cultures. **S.S.** maintained iPSC cultures and assisted in differentiation protocols. **V.B.** advised on flow cytometry experimental design and data interpretation. **J.-A.Y.** provided CRISPR guide RNAs and CRISPR-related technical guidance. **R.M.** prepared distinct α-synuclein (αSyn) fibril strains and gave inputs on fibrillar strain experimental inputs. **E.A.F.** provided critical insights into iDA related assays. **E.D.C.** provided supervision, technical expertise, experimental design, and reviewed/edited the manuscript. **A.A.** conceived, initiated and supervised the study, secured funding, coordinated the research team, and reviewed/edited the manuscript.

All authors reviewed and approved the final manuscript.

## Supplemental Information

Supplementary Figures S1-S12

Table S1. Excel spreadsheet listing CRISPRa screen data (genename, pvalue, log2FC, Cell number threshold)

Table S2. Excel spreadsheet listing CRISPRo screen data (genename, pvalue, log2FC, Cell number threshold)

Table S3. Excel spreadsheet of RNAseq data: NTG vs OXR1

Table S4. Excel spreadsheet of RNAseq data: EMC4 vs OXR1

Table S5. Excel spreadsheet listing iDA dopaminergic neurons media components

Table S6. Excel spreadsheet listing antibodies application and dilution

Table S7: Excel file containing qPCR Primers sequence

Document S1: Uncut western blot

Document S2: Excel File Source Data

