## Supplementary Figures S1-S12 for "Large-scale bidirectional arrayed genetic screens identify *OXR1* and *EMC4* as modifiers of α-synuclein aggregation"

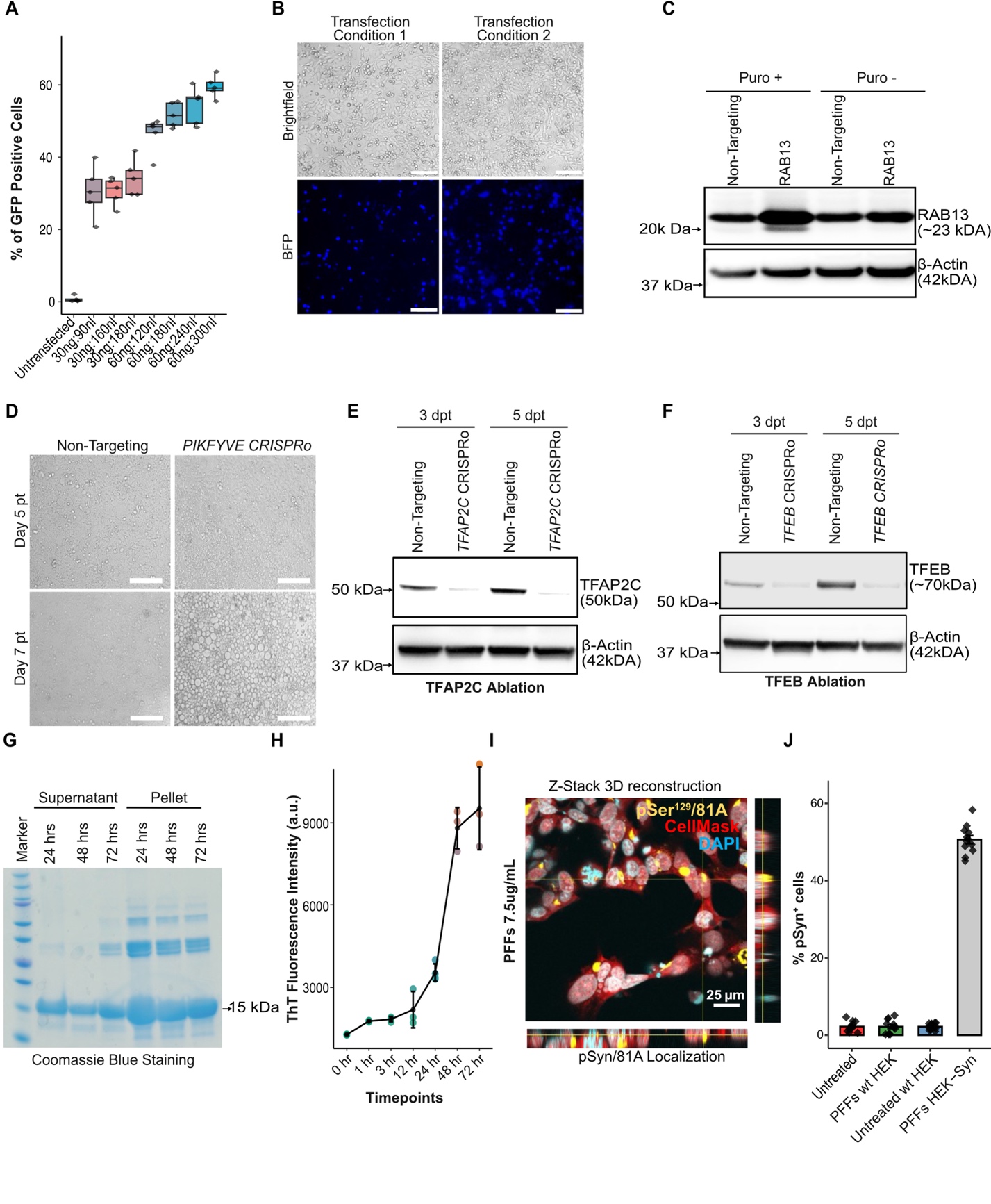

**Figure S1: High-content CRISPRa and CRISPRo α-Syn aggregation assay validation and workflow optimization.** **(A)** Transfection efficiency optimization in a 384-well plate using different ratios of plasmid DNA concentration to the transfection reagent Viafect. Box plots display the median (centre line), the 75th percentile (top edge), and the 25th percentile (bottom edge). **(B)** Representative micrographs showing brightfield (BF) and blue fluorescence (BFP) under two transfection conditions: (Left) Day 1 seeding at 5000 cells followed by transfection on Day 2; (Right) Day 1 seeding at 3000 cells followed by transfection on Day 3. Scale bar: 200 µm. **(C)** Western blot showing dCas9 activity in RAB13 CRISPRa guide-transfected cells under puromycin selection (+Puro) and without selection (-Puro). **(D)** Brightfield images for *PIKFYVE* ablation causes extensive vacuolation in CRISPRo cells. Scale bar: 200 µm. pt: post transfection. **(E, F)** Western blot analysis of TFAP2C and TFEB ablation efficiency in CRISPRo HEK293 cells at 3 and 5 days post-transfection (dpt). **(G)** Coomassie-stained gel showing progressive fibrillation of α-Syn over 72 hours. **(H)** Thioflavin T (ThT) assay to measure α-Syn fibrillation kinetics. Data are presented as mean ± SEM. **(I)** In vitro assessment of α-Syn PFF transduction using the transfection reagent Mirus in HEK293 cells, with 3D Z-stack reconstruction showing phosphorylated α-synuclein at Ser^129^ /81A localisation. CellMask (red), pSyn^129^ detected using the 81A (yellow), and DAPI-stained nuclei (blue). **(J)** Quantification of pSyn^129+^ cells under different experimental conditions: PFF-treated wild-type HEK cells (wt HEK; HEK293T cells with no or very low α-Syn expression) and PFF-treated HEK293 cells overexpressing α-Syn. Data: mean ± SEM.

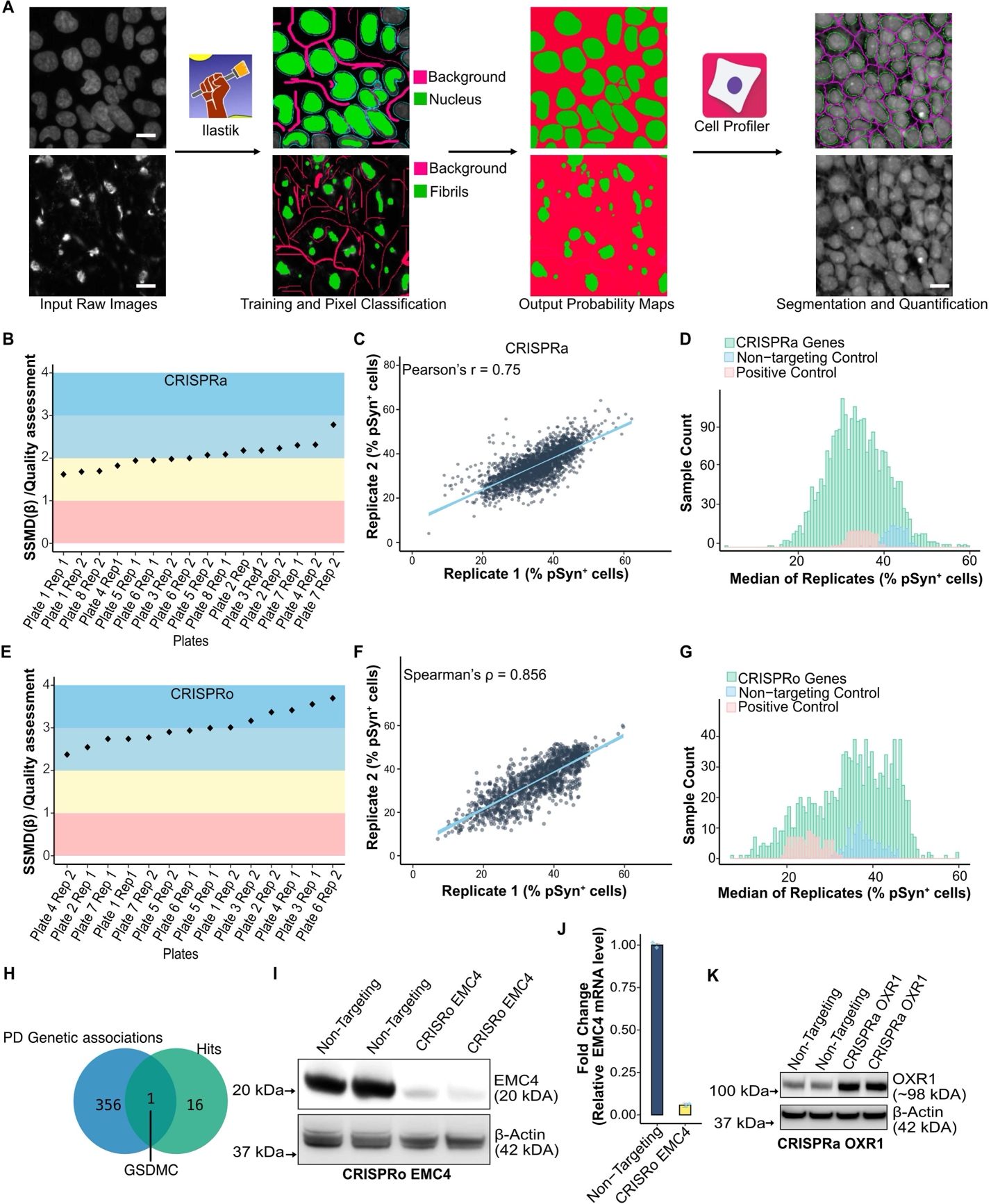

**Figure S2: Development of image analysis pipeline, data quality assessment, and formal validation of key hits.** **(A)** Workflow for segmentation and quantification using ilastik for pixel classification (nuclei: pink; fibrils: green; background: red) and CellProfiler for segmentation and quantification. Scale bar: 10 µm. (**B, E)** Strictly standardised mean difference (SSMD) scores assessing the quality of individual CRISPRa (**B)** and CRISPRo (**E)** plates based on non-targeting and positive controls. **(C), (F)** Scatter plots showing a correlation between duplicates of the primary CRISPRa (Pearson’s correlation) **(C)** and CRISPRo **(F)** (Spearman’s correlation) screens. (**D, G)** Histograms showing the frequency distribution of individual CRISPRa **(D)** and CRISPRo **(G)** screens, including non-targeting and positive controls. **(H)** Venn diagram showing the overlap of validated CRISPRa and CRISPRo hits with Parkinson's disease genetic associations from the Open Targets platform. **(I)** Western blot analysis of EMC4 protein levels in non-targeting and CRISPRo *EMC4* ablation conditions. (**J)** RT-qPCR showing the relative mRNA levels of *EMC4* in CRISPRo *EMC4* ablation cells. **(K)** Western blot analysis of OXR1 protein levels in CRISPRa *OXR1*-activated cells.

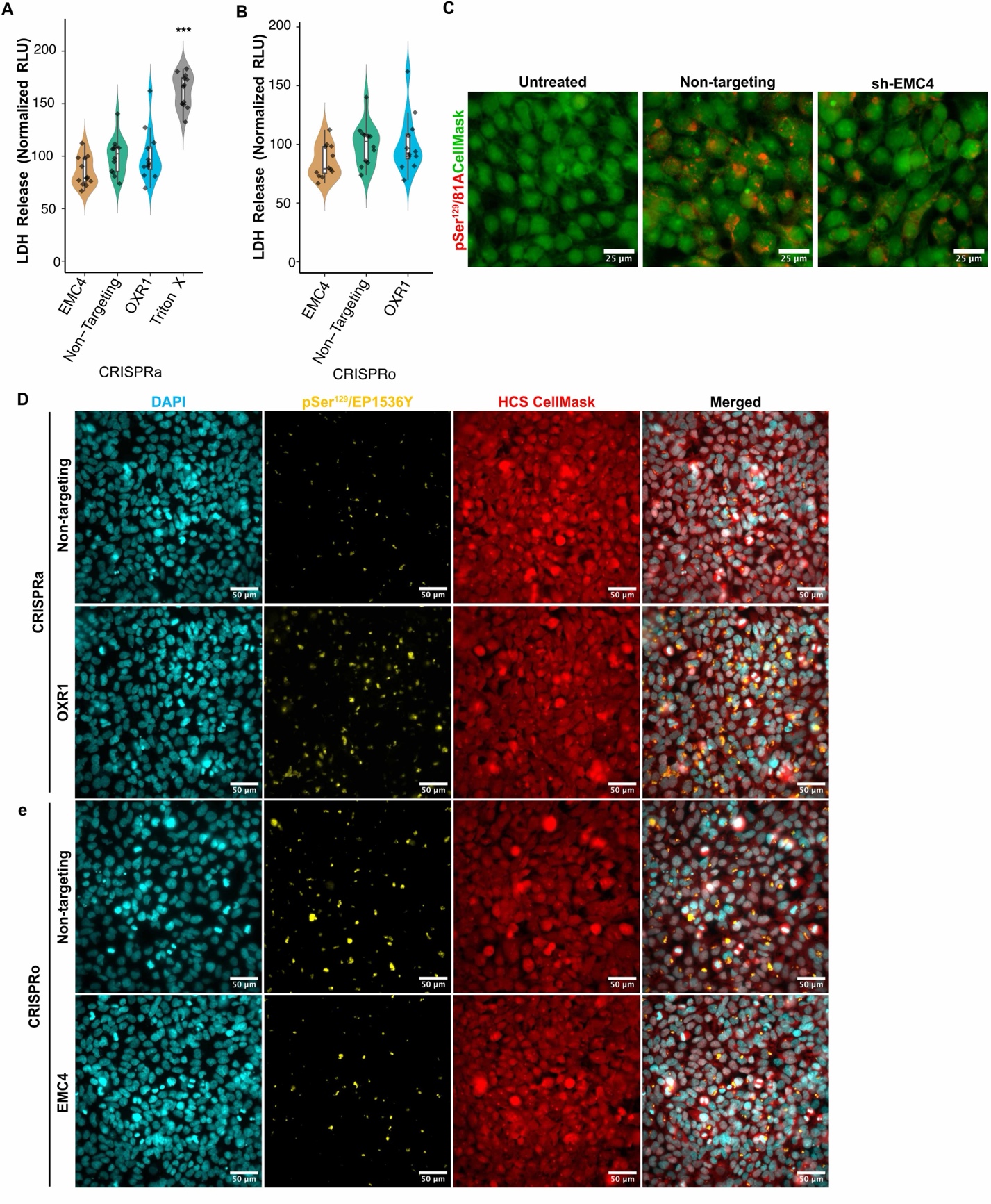

**Figure S3:** **Toxicity assessment and representative immunofluorescence images of HEK^Syn^ cells showing phosphorylated α-synuclein at Ser129 (pSyn^129^) aggregates.** **(A,B)** Measurement of lactate dehydrogenase (LDH) release as a marker of cytotoxicity, expressed as relative luminescence units (RLU) and normalised to the non-targeting control. Cells treated with Triton® X-100 served as a positive control. **(C)** Representative micrographs showing p-Ser^129^ α-Syn aggregates detected using the 81A antibody (red), and whole-cell staining with HCS CellMask (green). **(D-E)** DAPI-stained nuclei (cyan), pSyn^129^ aggregates detected with the EP1536Y antibody (yellow), and whole-cell staining with HCS CellMask (red). Rows represent different experimental conditions. CRISPR activation **(D)** and CRISPR ablation **(E)**. Violin plots represent data distribution. Inner box plots display the median (centre line), the 75th percentile (top edge), and the 25th percentile (bottom edge). Statistical comparisons were performed using one-way ANOVA followed by Dunnett's post hoc test, with significance levels denoted as follows: *P < 0.05, **P < 0.01, ***P < 0.001.

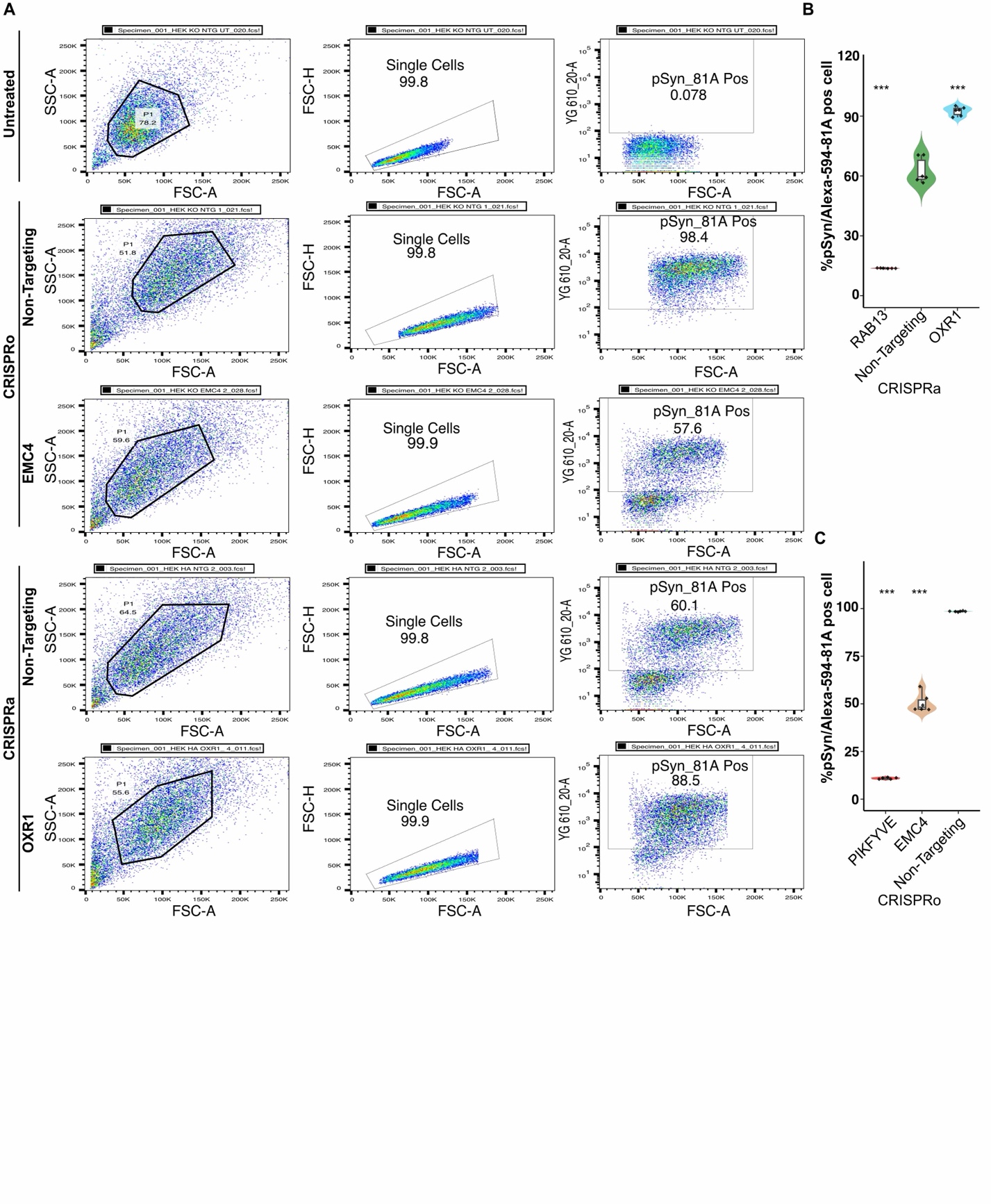

**Figure S4:** **Gating strategy for flow cytometry and measurement of phosphorylated α-synuclein at Ser^129^ (pSyn^129^) positive cells.** **(A)** Gating strategy for CRISPRo hits (upper panel) and for CRISPRa (lower panel). Representation of flow cytometry plots displaying single cell population and the pSyn^129^ (Alexa Fluor 594-labelled 81A antibody) positive cells. **(B)** Percentage of p-Ser^129^-positive cells across CRISPRa hits, based on flow cytometry analysis using the Alexa Fluor 594 81A antibody. **(C)** Percentage of p-Ser^129^-positive cells across CRISPRo hits, based on flow cytometry analysis using the Alexa Fluor 594 81A antibody. Violin plot quantifying the percentage of pSyn^129^ positive cells across different experimental conditions. Violin plots represent data distribution. Inner box plots display the median (centre line), the 75th percentile (top edge), and the 25th percentile (bottom edge). Statistical comparisons were performed using one-way ANOVA followed by Dunnett's post hoc test, with significance levels denoted as follows: *P < 0.05, **P < 0.01, ***P < 0.001.

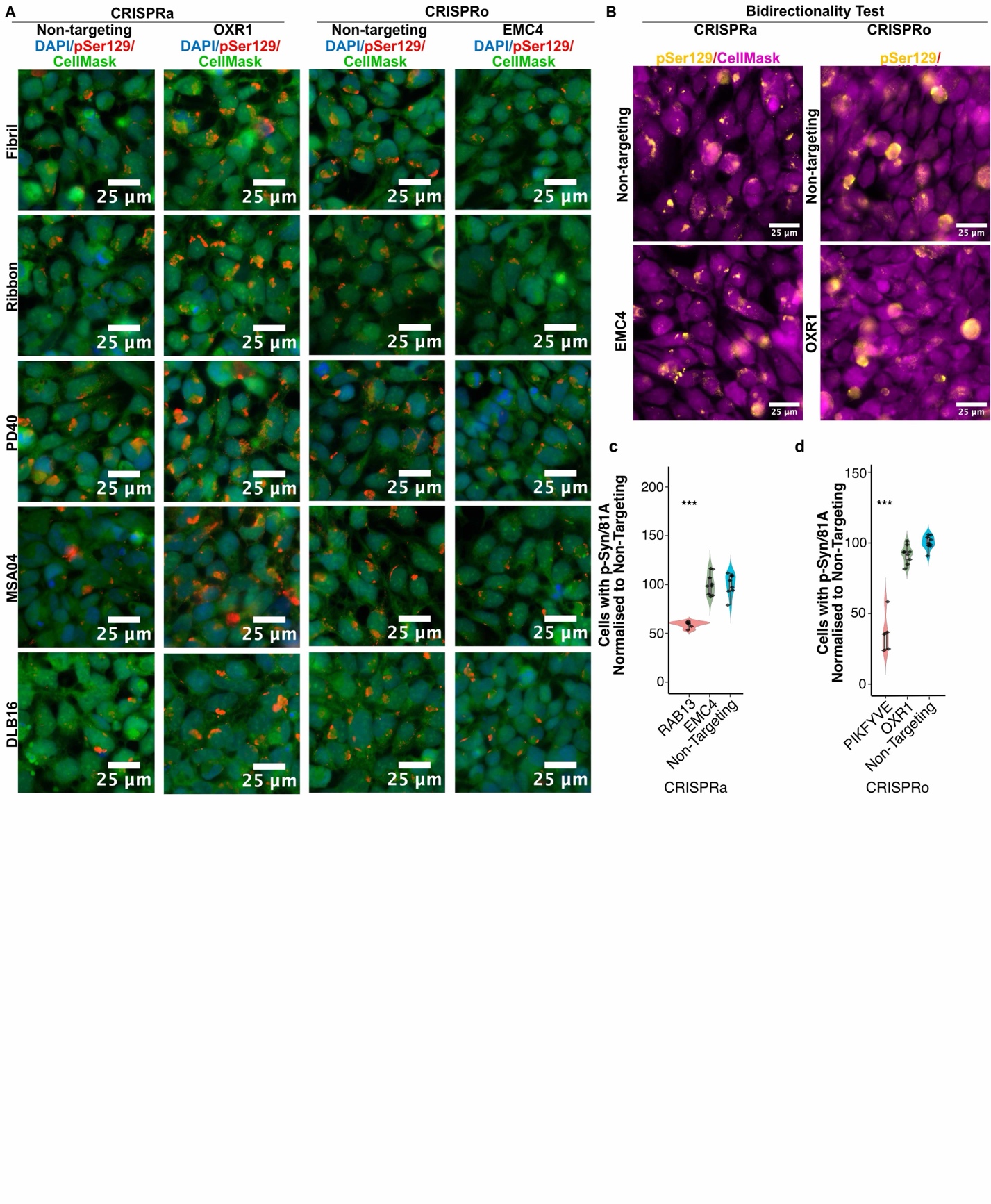

**Figure S5: Effects of CRISPR activation (CRISPRa) /CRISPR ablation (CRISPRo) perturbations on phosphorylated α-synuclein at Ser^129^ (pSyn^129^) levels in response to α-synuclein polymorphs.** **(A)** Representative immunofluorescence images of HEK293 cells showing phosphorylated α-synuclein at Ser^129^ (pSyn^129^) levels in response to human PD, DLB, and MSA patient-derived fibrils, as well as two distinct recombinant polymorphs: fibrils and ribbons strains. (Cyan: DAPI; Green: HCS CellMask; Red: pSyn^129^/81A). **(B)** Representative images illustrating the effect of hit gene perturbations (bi-directional effects) on pSyn^129^ levels using 81A antibody. (Magenta: HCS CellMask; Yellow: pSyn^129^/81A). Left column: CRISPRa; Right column: CRISPRo. **(C)** Effect activation of *EMC4* on pSyn^129^ levels. **(D)** Effect on pSyn^129^ levels following CRISPRo-mediated ablation of *OXR1*.

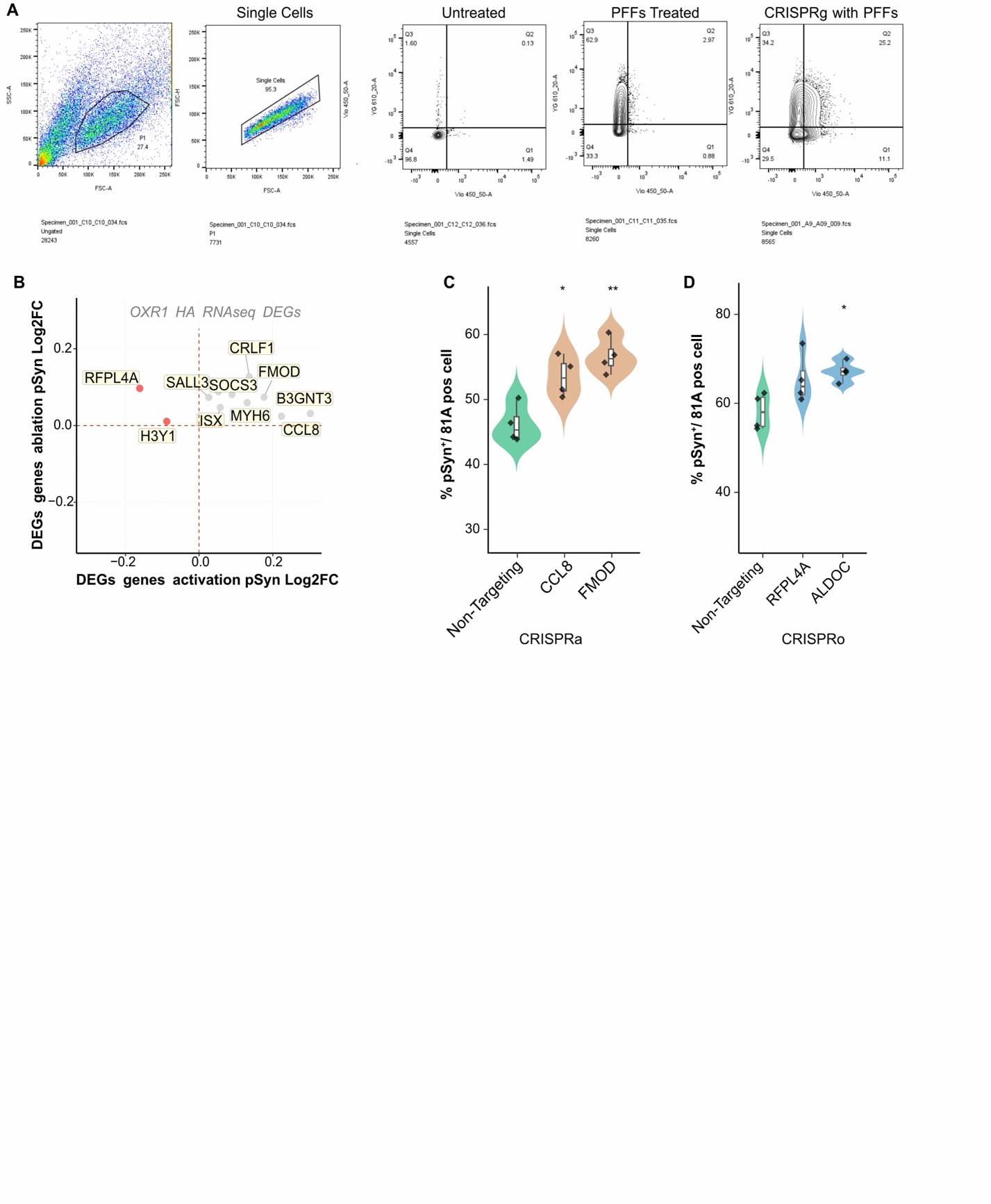

**Figure S6:** **Flow cytometry gating strategy and intersection analysis of differentially expressed genes (DEGs) modulating phosphorylated α-synuclein at Ser129 (pSyn^129^).** **(A)** Flow cytometry gating strategy for the mini-screen of DEGs, including OXR1 activation and EMC4 ablation conditions. **(B)** Intersection of log2 fold change in pSyn^129^ levels for individual perturbed DEGs upon OXR1 activation. **(C-D)** Effect of OXR1 activation DEGs on pSyn^129^ assed with immunofluorescence imaging. Violin plots represent data distribution. Inner box plots display the median (centre line), the 75th percentile (top edge), and the 25th percentile (bottom edge). One-way ANOVA followed by Dunnett's post hoc test; *P < 0.05, **P < 0.01, ***P < 0.001.

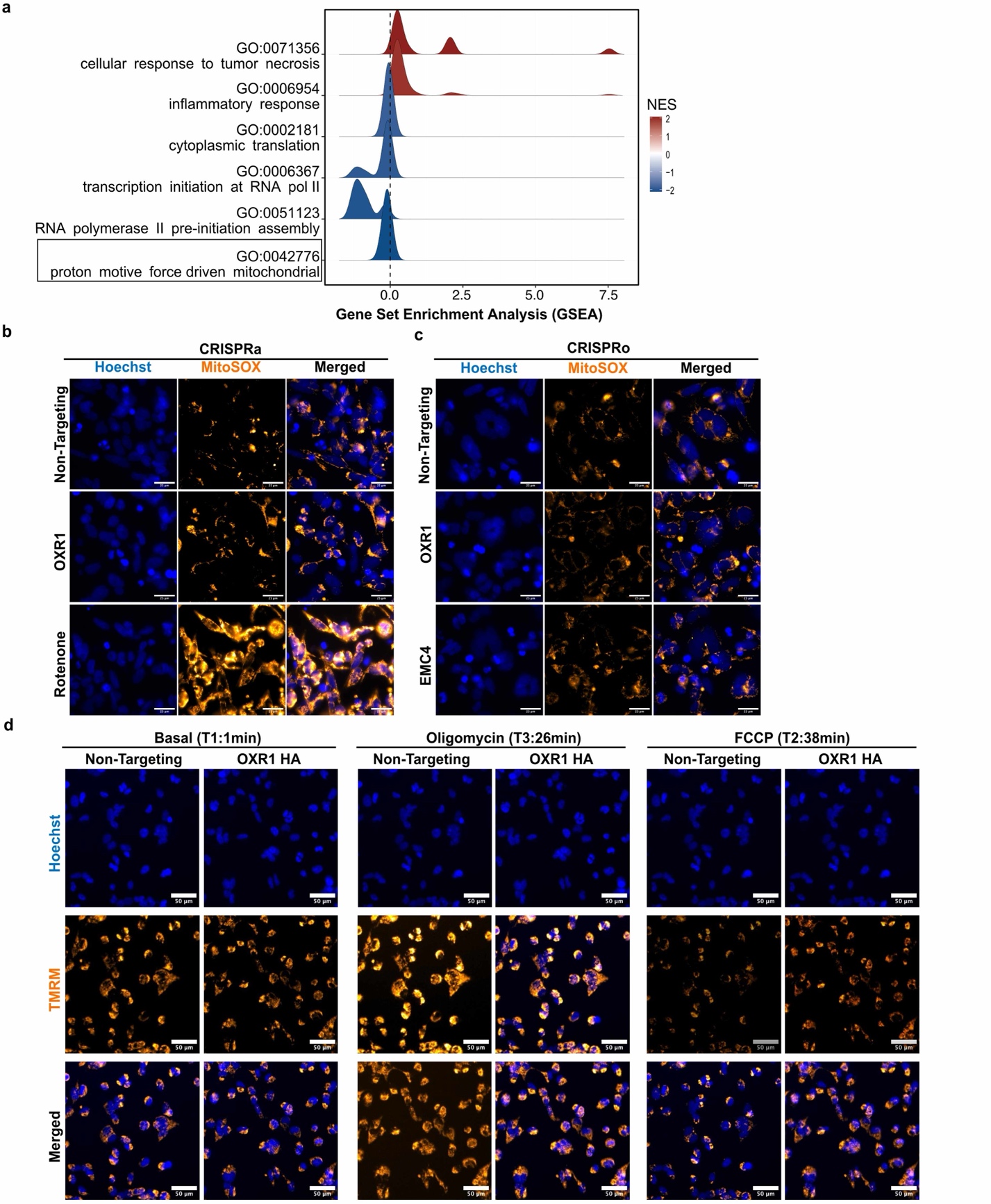

**Figure S7: Pathway enrichment analysis, measurement of MitoSOX-based superoxide levels and TMRM-based mitochondrial membrane potential using live-cell imaging. (A)** Gene set enrichment analysis (GSEA) shows pathways ranked by NES and highlights mitochondria-related pathways. Candidate terms were filtered based on False discovery rate (FDR) ≤ 0.05, and genes were ranked by log2 fold change. **(B)** Representative immunofluorescence images showing mitochondrial superoxide levels stained with MitoSOX™ Red dye upon activation of the hits. (Hoechst-stained nuclei: Blue; Orange-hot: MitoSOX™ Red). Rotenone-treated conditions served as the positive control. Scale bar: 25 μm. **(C)** Representative fluorescence images showing mitochondrial superoxide levels stained with MitoSOX™ Red dye upon ablation of the hits. (Hoechst-stained nuclei: Blue; Orange-hot: MitoSOX™ Red). Scale bar: 25 μm. **(D)** Representative immunofluorescence microscopy images showing mitochondrial membrane potential. (Hoechst-stained nuclei: Blue; Orange-hot: TMRM). Images shown were taken at basal conditions (1 min) and following treatment with oligomycin (26 min) and FCCP (38 min).

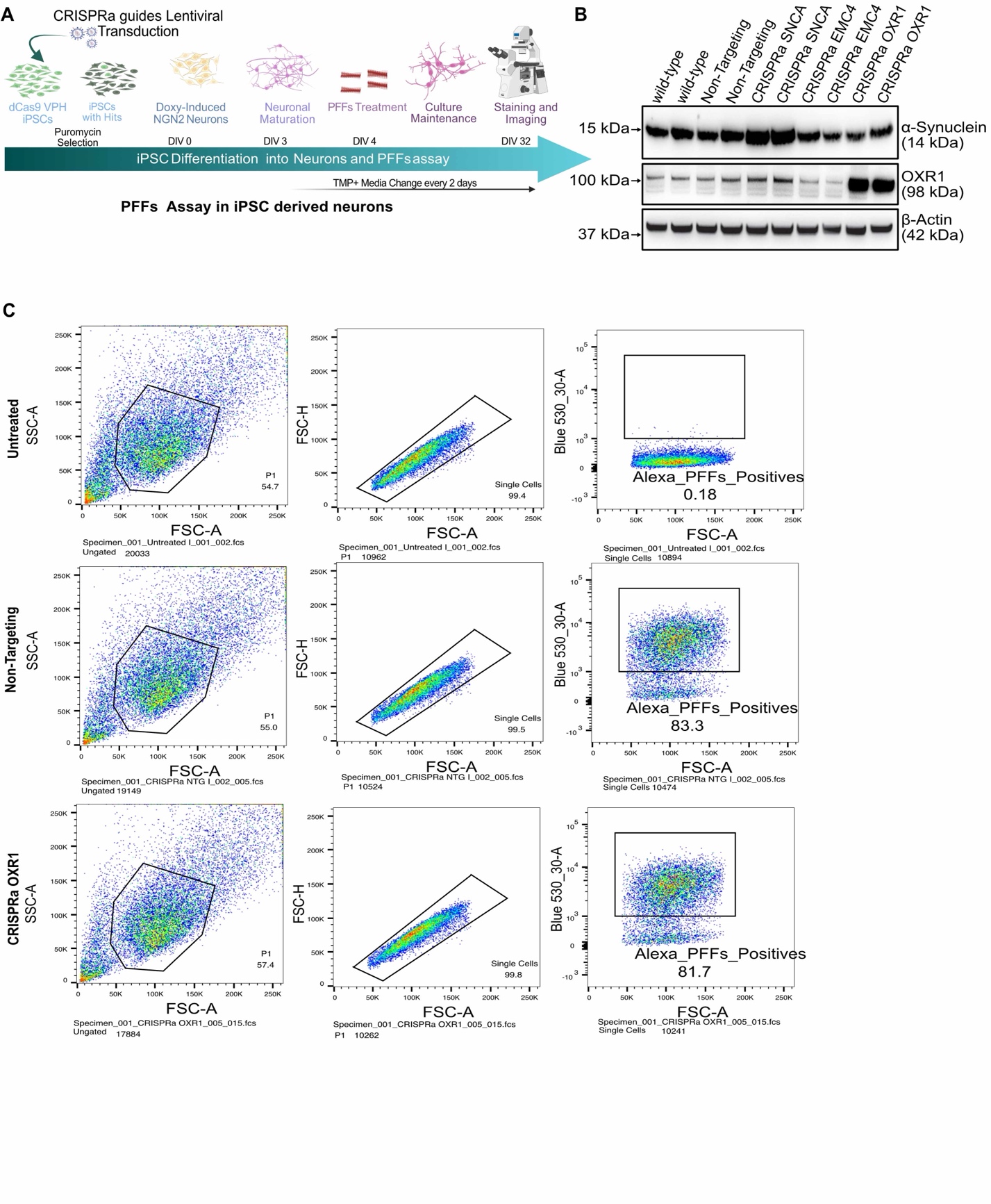

**Figure S8: Generation and analysis of human iPSC-derived cortical neurons.** **(A)** Schematic of iPSC line generation, differentiation, maturation, and PFF treatment in iPSC-derived cortical neurons. **(B)** Immunoblot analysis of α-synuclein levels in iPSC-derived cortical neurons. **(C)** Gating strategy for flow cytometry plots displaying single-cell populations and Alexa Fluor 488-positive cells.

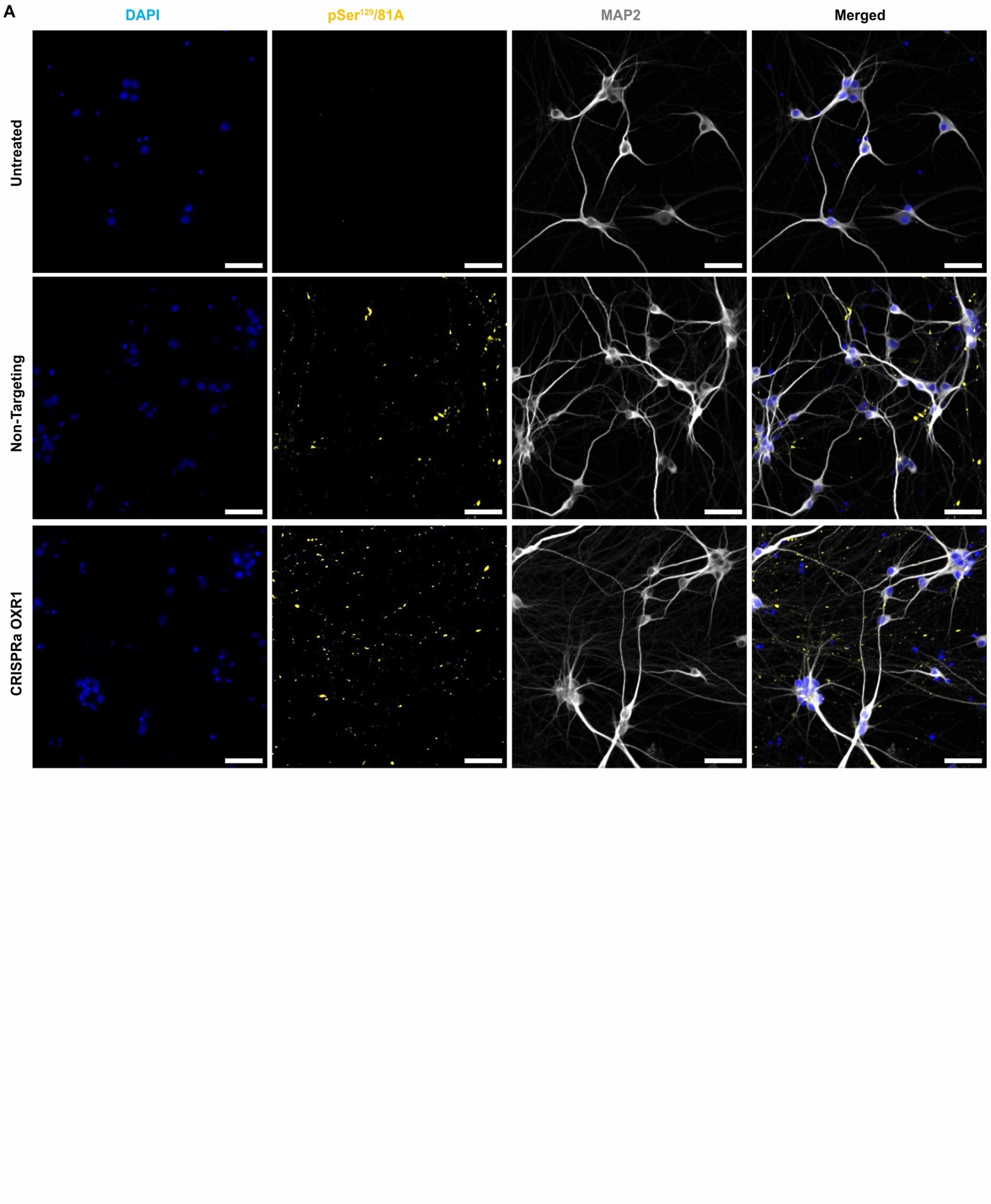

**Figure S9. Assessment of α-synuclein aggregates level in iPSC-derived dopaminergic neurons.** **(A)** Representative immunofluorescence images of human iPSC-derived dopaminergic neurons treated with CRISPRa targeting OXR1 or non-targeting (NTG) controls. iPSC-derived dopaminergic neurons were stained for nuclei (DAPI, blue), neuronal marker MAP2 (grey), and phosphorylated α-synuclein at Ser^129^ (pSyn^129^, detected via 81A antibody, yellow)**.** Scale bar: 25 µm.

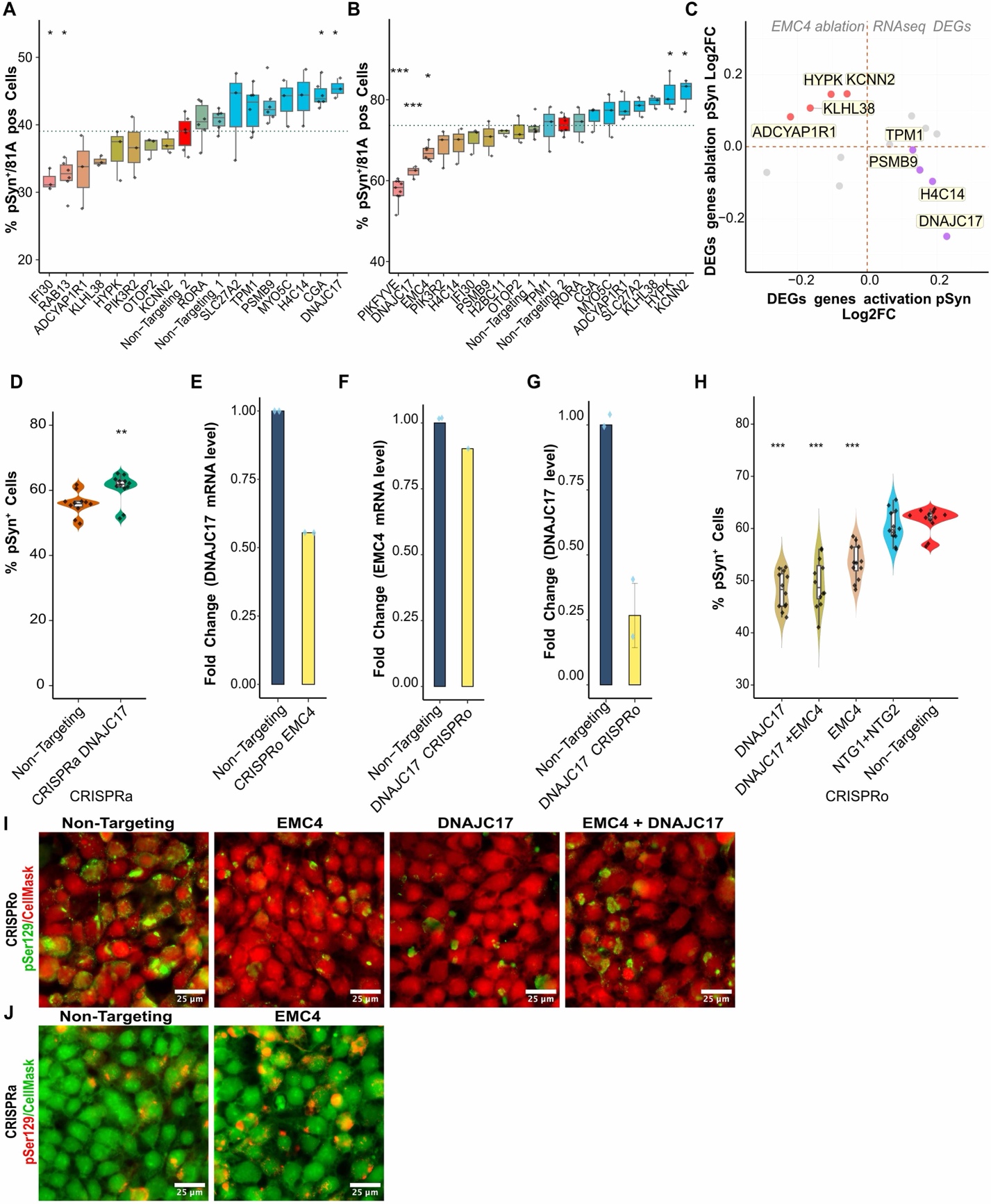

**Figure S10:** **Transcriptomic changes and functional effects of downstream genes on phosphorylated α-synuclein at Ser^129^ (pSyn^129^). (A-B)** Individual perturbation of DEGs upon EMC4 ablation. Flow cytometry-based percentage of pSyn^129+^cells following CRISPR activation (**A**) and CRISPR ablation (**B**). **(C)** Intersection of log2 fold changes in pSyn^129^ levels for individual perturbed DEGs upon EMC4 ablation. **(D)** Effect of *DNAJC17*activation on pSyn^129^ levels. Statistical comparisons were performed using Welch’s t-test (unequal variance t-test). **(E)** RT-qPCR analysis of *DNAJC17*mRNA levels upon EMC4 ablation. **(F)** RT-qPCR analysis of EMC4 mRNA levels upon *DNAJC17*ablation. **(G)** RT-qPCR analysis of *DNAJC17*mRNA levels upon *DNAJC17*ablation. **(H)** Effect of the combined ablation of EMC4 and *DNAJC17*on pSyn^129^ levels. **(I)** Representative immunofluorescence images showing pSyn^129^ levels in cells with combined ablations of *EMC4* and *DNAJC17* (Green: HCS CellMask; Red: pSyn^129^/81A). **(J)** Representative immunofluorescence images showing pSyn^129^ levels in DNAJC17-activated cells (Green: HCS CellMask; Red: pSyn^129^/81A). Bar plots data are presented as mean ± SEM. Violin plots represent data distribution. Box plots display the median (centre line), the 75th percentile (top edge), and the 25th percentile (bottom edge). Statistical comparisons were performed using one-way ANOVA followed by Dunnett's post hoc test. *P < 0.05, **P < 0.01, ***P < 0.001.

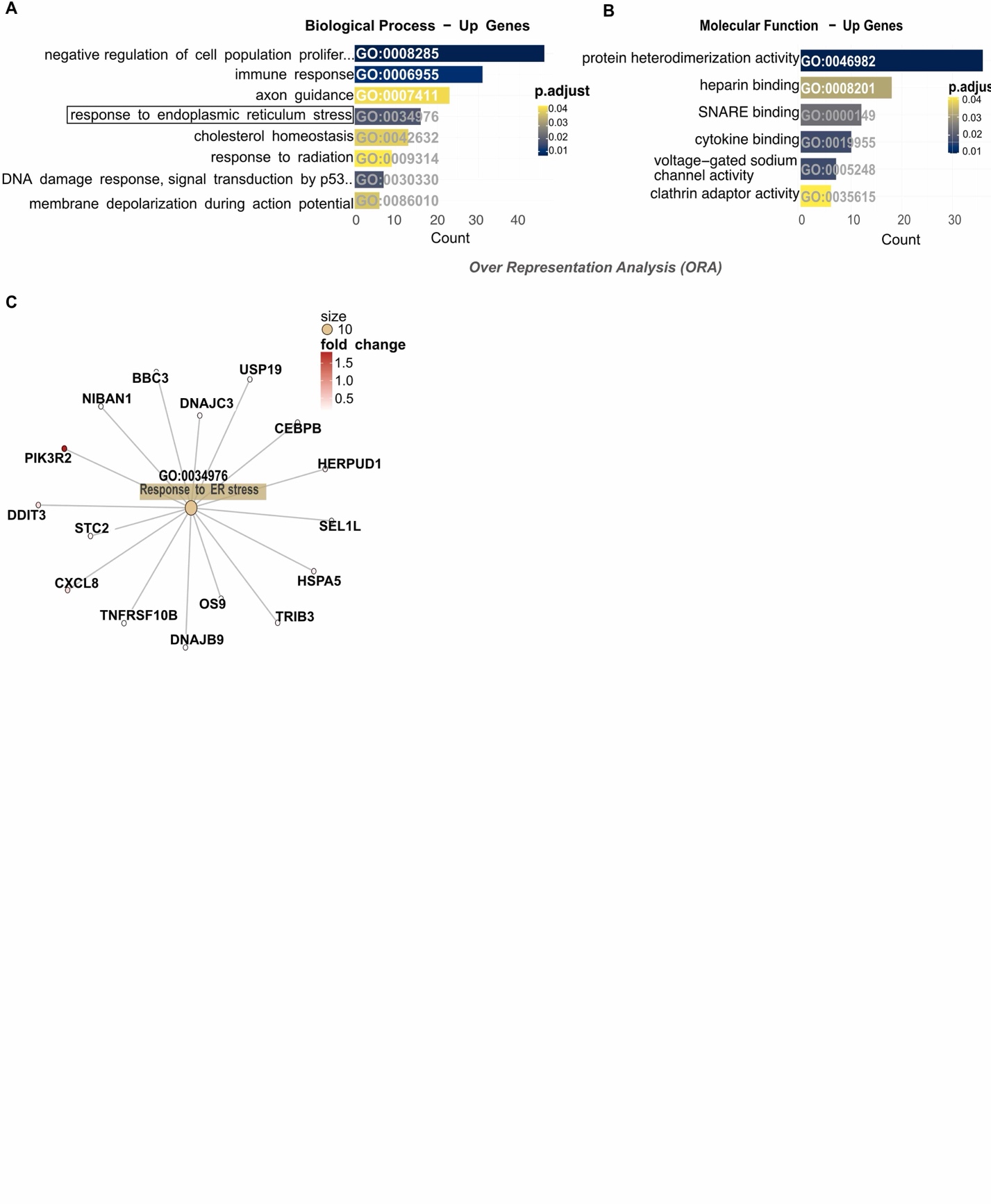
**Figure S11: Pathway enrichment analysis in *EMC4* ablation cells.** **(A)** Over-representation analysis (ORA) of biological process for upregulated genes. Bar plot summarises enriched GO terms, highlighting ER stress-related pathways. Cut-off: Candidate terms FDR ≤ 0.05. **(B)** ORA of molecular functions for upregulated genes. Cut-off: Candidate terms FDR ≤ 0.05. **(C)** Network analysis of the response to endoplasmic reticulum stress (GO:0034976). Nodes represent genes, edges denote gene-term associations, and node colour intensity reflects fold change. Data are derived from RNA sequencing of *EMC4* ablation vs. Non-targeting control cells.

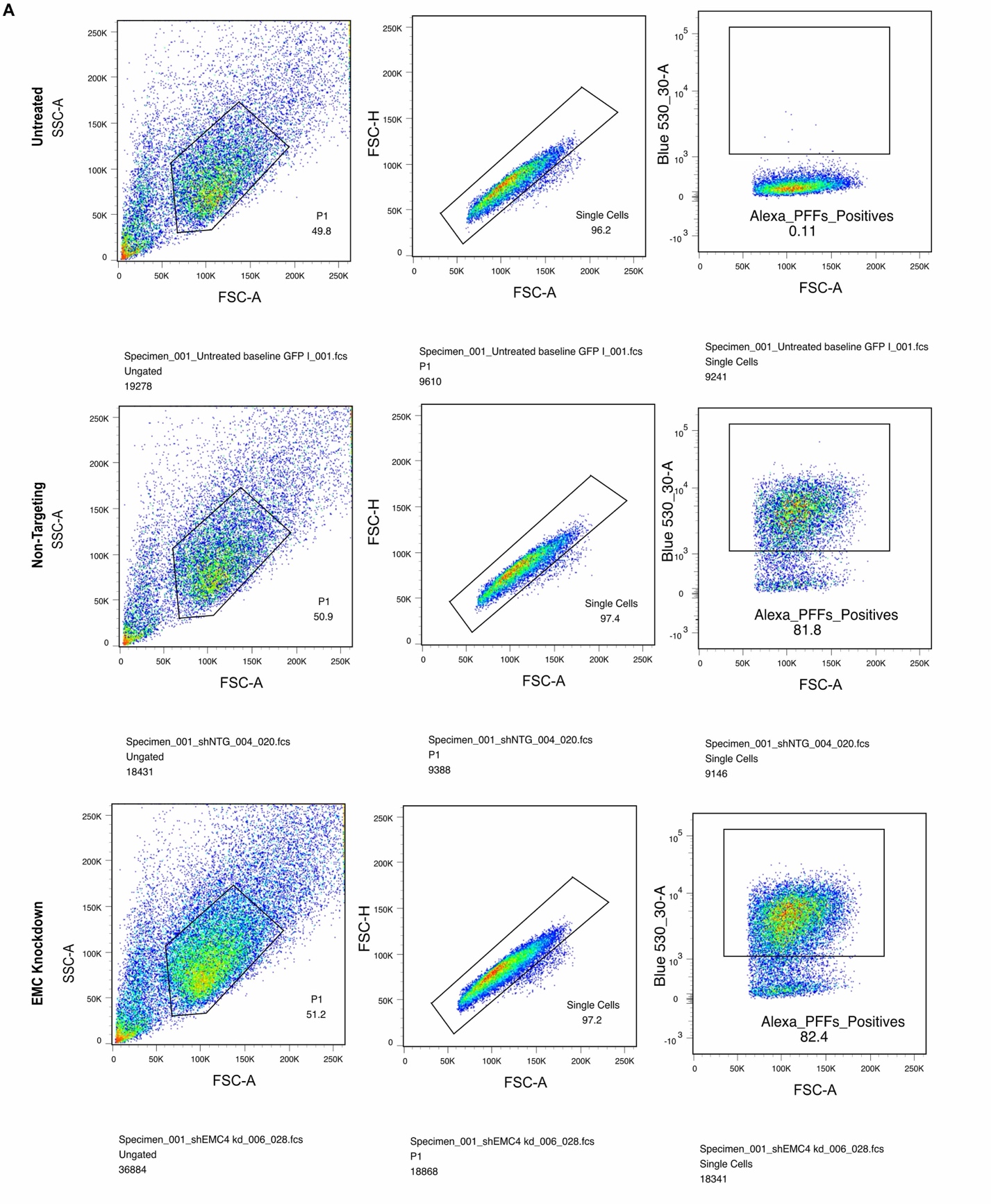

**Figure S12. Gating strategy for quantifying α-Syn PFF uptake by flow cytometry.**

**(A)** Representative flow cytometry plots illustrating the sequential gating strategy used to quantify Alexa Fluor 488-labelled α-Syn PFF-positive cells across experimental conditions.
