## Supplementary material for "Large-scale bidirectional arrayed genetic screens identify *OXR1* and *EMC4* as modifiers of α-synuclein aggregation": Document S1.docx

**Western Blot Uncropped Data**

**Figure 1**

**Figure 1a**

i dCas9 and Cas9

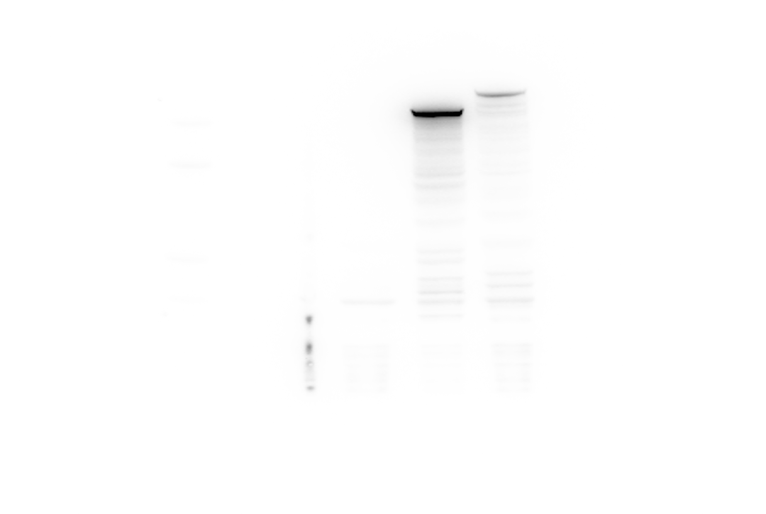

ii B-actin

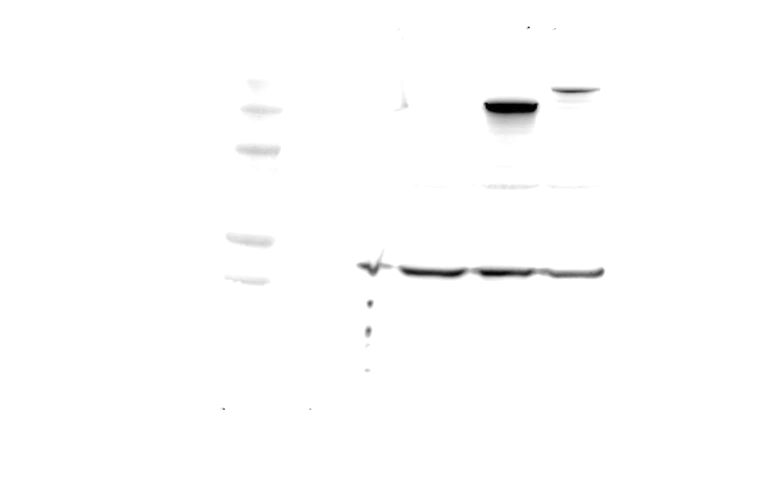

iii α-Synuclein

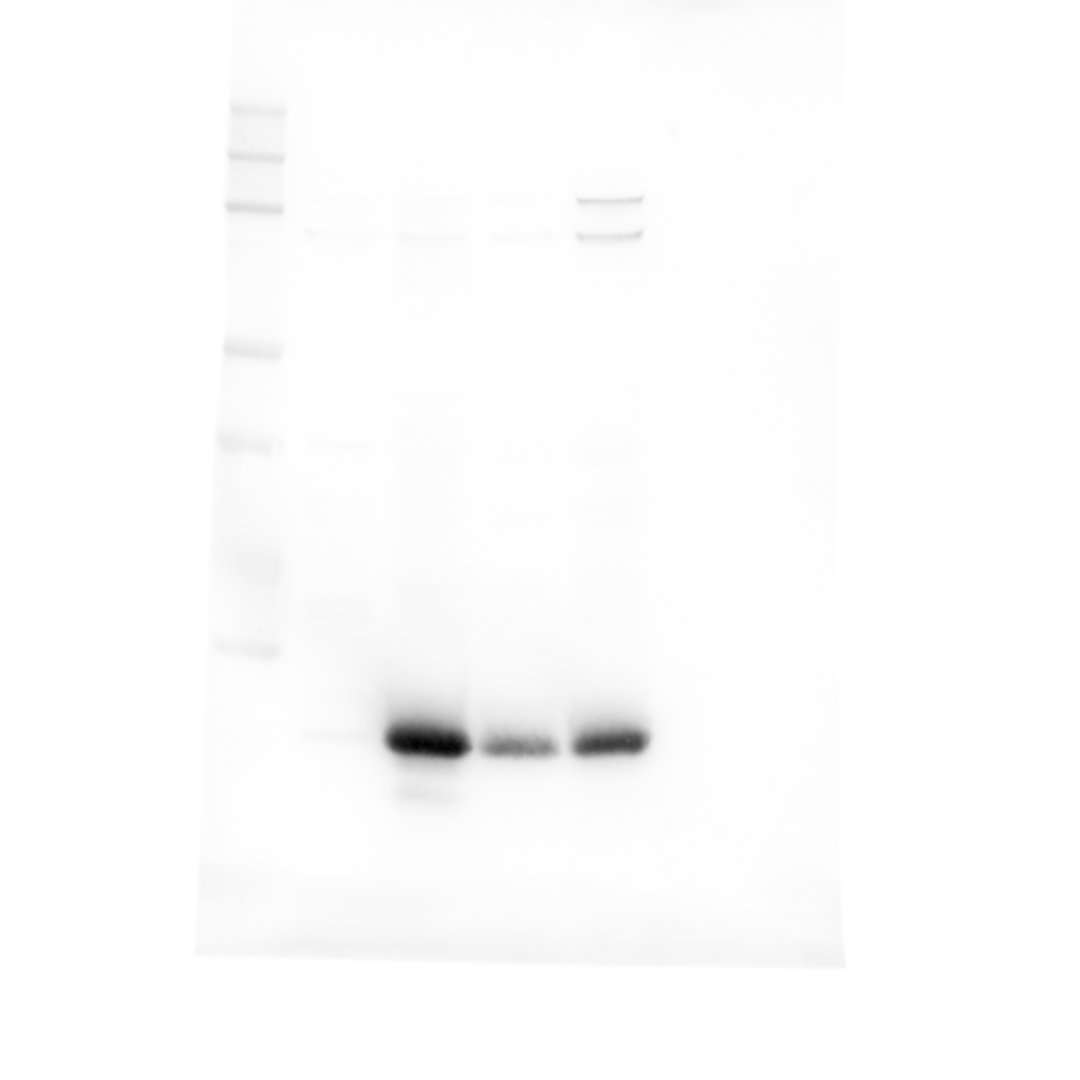

iv B-Actin

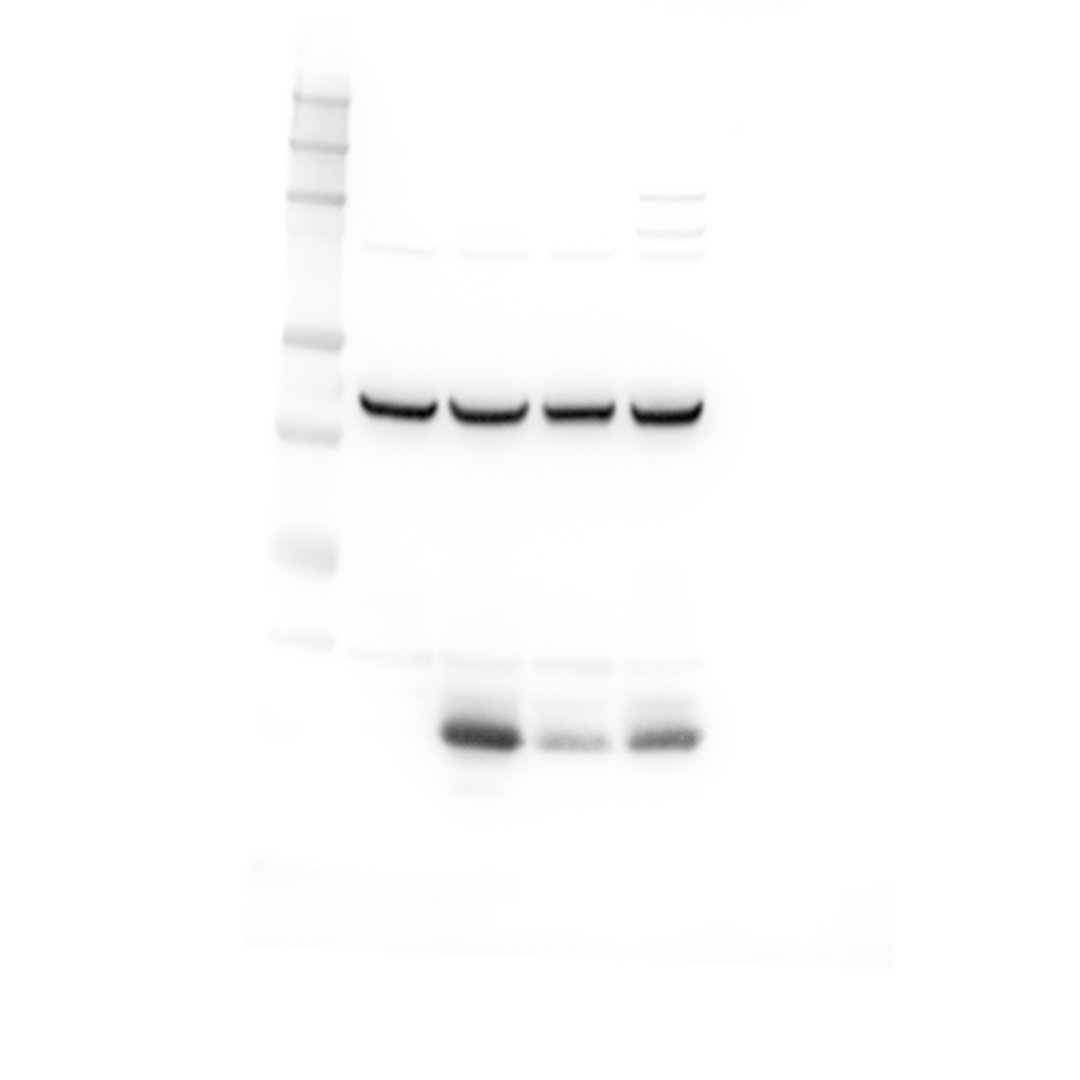

**Figure 1c**

i KDM4A

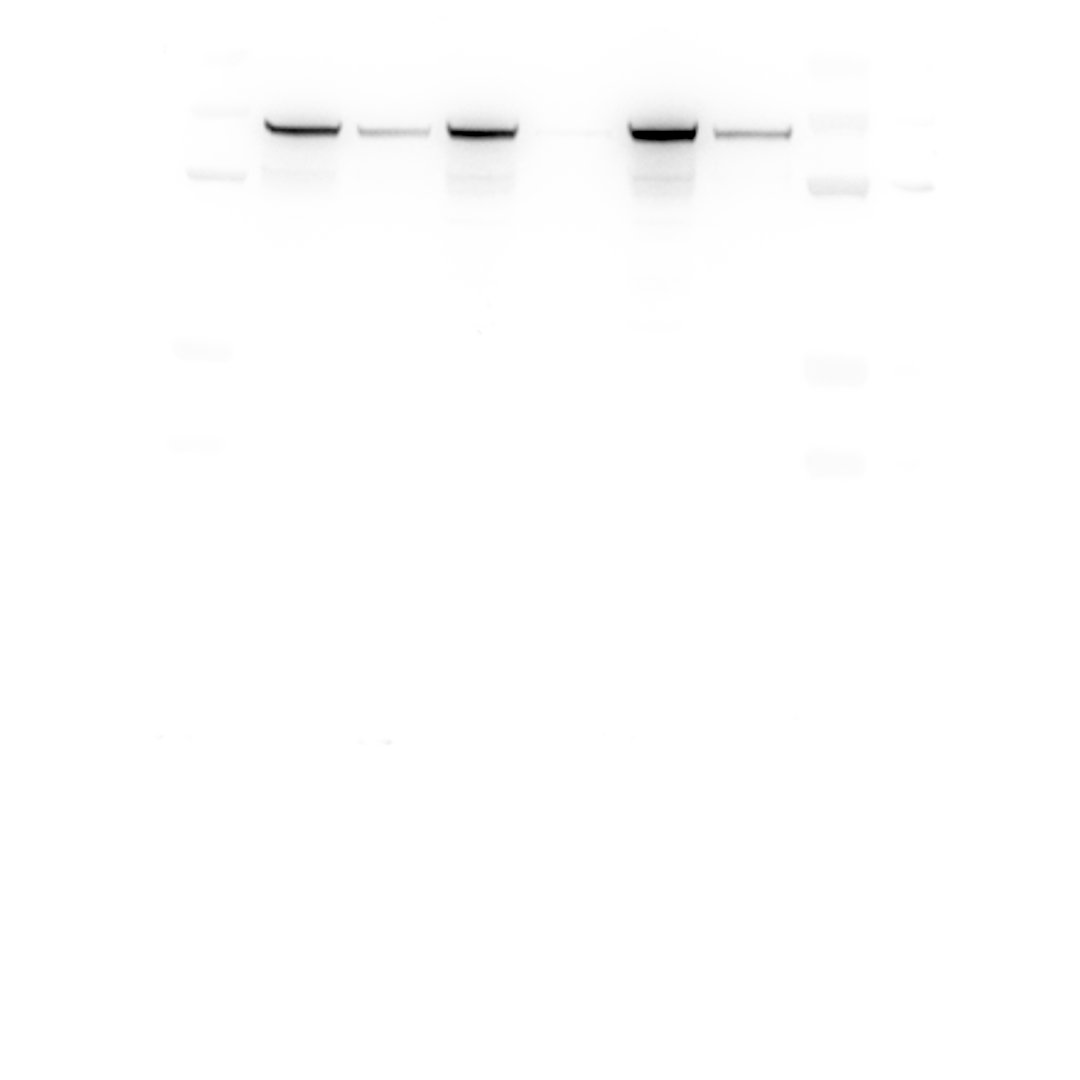

ii B-Actin

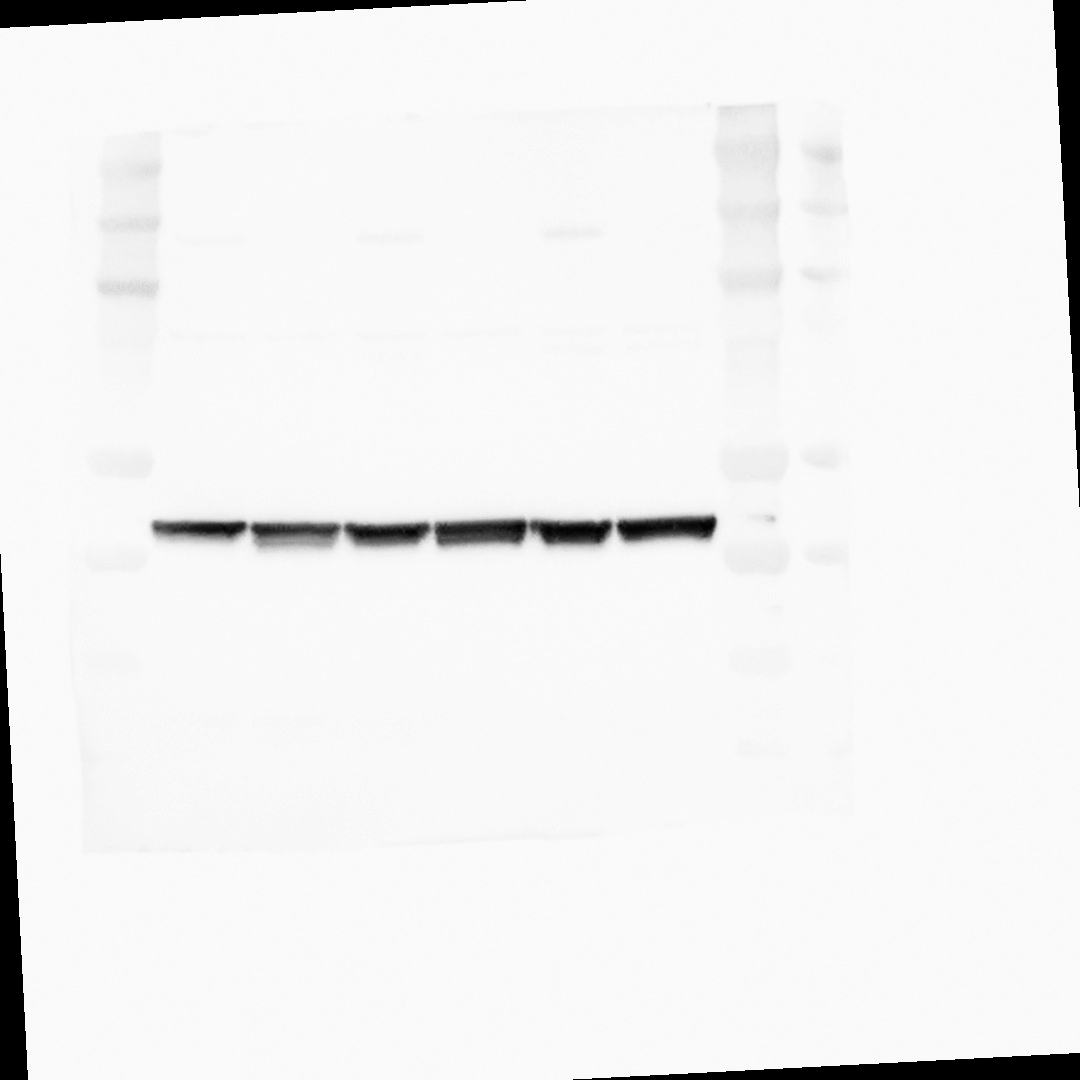

**Figure 3**

**Figure 3c**

i EMC4

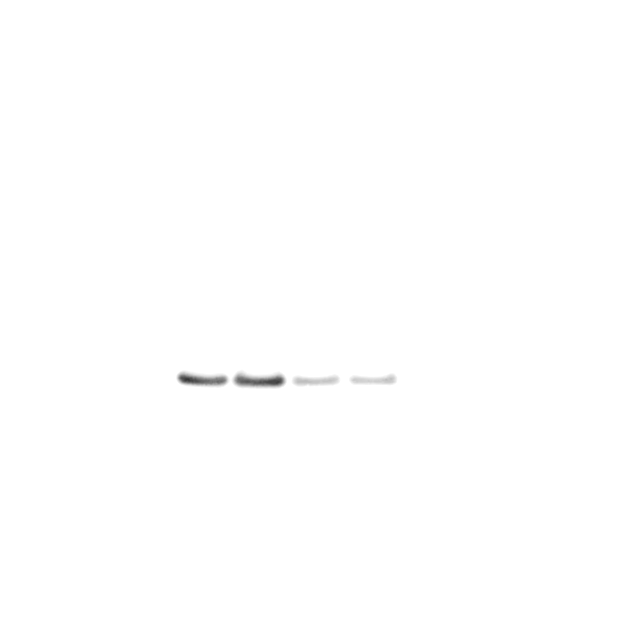

ii B-Actin

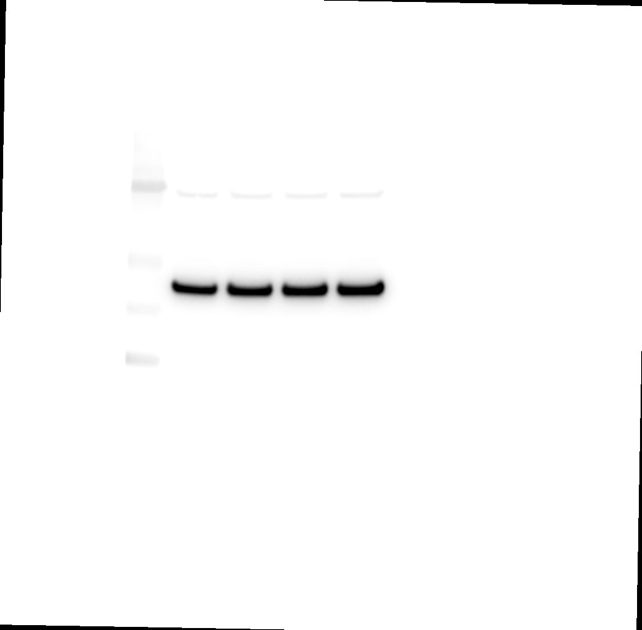

iii α-Synuclein

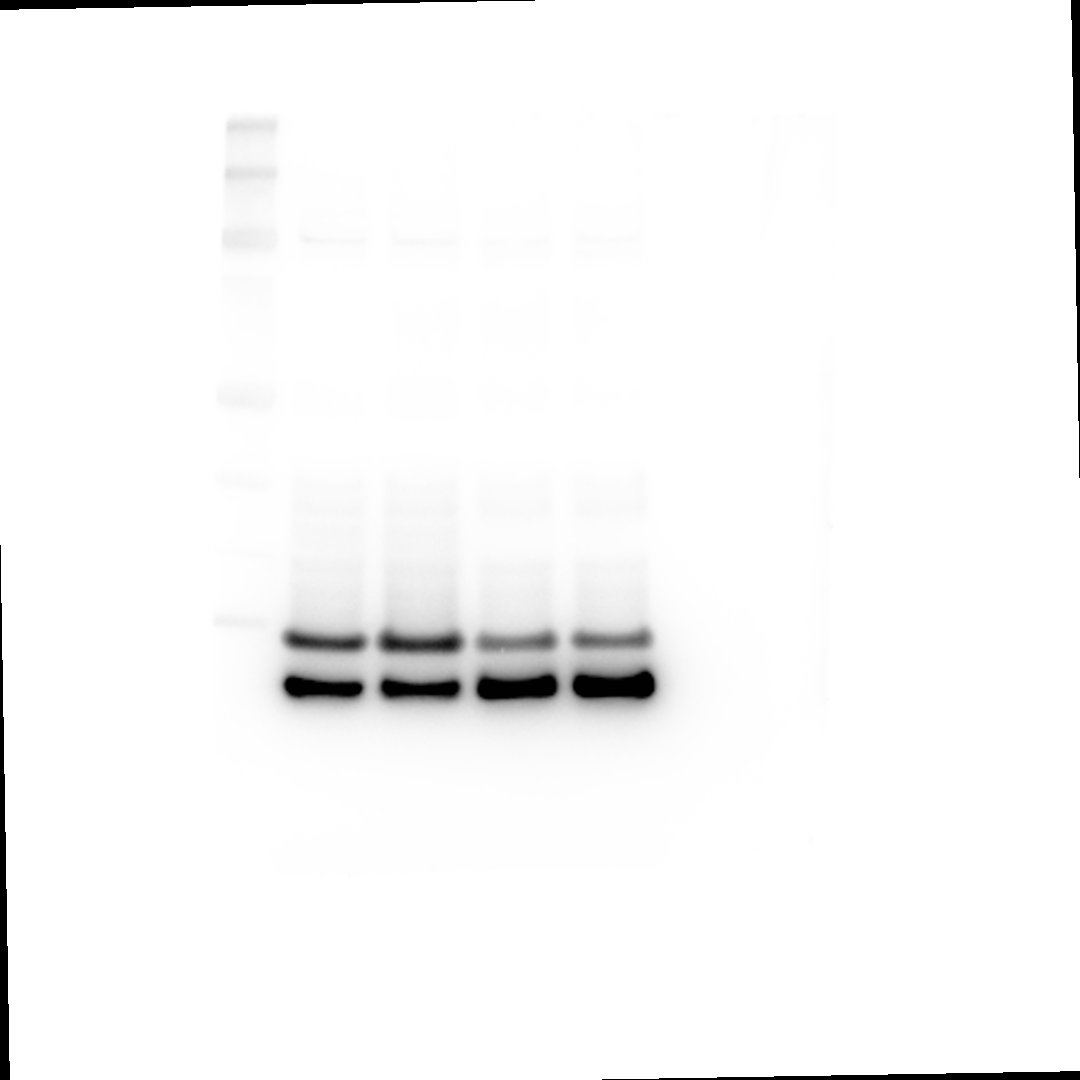

**Figure 5**

**Figure 5c**

i OXR1

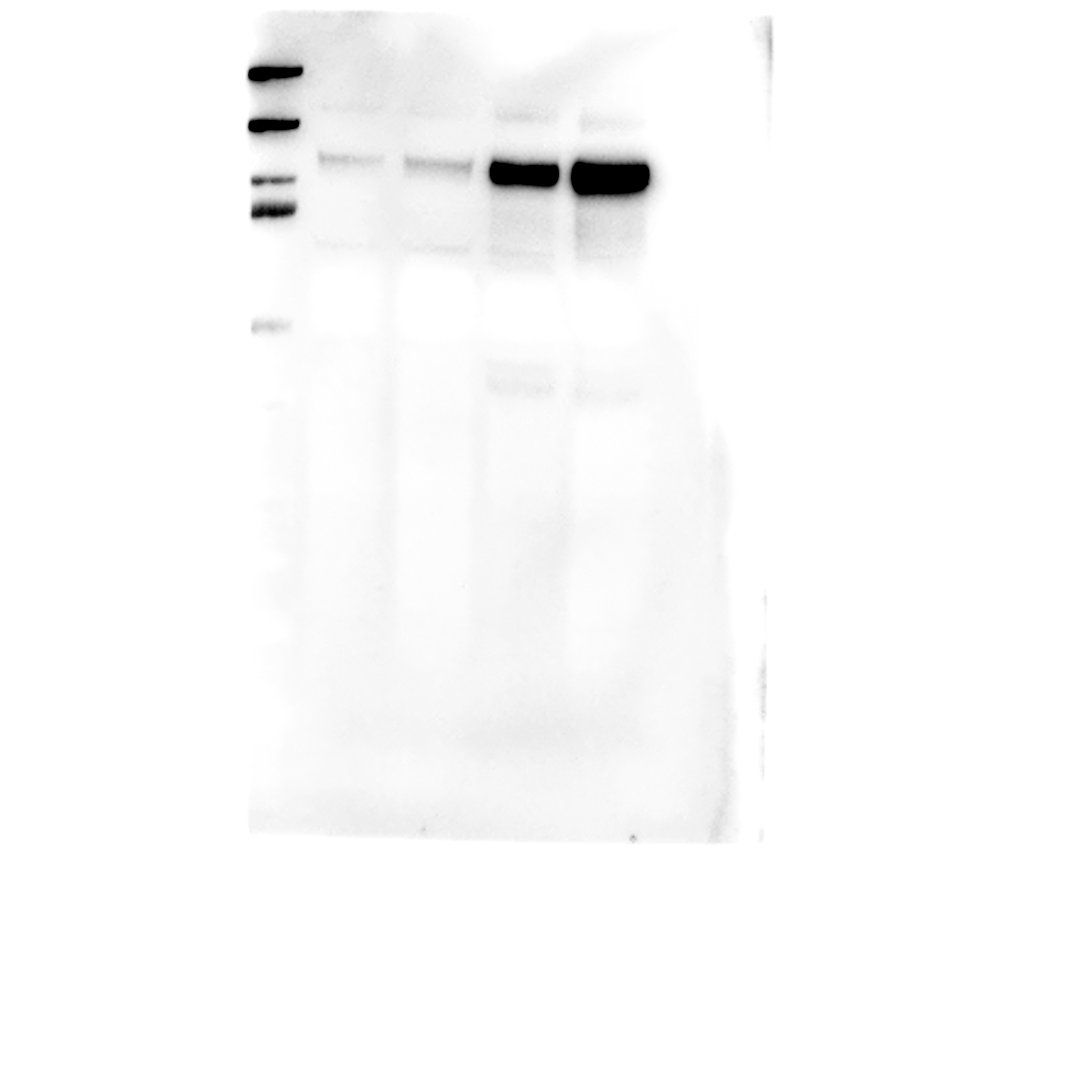

ii B-Actin

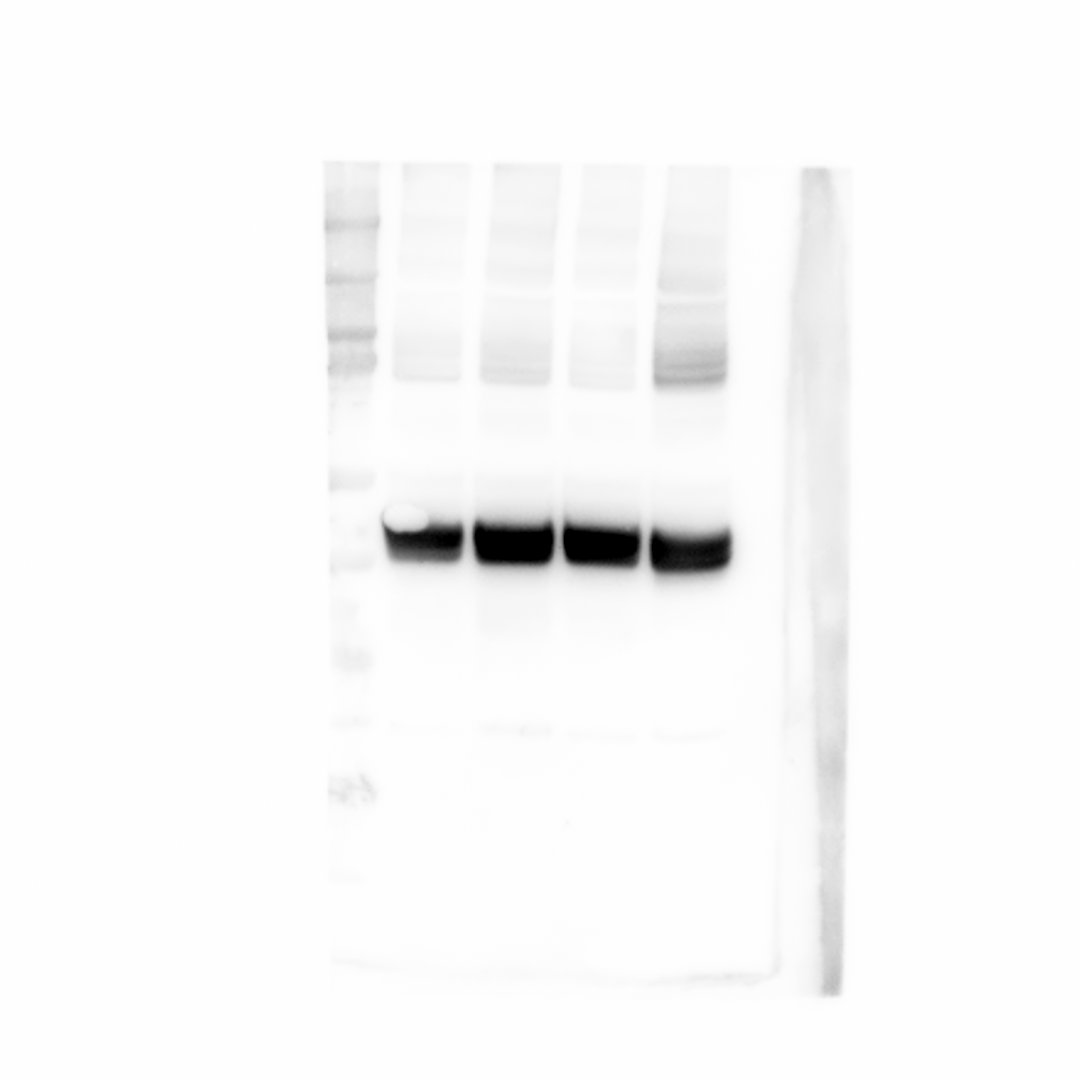

**Figure 6**

**Figure 6c**

i α-Synuclein

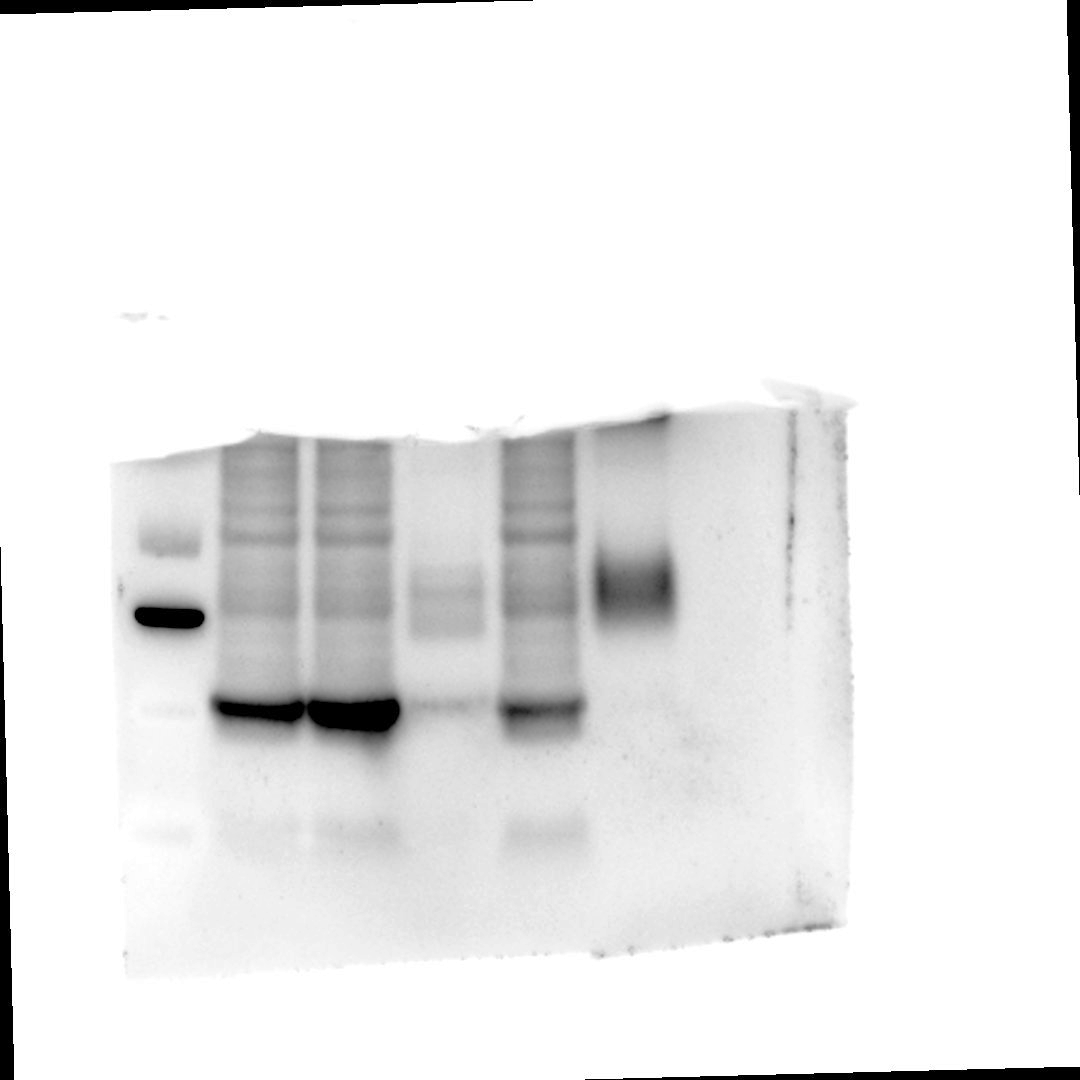

ii EMC4

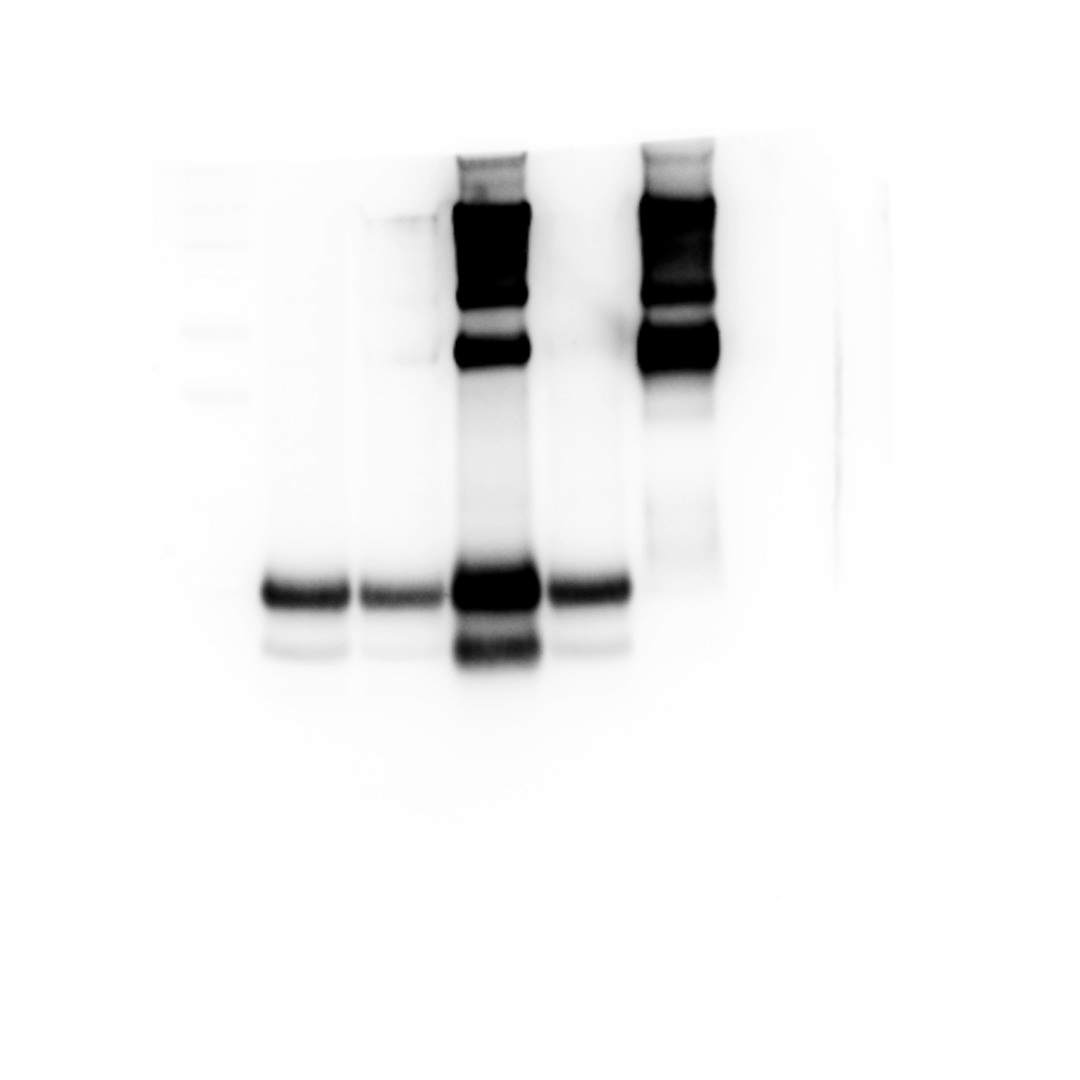

iii EMC4

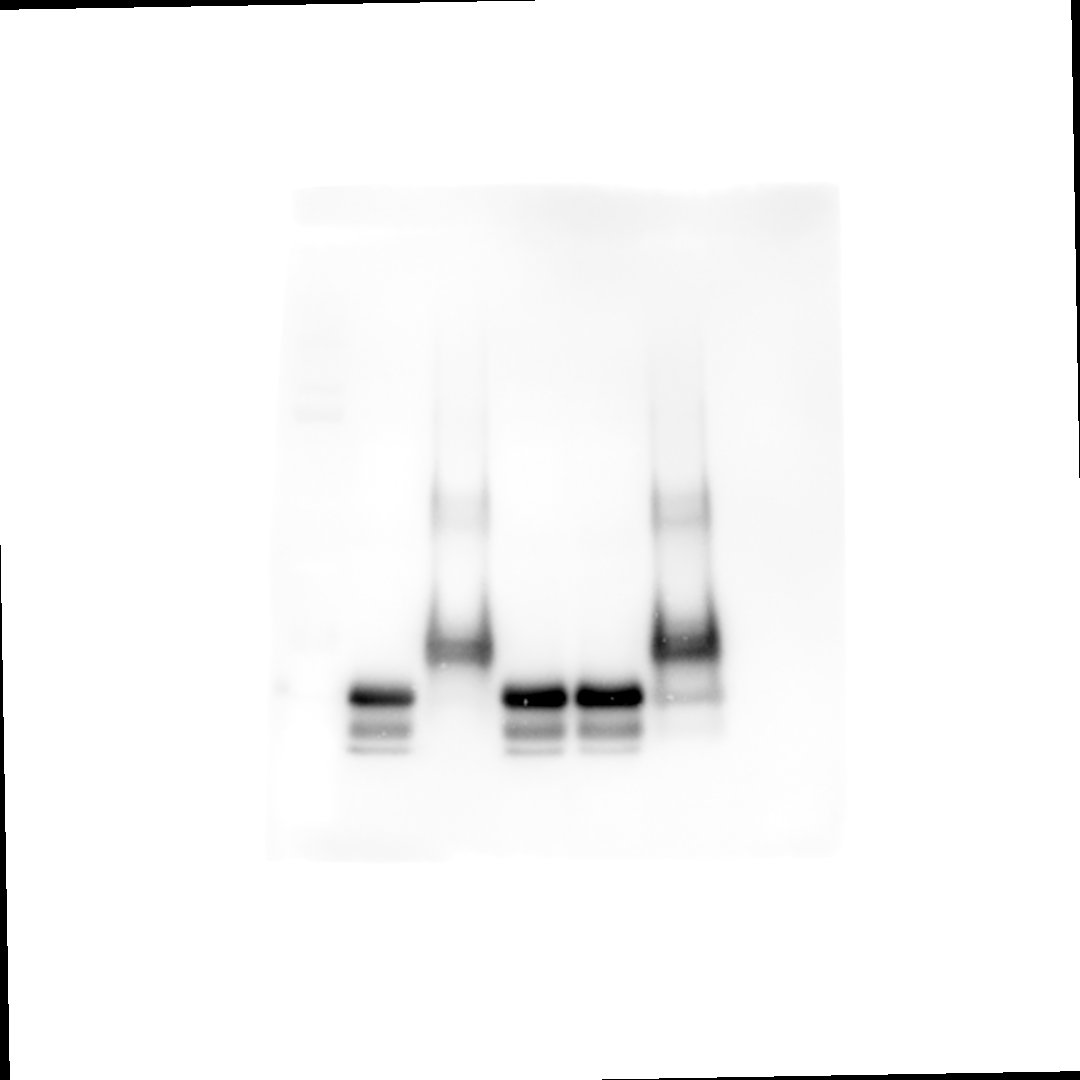

iv α-Synuclein

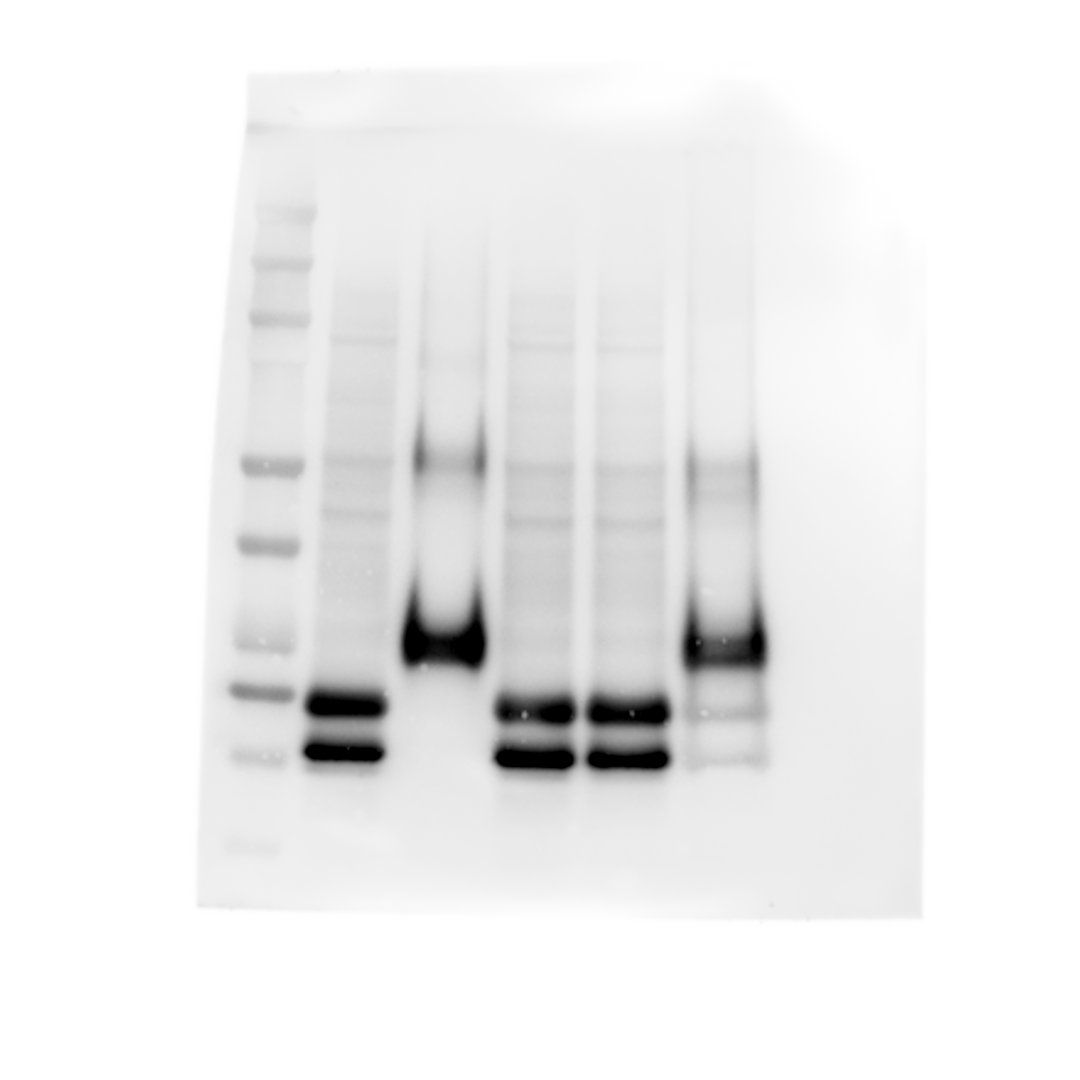

**Figure 6 e**

i LAMP1

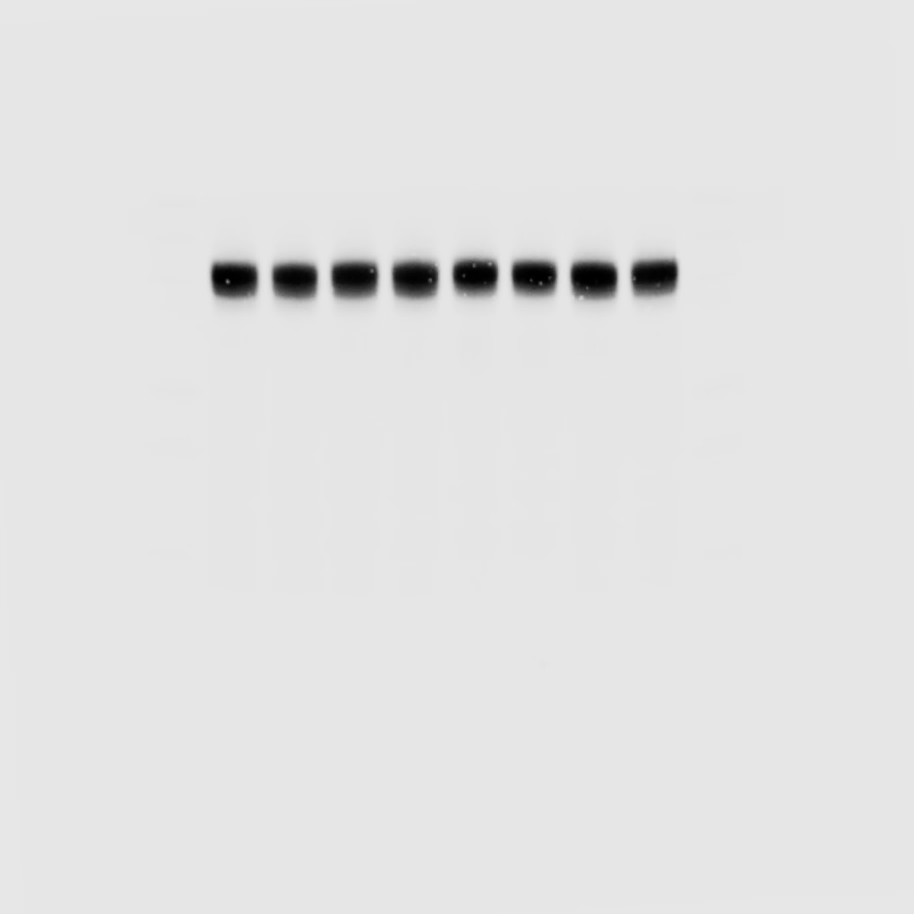

ii LC3B

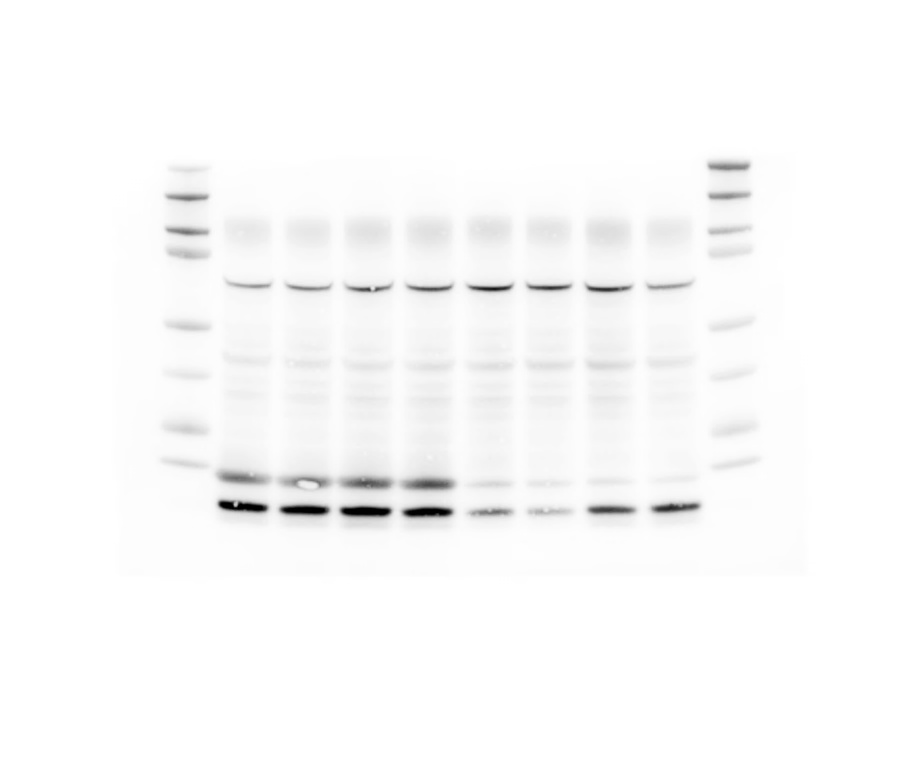

iii p62

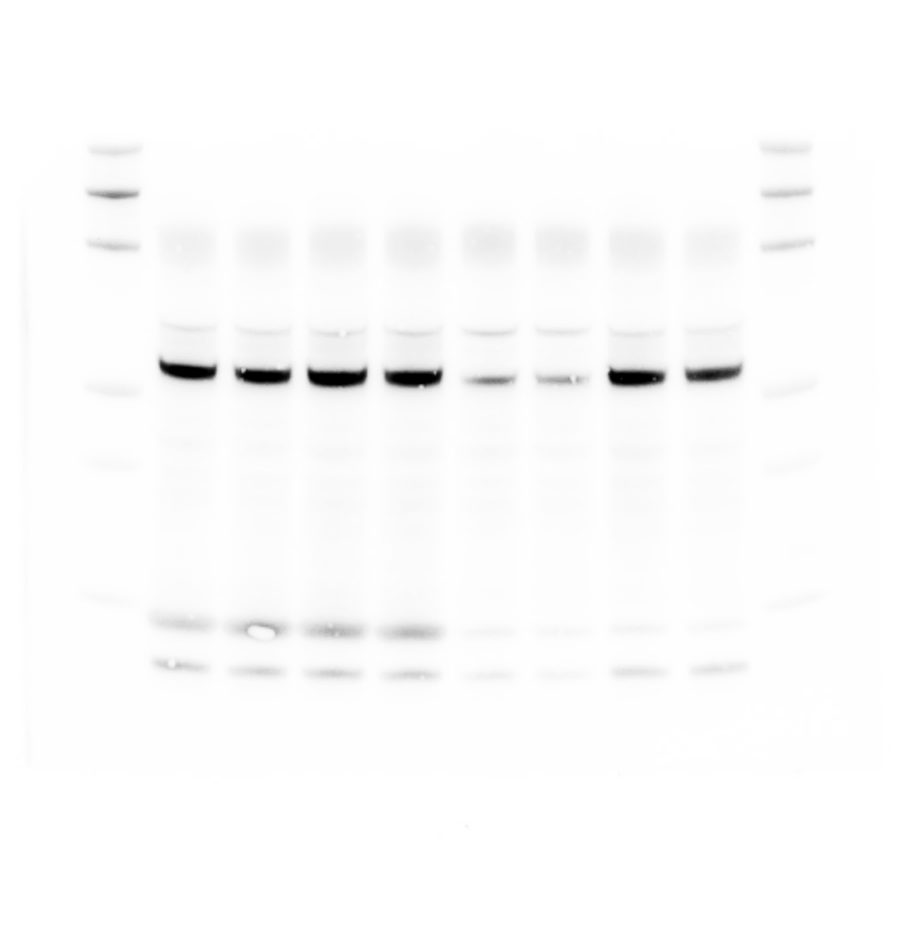

iv EMC4

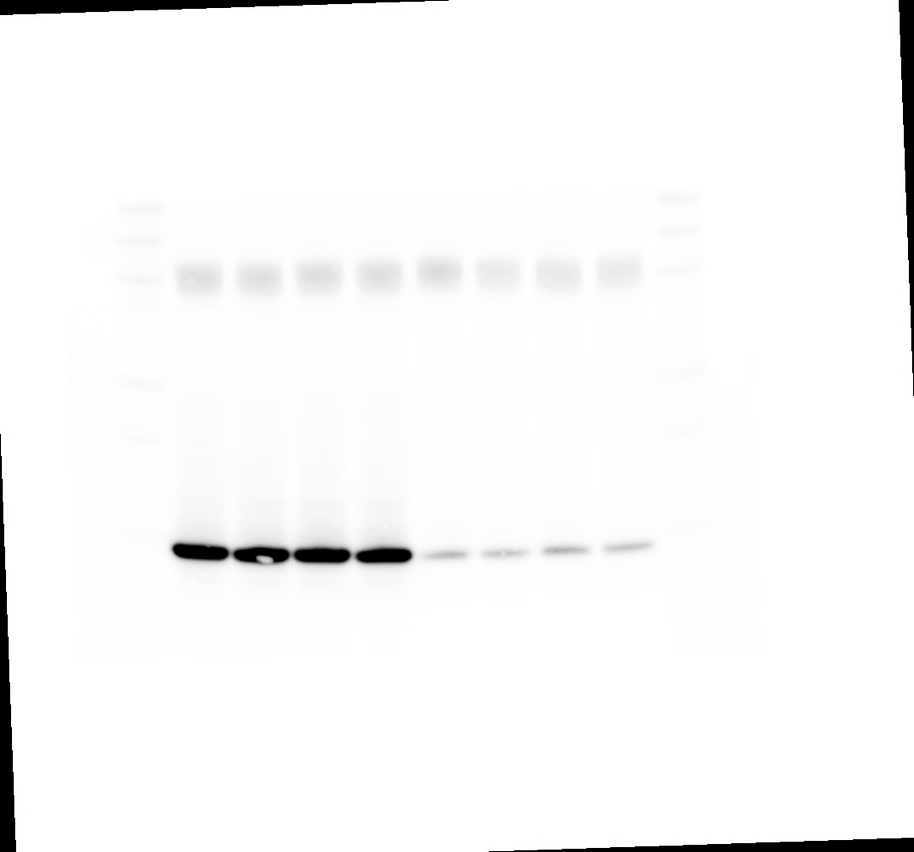

v B-Actin

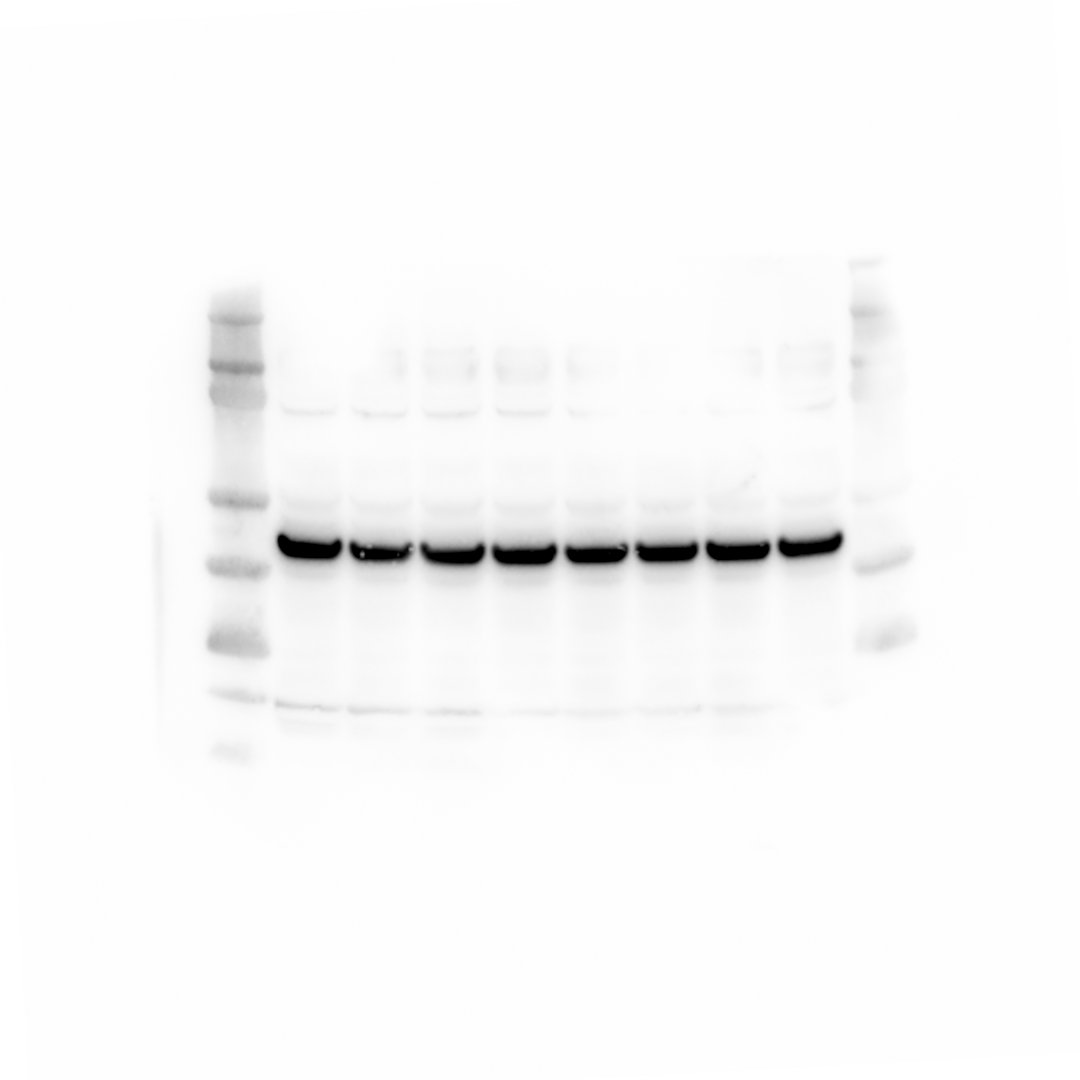

**Figure 7**

i EMC4

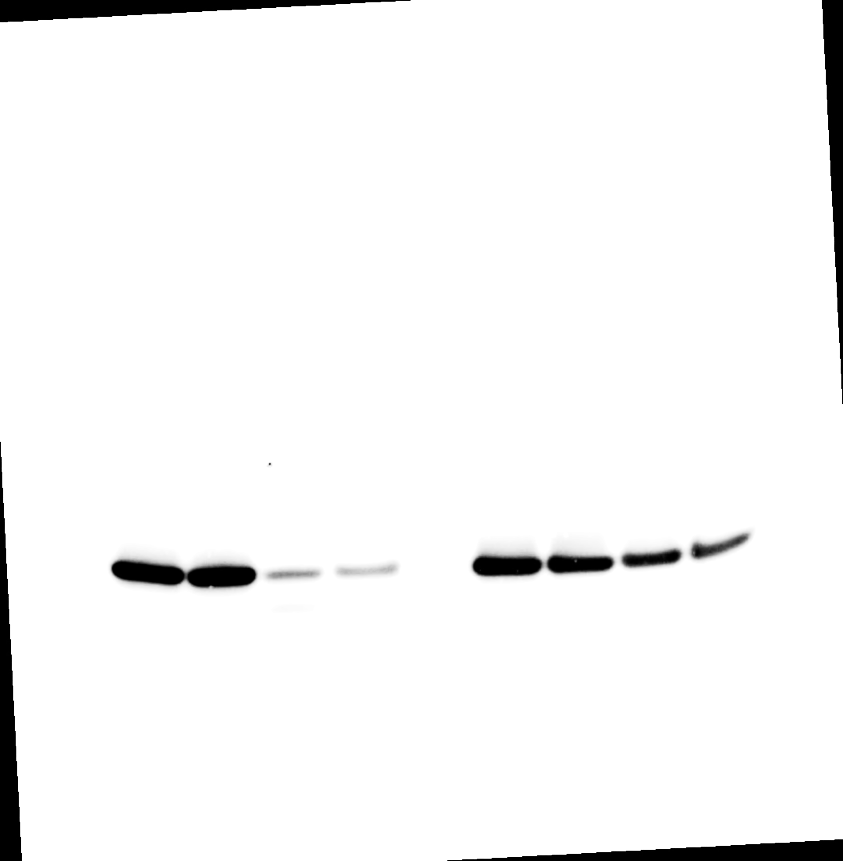

ii LC3B

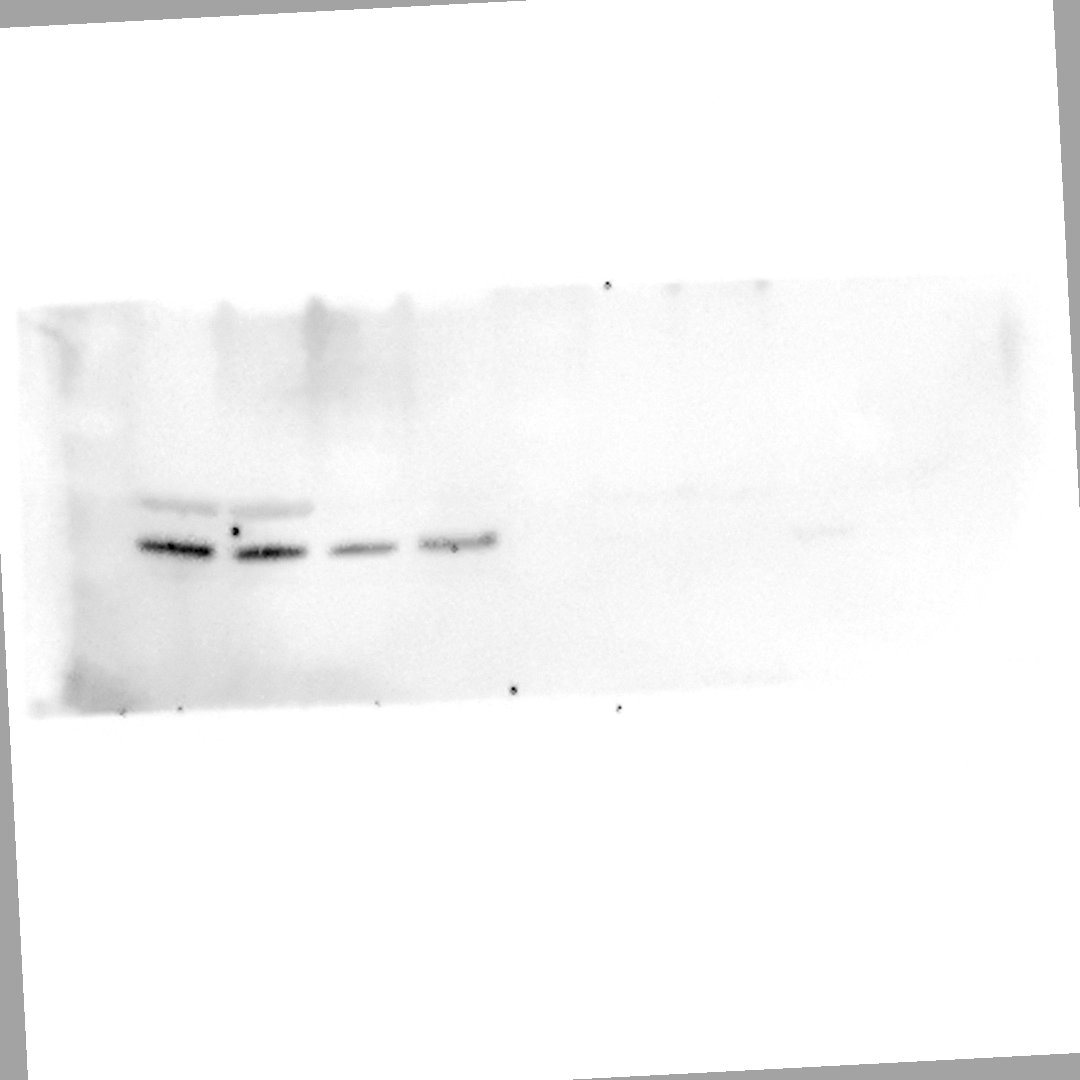

iii LAMP1

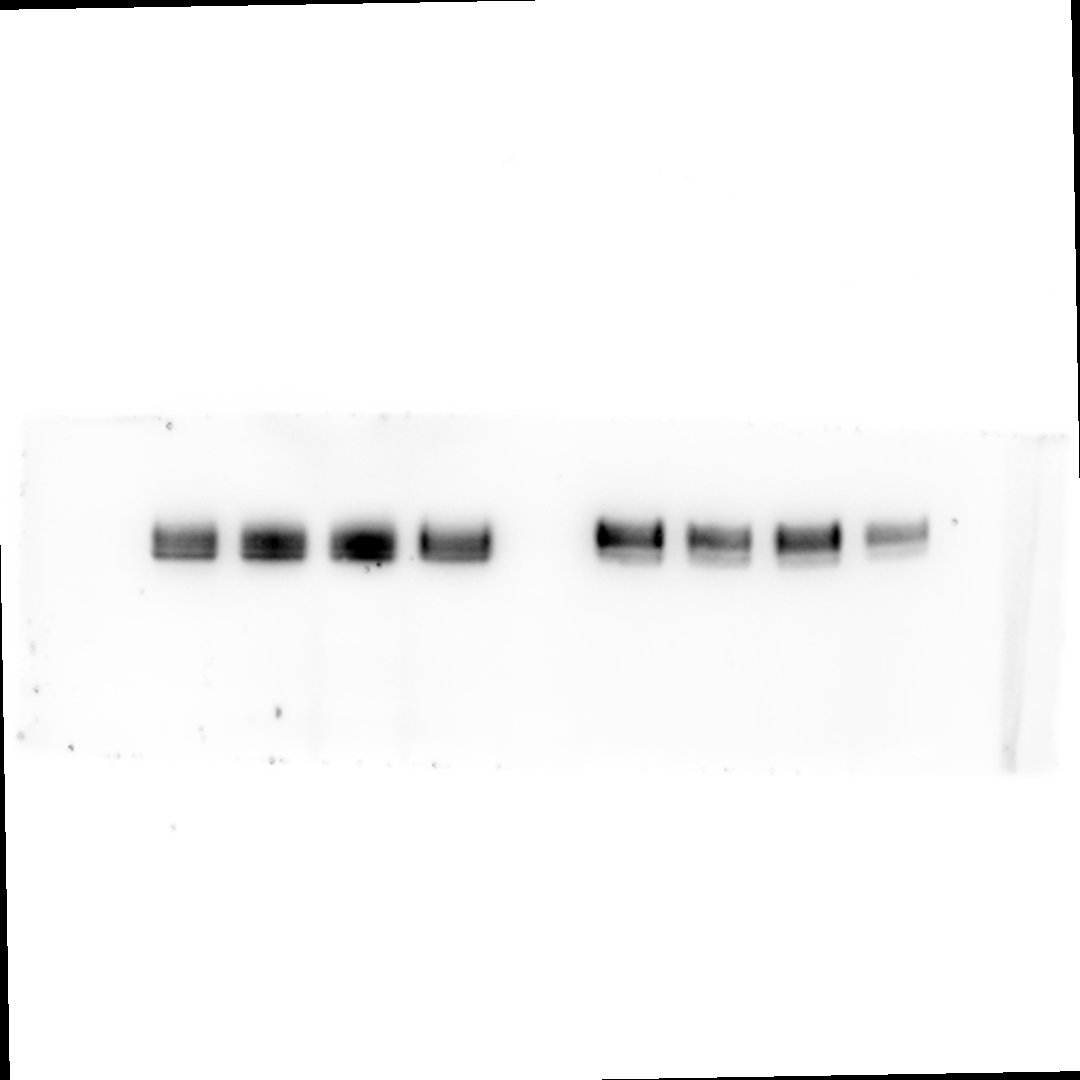

iv B-Actin

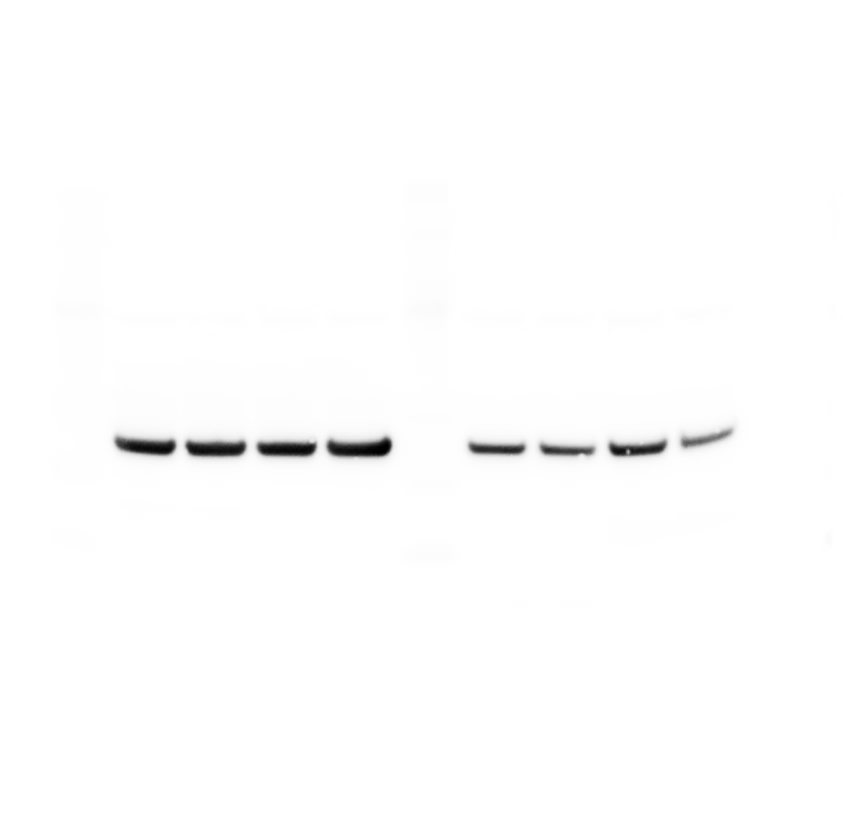

**Supplementary Figure 1**

**Supp. 1 c**

i) RAB13

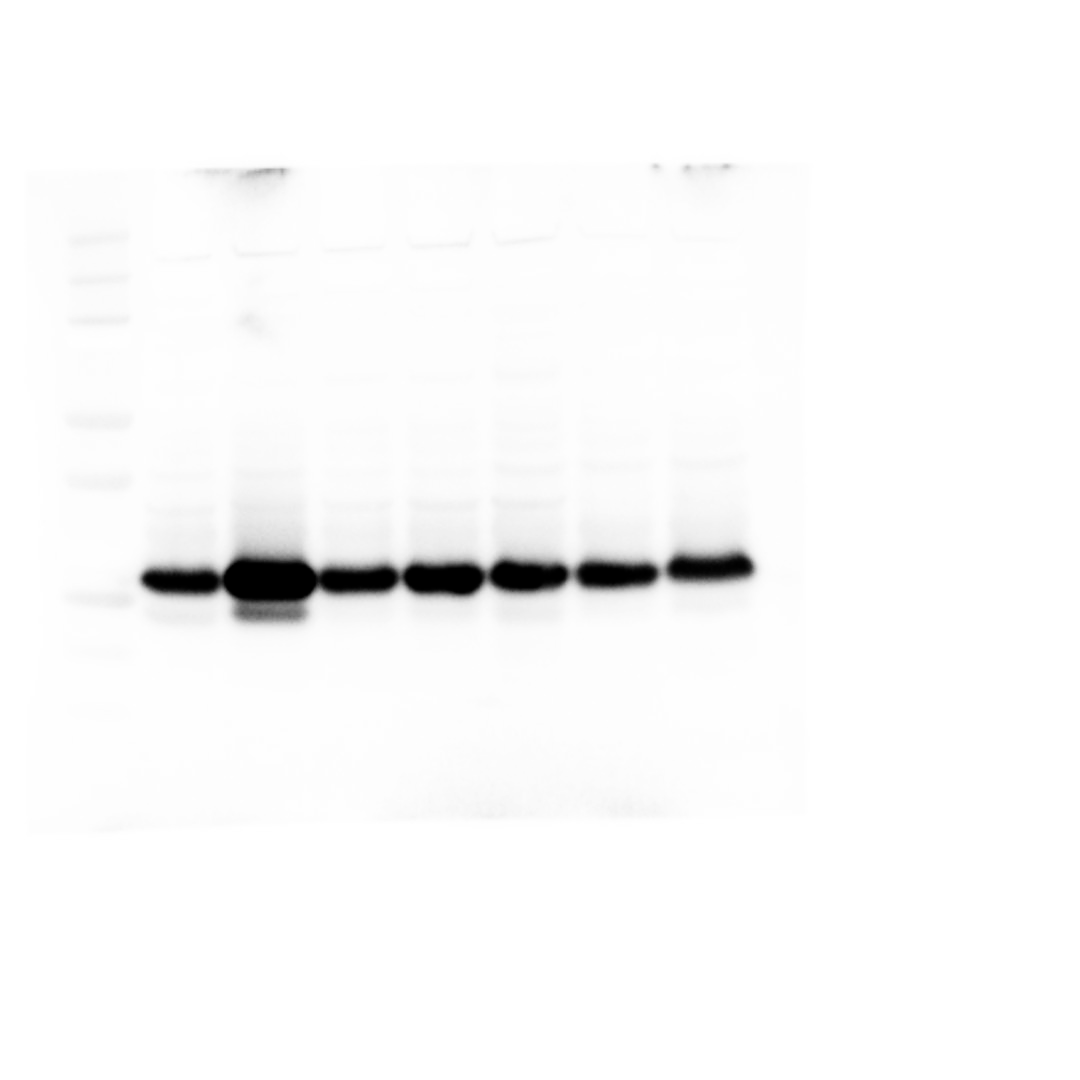

ii) B-Actin

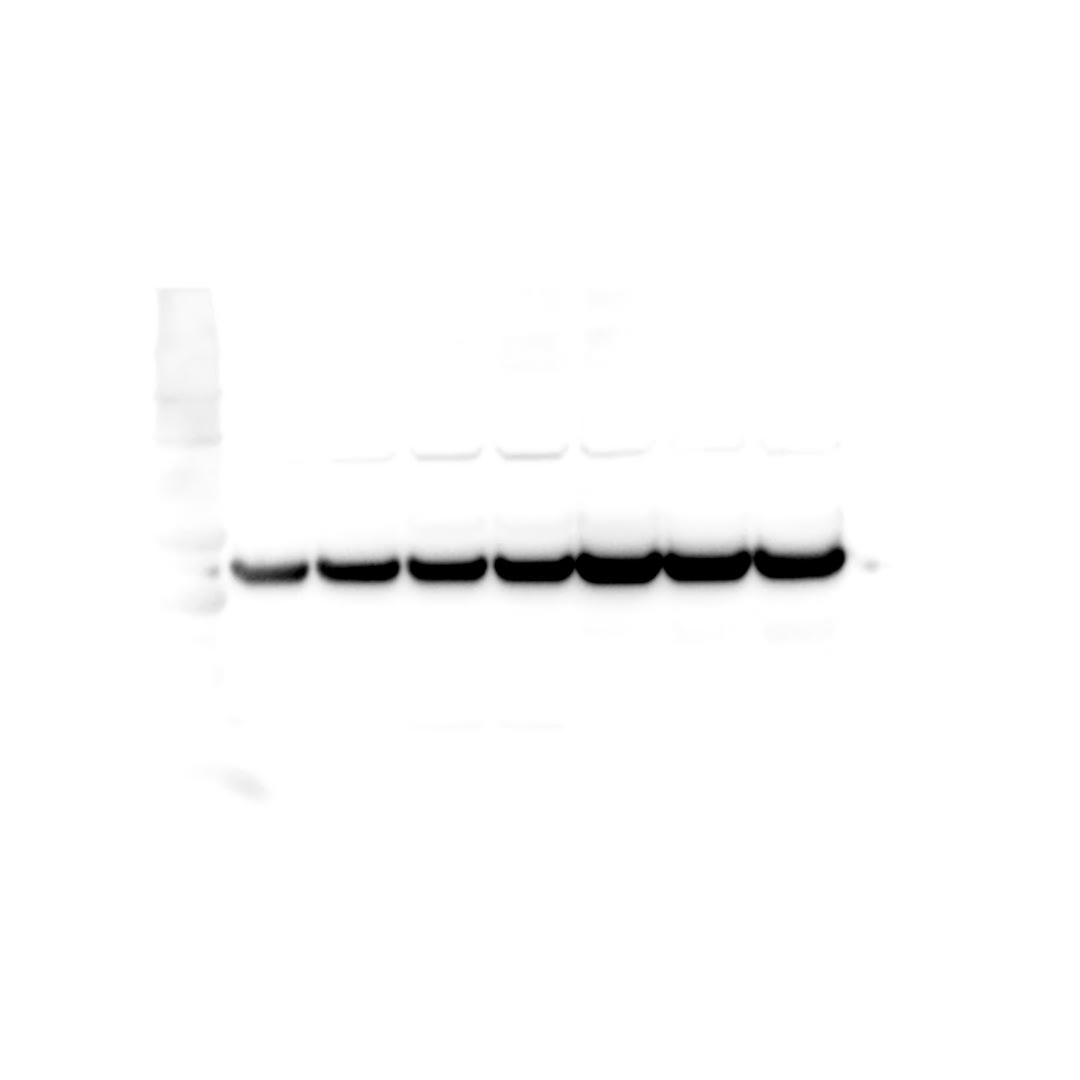

**Supp. 1e)**

i) TFAP2C

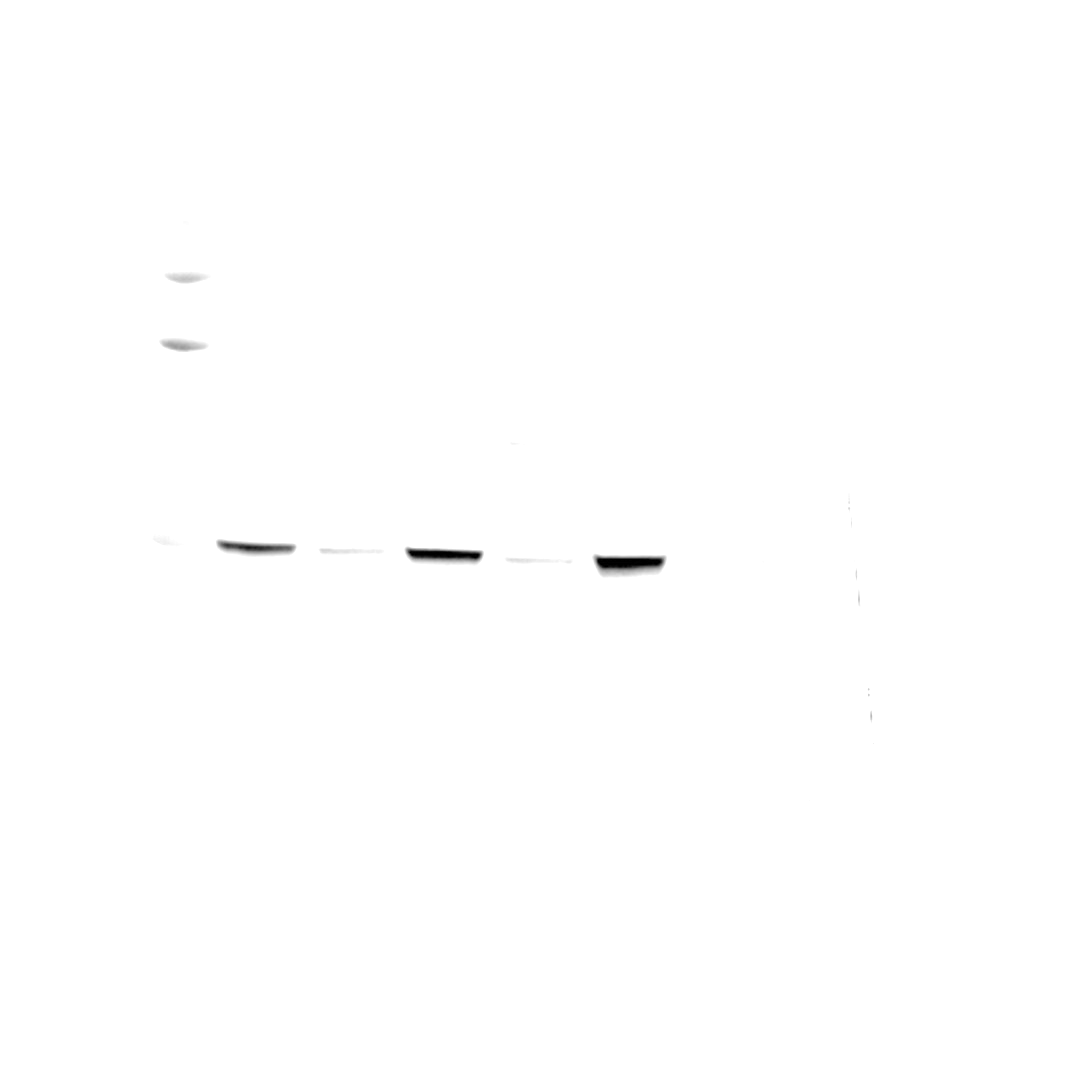

ii) B-Actin

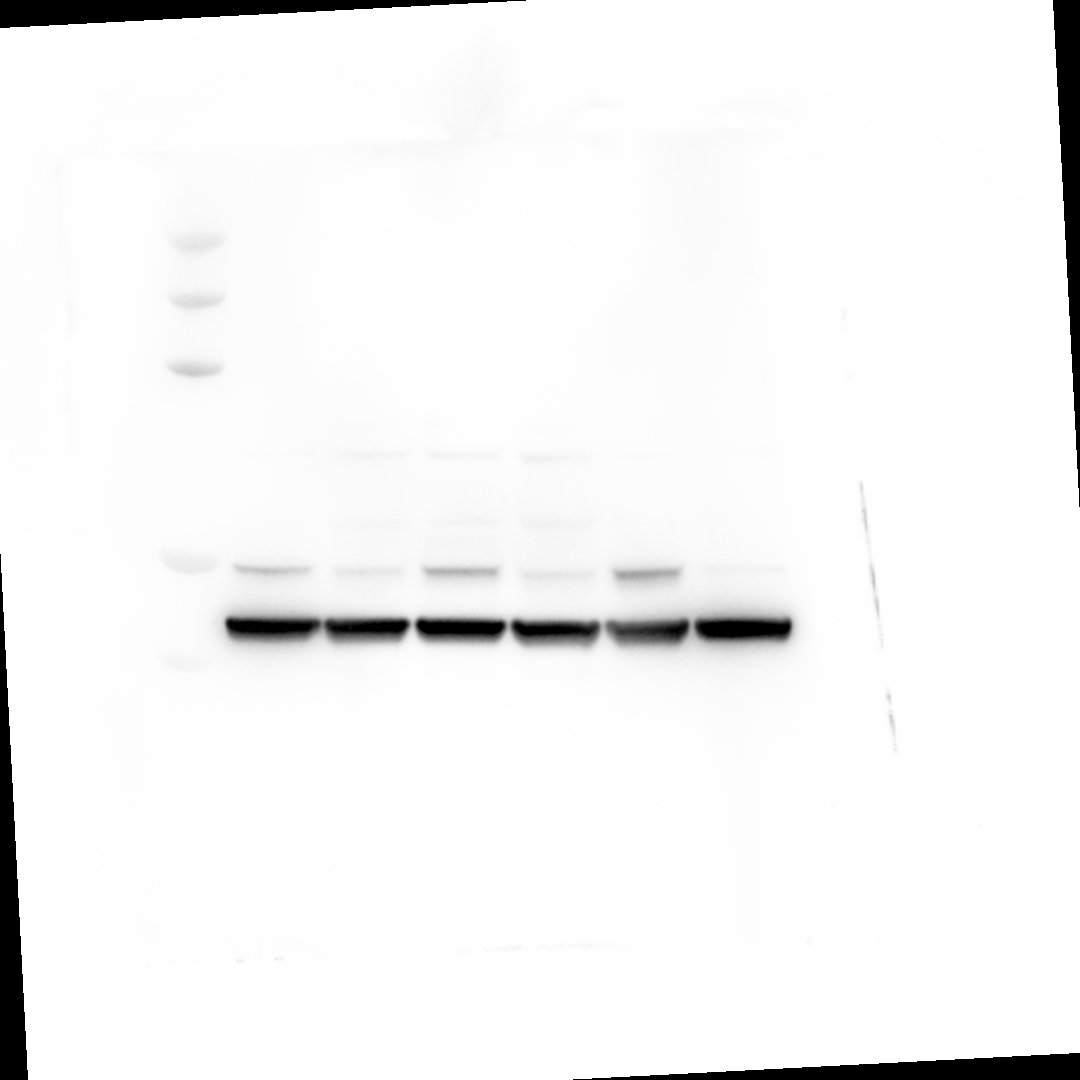

**Supp. 1f)** i) TFEB

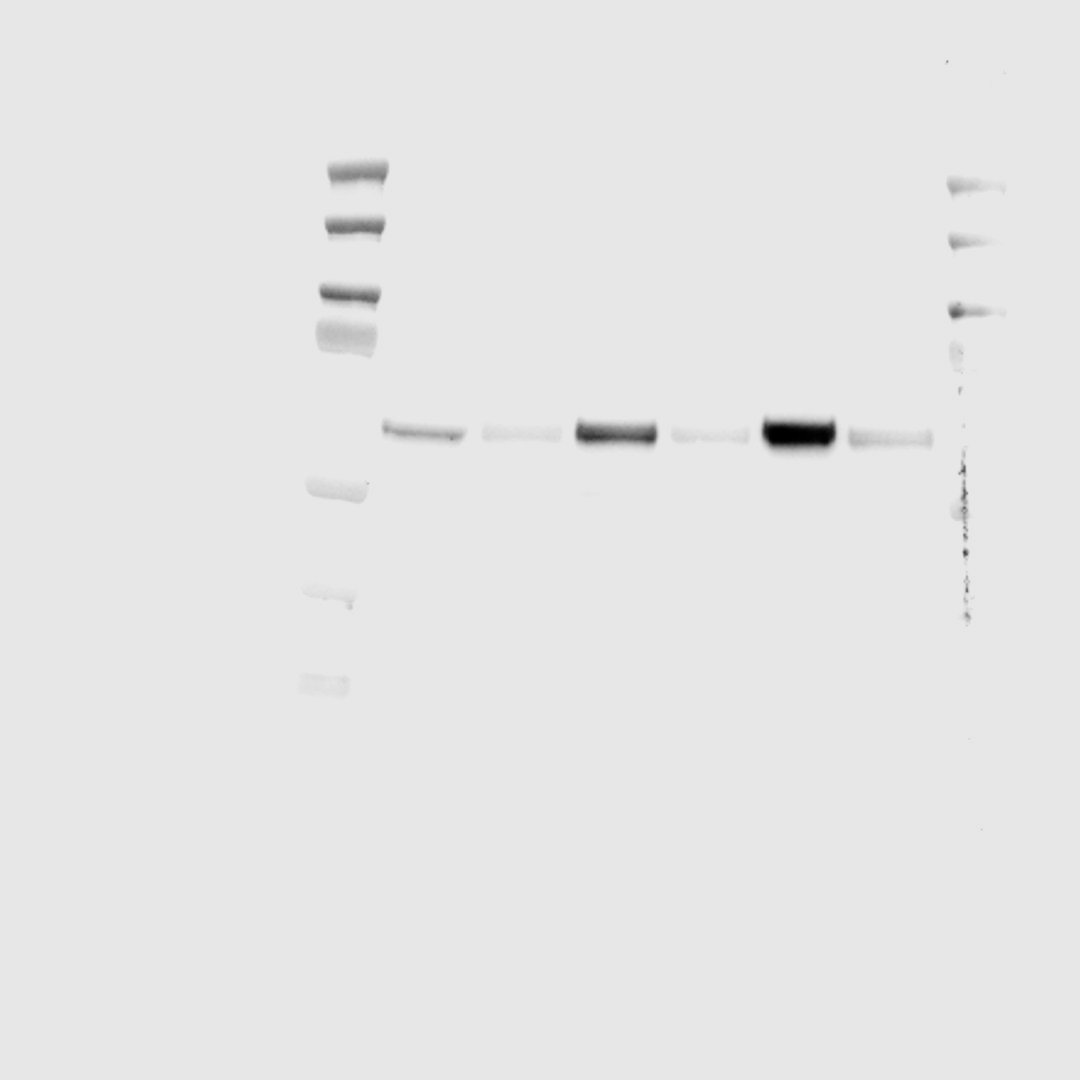

ii) B-Actin

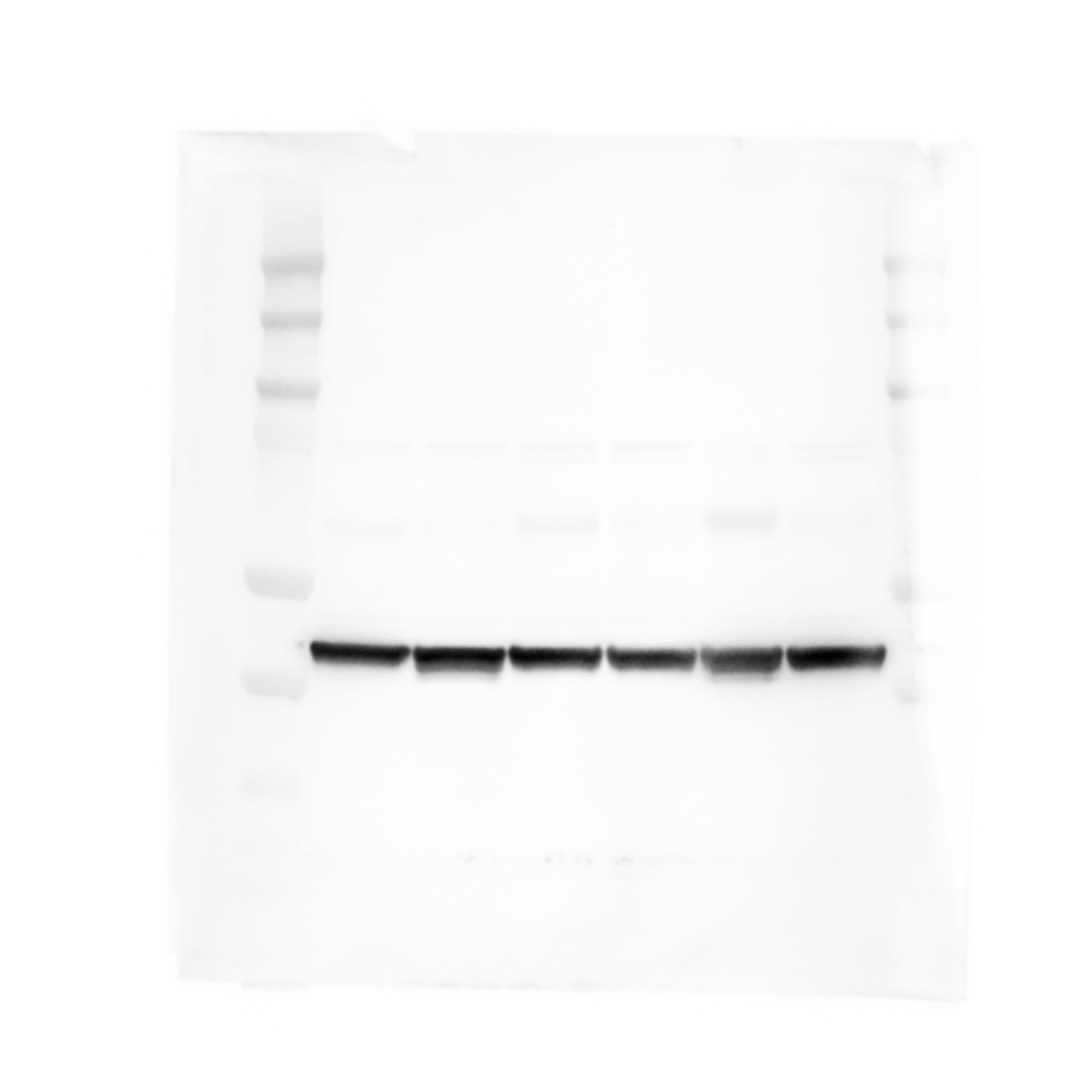

**Supplementary Figure 2**

**Supp. 2i**

i ) EMC4

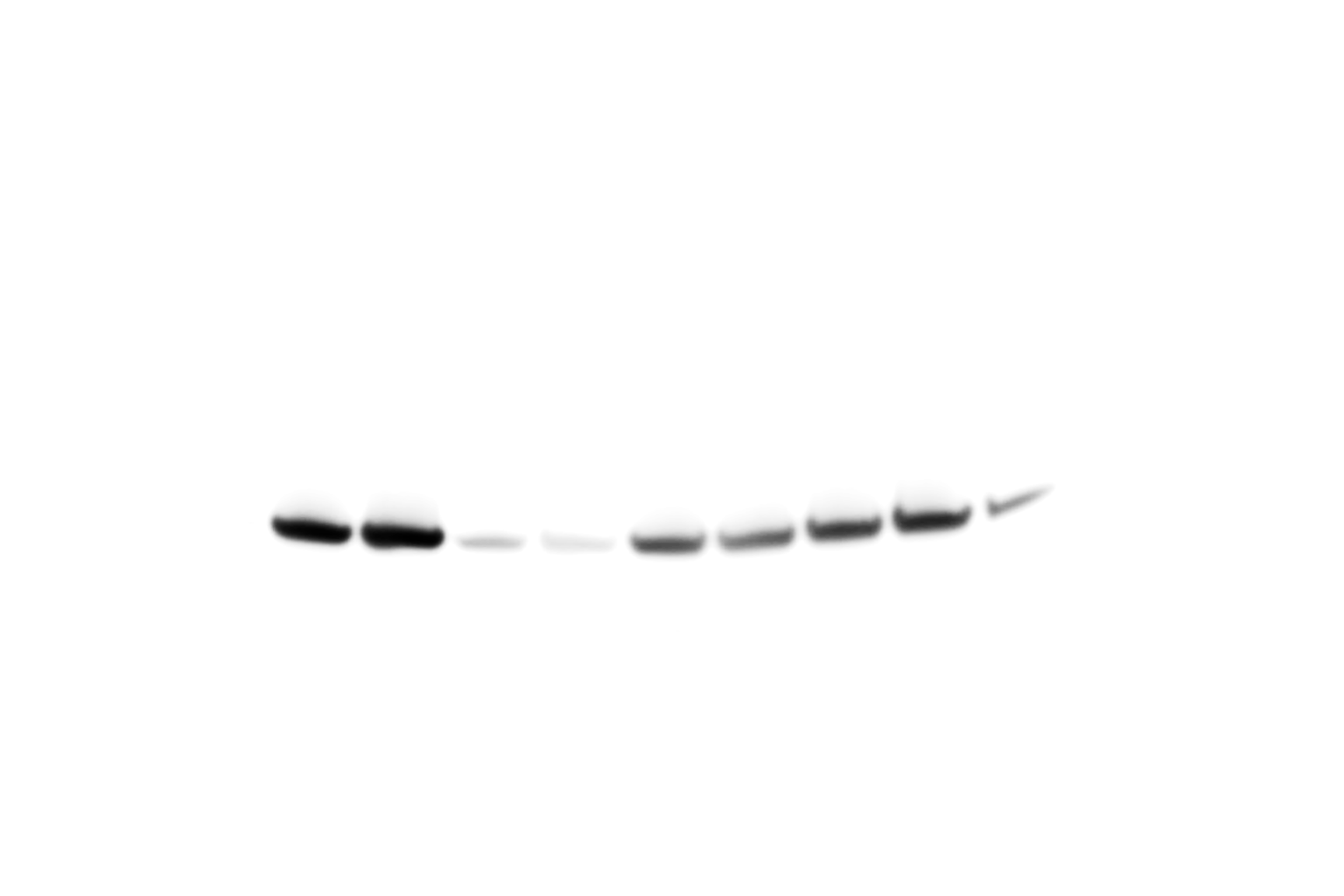

ii) B-Actin

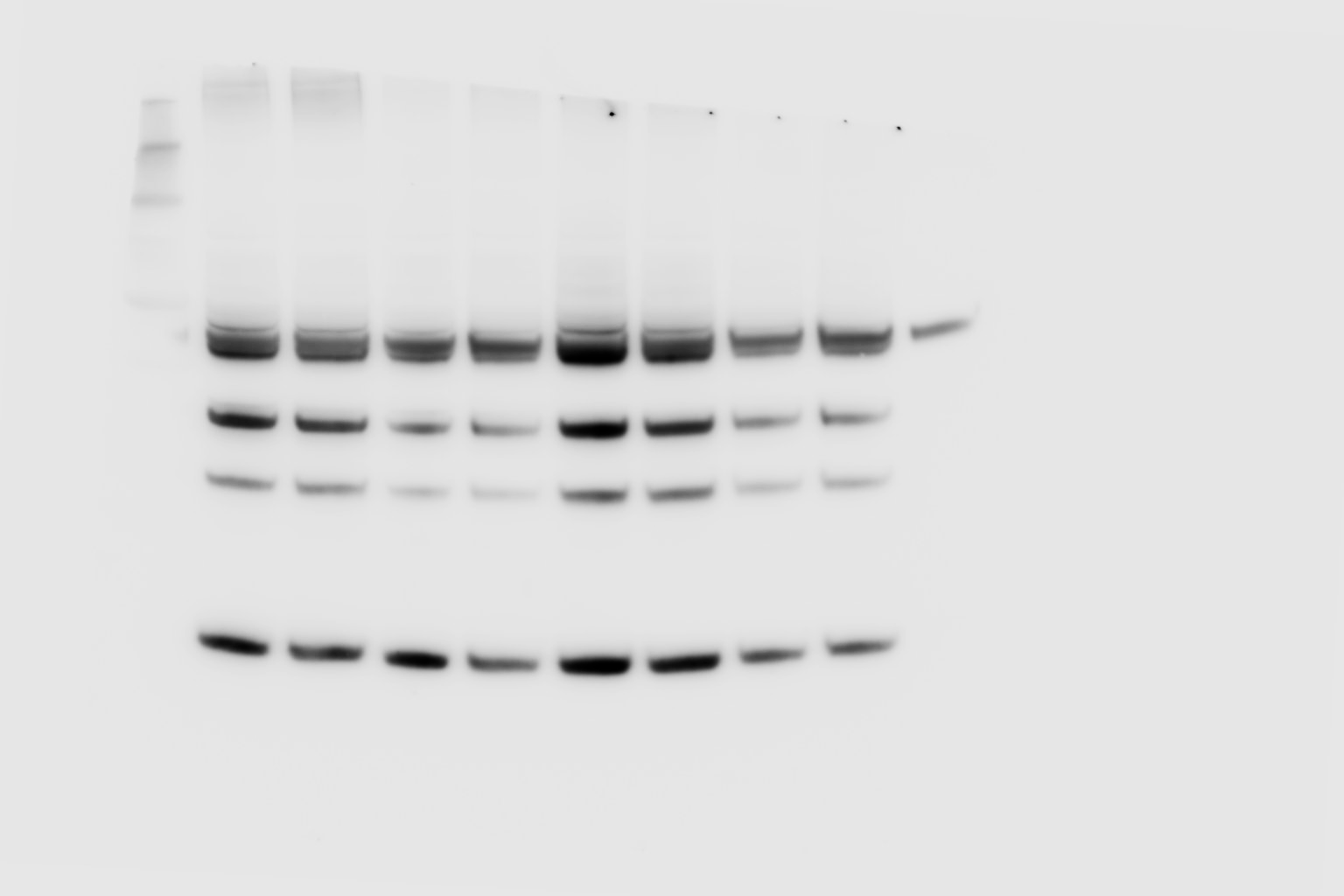

**Supp. 2k**
i) OXR1

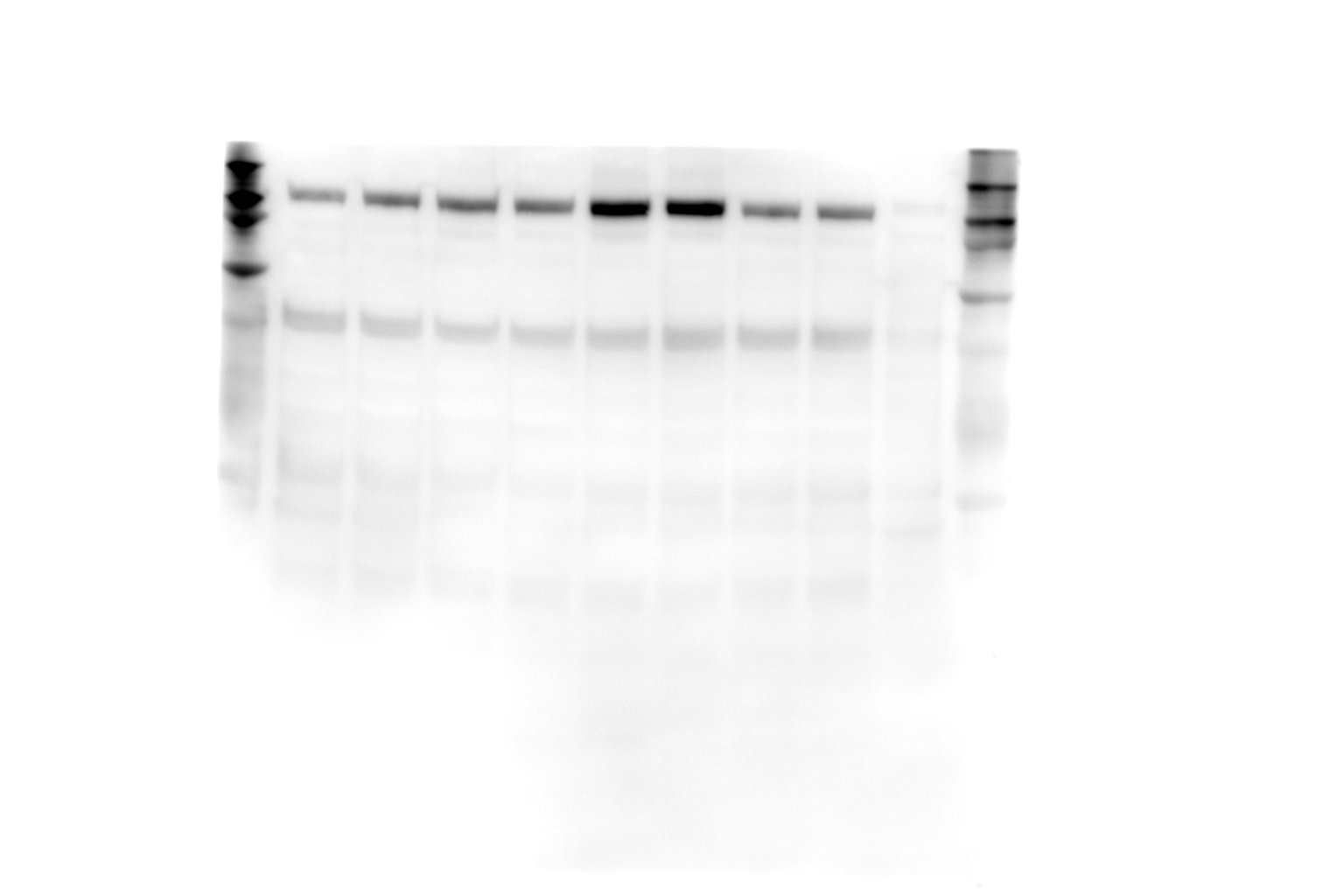

ii) B-Actin

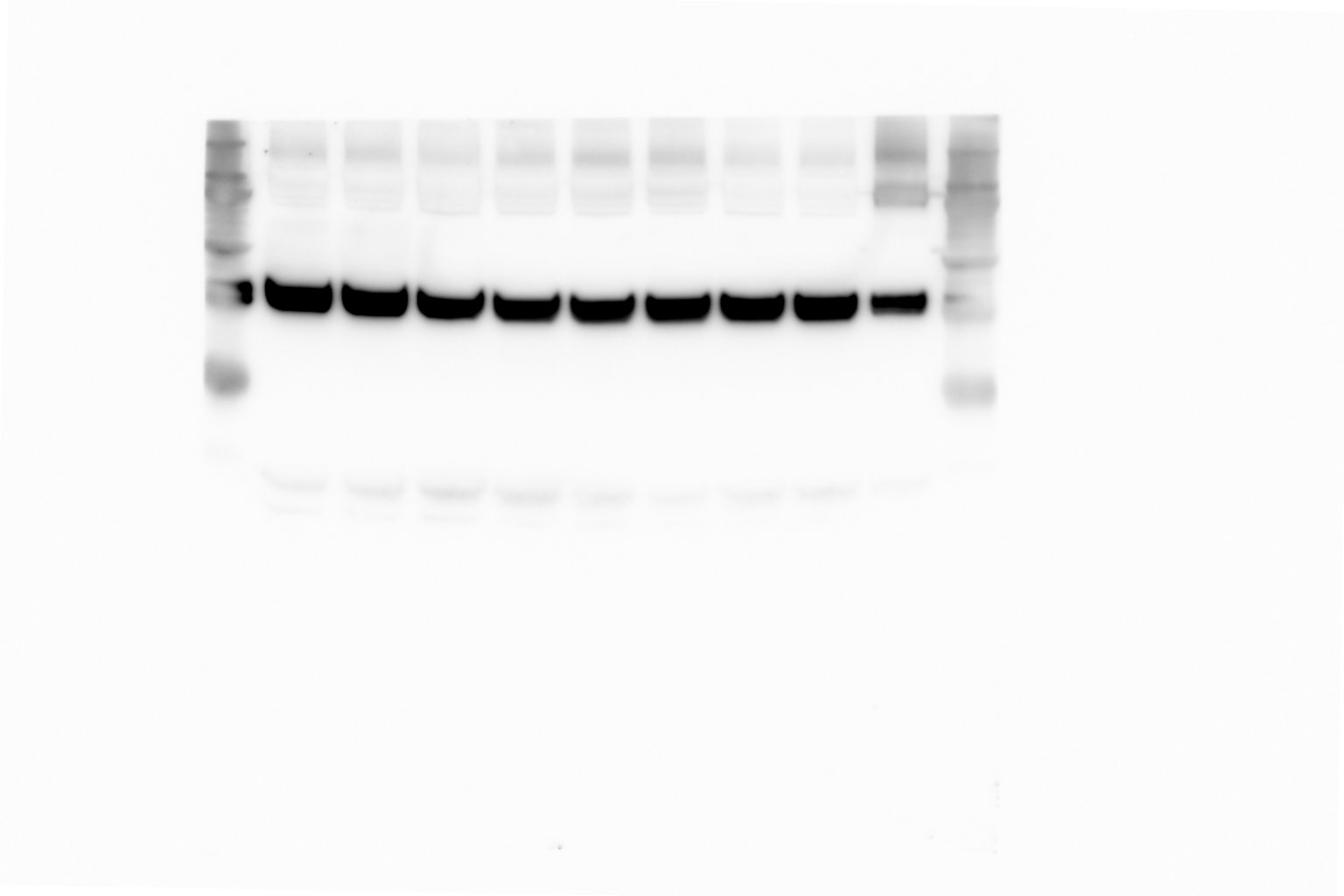

**Supplementary Figure 8**

**Supp. 8b**

i) α-Synuclein

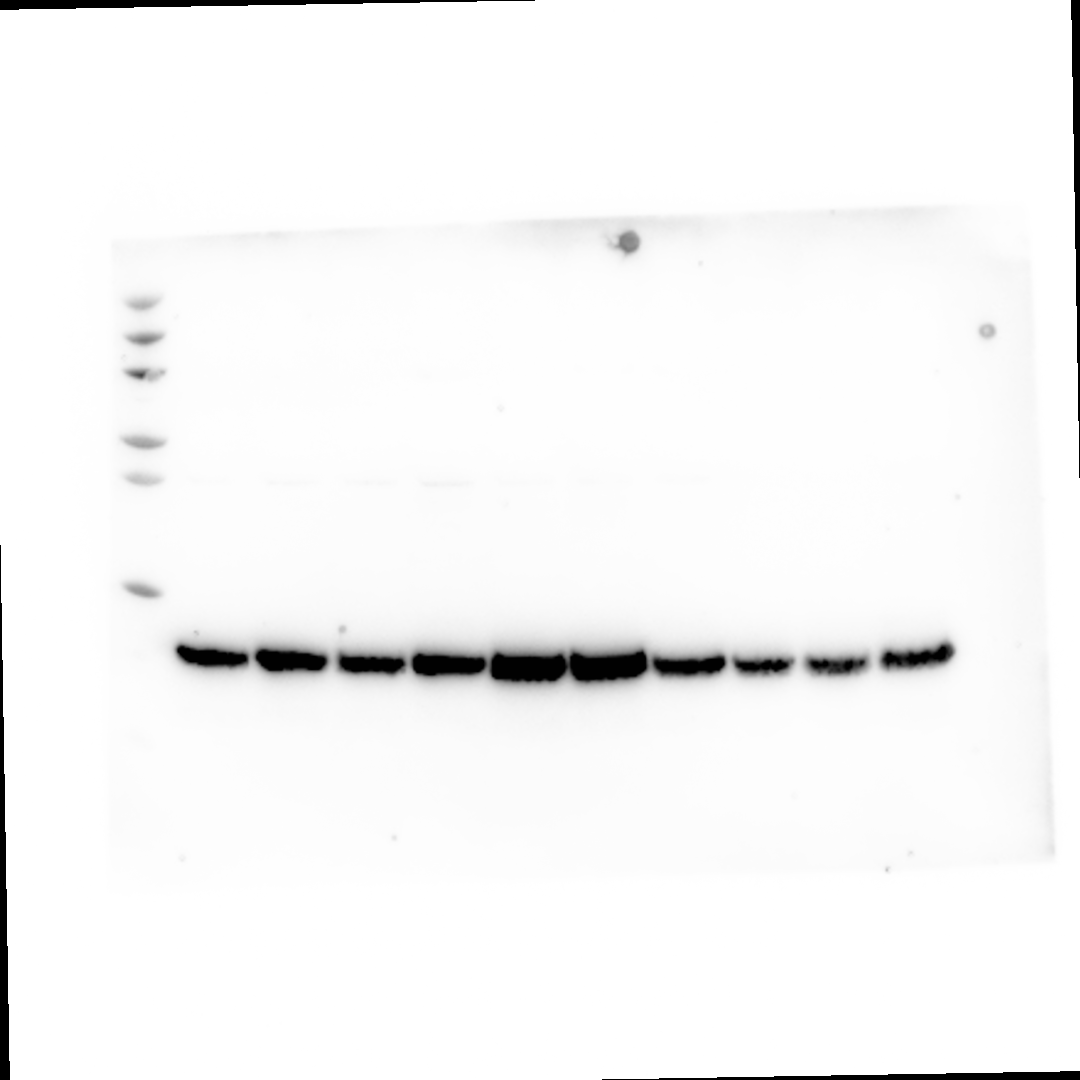

ii) OXR1

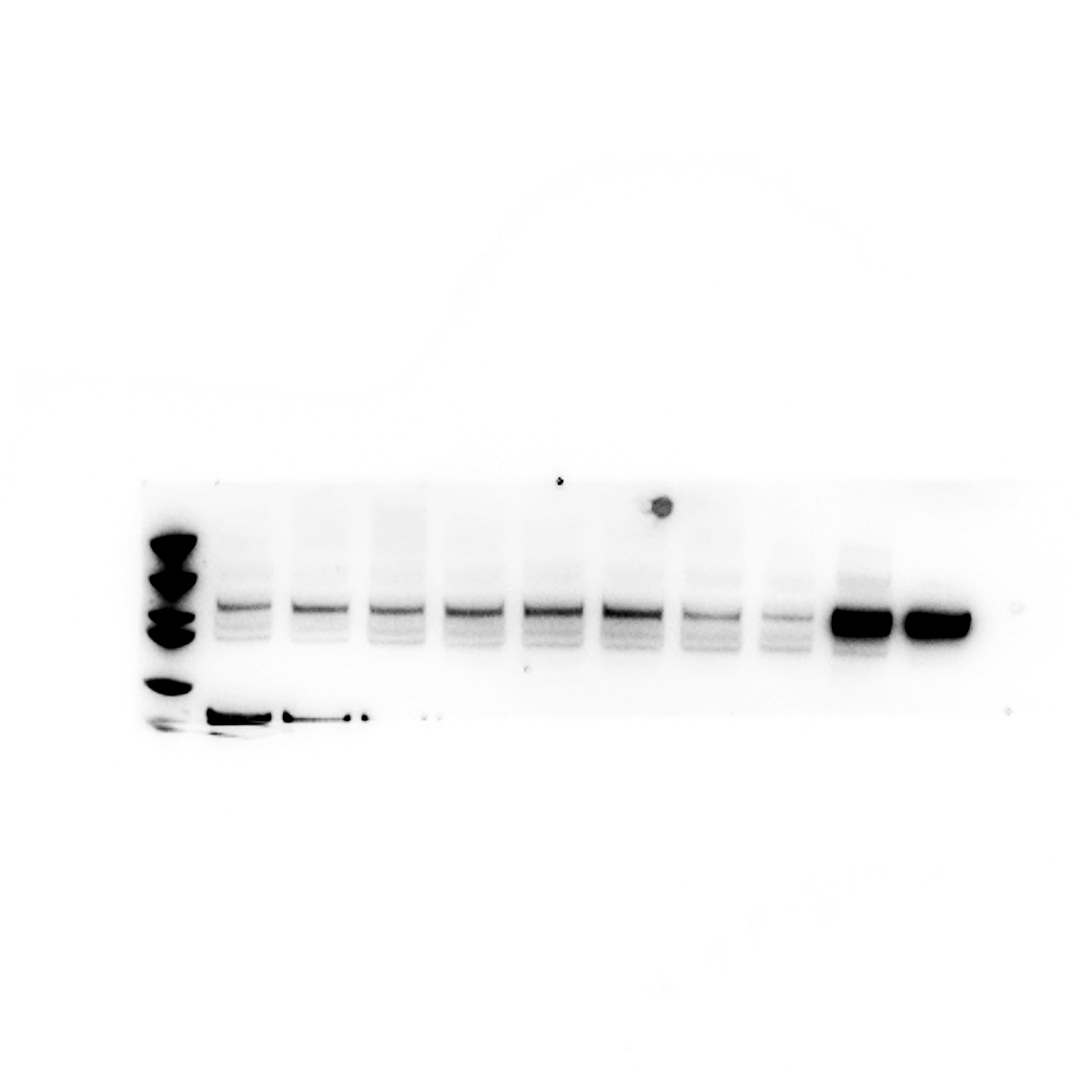

iii) B-Actin

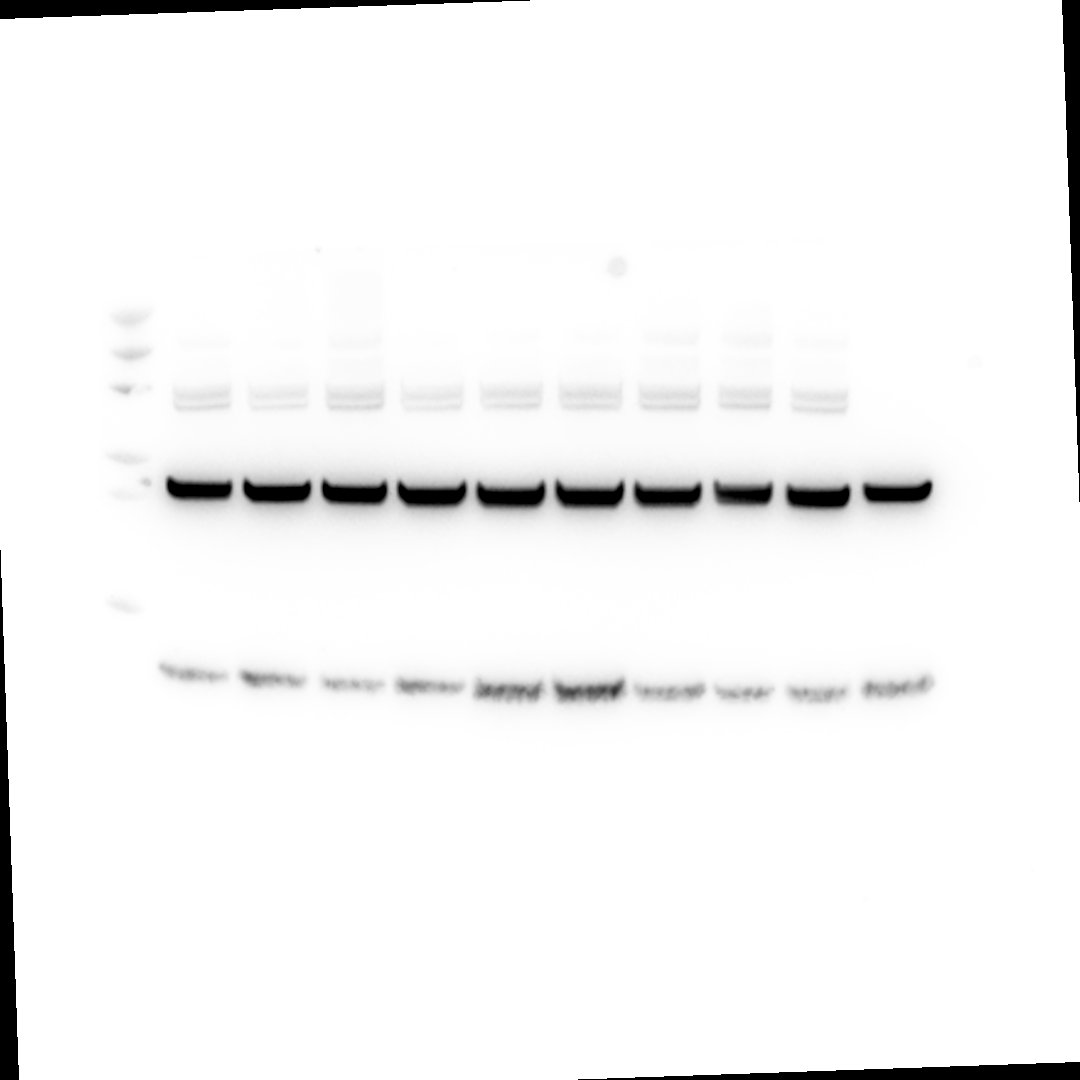
